# A Cross-kingdom Effector Modulates EDS1-dependent TIR-NLR-mediated Plant Immunity

**DOI:** 10.64898/2026.05.21.726970

**Authors:** Musharaf Hossain, Yangdou Wei, Peta C. Bonham-Smith, Christopher D. Todd

**Affiliations:** Department of Biology, University of Saskatchewan, Saskatoon, SK, S7N 5E2, Canada

**Keywords:** *Plasmodiophora brassicae*, Biotroph, Cross-kingdom, Effector, TIR-NLR, Effector-triggered immunity

## Abstract

In the ongoing battle between plants and pathogens, successful pathogens promote colonization by evolving new effectors that manipulate counteractive host responses, such as intracellular nucleotide-binding leucine-rich repeat (NLR)-triggered immunity. Here, we identify a secreted effector, PbSTMI, from the unique intracellular protist *Plasmodiophora brassicae*, that is conserved across divergent plant pathogens and broadly suppresses ENHANCED DISEASE SUSCEPTIBILITY 1 (EDS1)-dependent plant immunity. PbSTMI directly binds EDS1 and promotes its proteasome-mediated degradation, thereby diminishing downstream EDS1 oligomerization required for resistance. PbSTMI overexpression suppresses toll/interleukin-1 receptor (TIR) domain-containing NLR (TIR-NLR)-driven autoimmunity in *Arabidopsis* and further attenuates salicylic acid-mediated signaling in the autoimmune background. It blocks flg22-induced accumulation of EDS1 and regulates SA-mediated PAMP-triggered immunity (PTI) responses. Oligomerization among PbSTMI and its paralogs suggests coordinated immune suppression during different stages of *P. brassicae* infection and clubroot disease progression. PbSTMI suppression of EDS1-dependent TIR-NLR-mediated pathways increases susceptibility to a broad range of phytopathogens, including the biotrophs *P. brassicae* and *Erysiphe cichoracearum* and the hemibiotrophic *Colletotrichum higginsianum*. Together, these findings reveal a conserved pathogen strategy that disarms a central immune hub, tipping the balance of the plant-pathogen arms race decisively in favor of the pathogen.

## 1. Introduction

*Plasmodiophora brassicae*, a unique obligate biotrophic protist that causes clubroot disease with characteristics of root gall formation, can evade plant immune responses upon invading root tissues (1,2). Throughout its intracellular lifestyle, it secretes a large number of effector proteins into host cells that manipulate host defense signaling and intracellular trafficking and alter host hormone pathways to promote *P. brassicae* colonization of root tissues (1–3). During natural infection, the co-evolutionary arms race between plants and pathogens determines whether colonization succeeds or robust host resistance emerges. Resistance (R) genes linked to quantitative trait loci that confer resistance to *P. brassicae* in canola and other Brassica species have been identified (4–7). Equally, the rapid loss of resistance in previously clubroot-resistant varieties may be linked to the array of secreted effectors from virulent, rapidly evolving *P. brassicae* pathotypes (8), underscoring the need for more comprehensive research to understand *P. brassicae* pathogenesis. However, with its obligate biotrophic nature, intracellular colonization, and rapid diversification into distinct pathotypes, together with no current methods for genetic manipulation, detailed mechanistic research of *P. brassicae* is limited in scope.

Plants have a sophisticated immune surveillance system that enables them to detect and respond to pathogen infection for resistance. Plant immunity primarily relies on two key elements: pattern-triggered immunity (PTI) and effector-triggered immunity (ETI) (9). PTI is initiated when pattern recognition receptors (PRRs) at the cell membrane recognize conserved pathogen-associated molecular patterns (PAMPs). Conversely, ETI is initiated when pathogen effectors compromise PTI, necessitating a second line of defense for plant resistance (9). Resistance (R) proteins mediate ETI, the largest group of which are intracellular nucleotide-binding leucine-rich repeat (NLR) proteins that, directly or indirectly, detect specific pathogen Avr effectors, leading to robust, durable immune responses often associated with the hypersensitive response (HR) (10,11).

NLRs are highly conserved between animals and plants and are broadly categorized into two main types based on their N-terminal domains: toll/interleukin-1 receptor (TIR) domain-containing NLRs (TIR-NLRs) and coiled-coil (CC) domain-containing NLRs (CC-NLRs) (12–14), which are involved in oligomerization, downstream immune signaling, and the execution of cell death (15,16). NLRs also contain the ATP/ADP-binding molecular switch (NB-ARC), shared by APAF-1, R proteins, and CED-4 (11,17), and a C-terminal leucine-rich repeat (LRR) domain that facilitates pathogen effector recognition and binding (18,19). TIR-NLRs and CC-NLRs have complementary roles in plant immunity, and their interactions highlight the specificity and complexity of NLR-mediated immune responses (12).

Upon recognizing a pathogen effector, sensor TIR-NLRs signal a cellular immune response by triggering heterodimerization of the lipase-like protein ENHANCED DISEASE SUSCEPTIBILITY 1 (EDS1) with PHYTOALEXIN DEFICIENT 4 (PAD4) or EDS1 with SENESCENCE-ASSOCIATED GENE 101 (SAG101), leading to activation of distinct defense responses (20,21). Additionally, a less abundant subclass of CC-NLRs, the CC-RPW8-domain (possessing an RPW8-like CC domain - CCR) containing helper NLRs (also known as RNL) (22), including ACCELERATED DISEASE RESISTANCE 1 (ADR1, ADR1-L1 & ADR1-L2 in Arabidopsis; (20,23) and N REQUIRED GENE 1 (NRG1; (20,24,25), are involved in mediating immune responses initiated by TIR-NLR-sensor NLRs (14,26,27). The SUPPRESSOR OF NON-EXPRESSOR OF PATHOGENESIS-RELATED GENES 1, CONSTITUTIVE 1 (SNC1), is a sensor NLR that associates with the EDS1-PAD4-ADR1 module and triggers immune responses while guarding the CCR domain of ADR1-L1/L2 (28,29). Similarly, the sensor NLR, CHILLING SENSITIVE 3 (CHS), together with its genetically linked partner, the executer NLR, CONSTITUTIVE SHADE-AVOIDANCE 1 (CSA1), transduces a pathogen effector signal via the EDS1-SAG101-NRG1 module, triggering downstream immune responses, including cytoplasmic calcium ion (Ca^2+^) influx and subsequent cell death (12,28,30). Gain-of-function (GOF) mutants of both ADR1 and NRG1 show severe dwarfism and autoimmunity due to increased cytoplasmic Ca^2+^ influx through plasmamembrane enrichment of oligomerized mutant Ca^2+^ channels (31). Salicylic acid (SA)-mediated responses to pathogen attack are intricately linked with NLR-mediated immunity. Accumulation of SA enhances expression of NLR genes and other defense-related genes, thereby amplifying and coordinating a robust defense response (14,32). In addition to being vital regulators of the TIR-NLR-mediated immunity pathway, EDS1 and PAD4 are also central to SA signaling (33–35). EDS1-NPR1 interactions induce pathogenesis-related (PR) gene expression crucial for both SA-dependent and NLR-mediated immunity (36). Recent studies have also identified a convergence of PTI and TIR-NLR-mediated effector-triggered immunity (ETI) in Arabidopsis, underscoring the role of EDS1 and its coordination with LRR-containing cell-surface receptors in enhancing plant resistance (37,38). However, understanding the diverse mechanisms by which this robust, multilayered resistance is constrained by the rapid evolution of plant pathogens demands more comprehensive research. Several studies show that diverse pathogen and pest effectors can undermine plant immunity by targeting central TIR-NLR signaling components such as EDS1, PAD4, and ADR1, either by sequestration into cellular compartments, such as processing bodies (39) or by disrupting the essential interactions required for effective immune activation (40,41).

In this study, we have identified a *P. brassicae* effector, PbSTMI, that suppresses EDS1-dependent plant NLR immunity and is conserved across diverse plant pathogens. We show that PbSTMI attenuates autoimmunity mediated by the GOF TIR-NLR mutants, *snc1* and *chs3-1*, by interacting with and degrading AtEDS1. In addition to alleviating NLR immunity, PbSTMI-mediated EDS1 turnover suppresses SA biosynthesis and impedes SA-dependent immune signaling in the *snc1* background, thereby impairing plant resistance against a broad range of phytopathogens. Furthermore, PbSTMI inhibits flg22-triggered, SA-mediated late PTI responses, thereby disrupting the coordinated efforts of plant resistance. Homo- and hetero-oligomerization among PbSTMI paralogs with distinct expression patterns suggests a possible spatial and temporal regulation of plant immunity during clubroot infection. Our findings highlight the impact of pathogen effectors on NLR-mediated immunity and host defenses, underscoring the need for a comprehensive approach to plant growth and the establishment of durable resistance.

## 2. Results

### 2.1. PbSTMI is a *Plasmodiophora brassicae* effector with cross-kingdom conservation among plant pathogens

RNA sequencing and cDNA library screening of tissues infected with *P. brassicae* (pathotype 3) have identified several effectors, including PbSTMI, which are particularly abundant during secondary infection (1,2). A BlastP search against the NCBI non-redundant database identified orthologs to PbSTMI in the Ascomycetes and Basidiomycetes, within the super kingdom Eumycota (true fungi) (**Figure 1A, Table S1**). A phylogenetic tree of PbSTMI and its orthologs shows separate biotrophic and hemibiotrophic clusters, indicating conservation of the effector family across pathogens with different infection strategies (**Figure 1A**). The patchy distribution of this effector across the protist *P. brassicae*, and orthologs in certain ascomycetes such as *Colletotrichum* and *Pyricularia*, as well as basidiomycetes like gall-inducing smut fungi, along with its absence from other related taxa, suggests a complex evolutionary history of lineage-specific gene loss and deviations from strict vertical inheritance, possibly due to host-driven selective pressures associated with biotrophic and hemibiotrophic parasitism.

**FIGURE 1:**
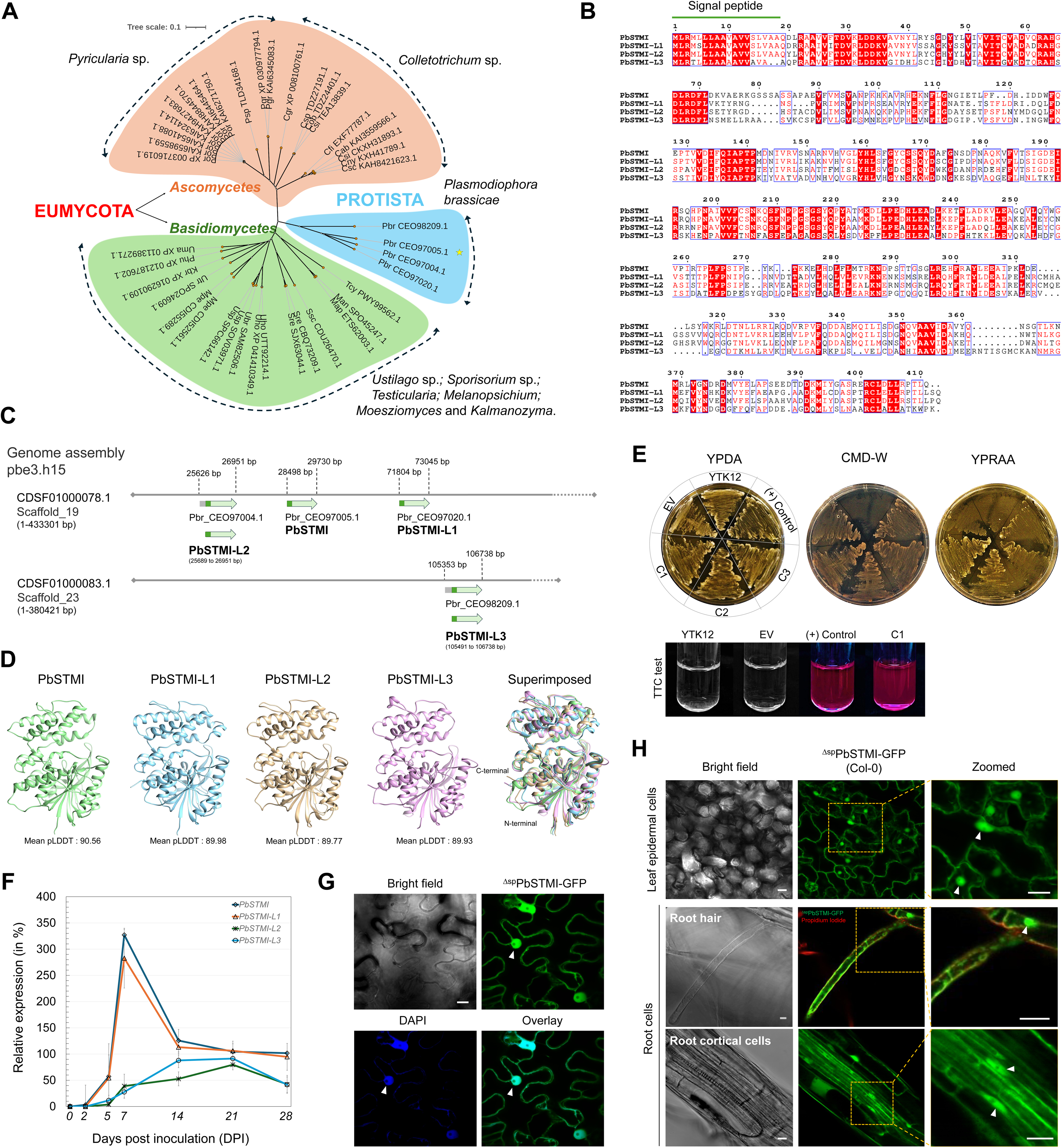
Cross-kingdom conservation, genome assembly, sequence and structural modelling of the PbSTMI effector family. A. Phylogenetic tree generated with Clustal Omega from full-length amino acid sequences of PbSTMI orthologs identified by BlastP searches against the NCBI non-redundant database and visualized with ITOL. The orthologs from three broad plant pathogen groups form separate phylogenetic clusters, each shaded in a different color. Ascomycetes (orange) and Basidiomycetes (green) belong to the super-kingdom Eumycota, while *P. brassicae* belongs to the kingdom Protista (cyan). The yellow star denotes the *P. brassicae* effector PbSTMI (Pb_CEO97005.1), the input sequence for the BlastP searches. B. Amino acid sequence alignment of PbSTMI and its paralogs using Clustal Omega. Signal peptides were predicted using SignalP 5.0 and indicated with a green line. Highly conserved amino acids are highlighted in red with conserved sequences boxed in blue. The image was created using the ESPript 3.0 server. C. Genomic locations of the PbSTMI gene and its three paralogs in *P. brassicae* pbe3.h15 genome assembly. The four PbSTMI genes are distributed in two different pbe3.h15 genome assemblies, SGFA01000078.1 (Pbr_CEO97004.1, Pbr_CEO97005.1, and Pbr_CEO97020.1) and SGFA01000083.1 (Pbr_CEO98209.1). The genomic regions (in base pairs) for each paralog are marked with box arrows; the hatched pattern within the arrows indicates N-terminal extension sequences identified by ATGpr. The green arrows represent Pbr_CEO97005.1 (PbSTMI), Pbr_CEO97020.1 (PbSTMI-L1), Pbr_CEO97004.1 (PbSTMI-L2), and Pbr_CEO98209.1 (PbSTMI-L3), along with their respective genomic locations, with the dark green segment of each arrow representing the predicted signal peptide sequence. D. 3D models of PbSTMI (green), PbSTMI-L1 (cyan), PbSTMI-L2 (gold), and PbSTMI-L3 (magenta) generated using AlphaFold 2.0 at Neurosnap and visualized using UCSF Chimera. The best protein model for each effector, based on the highest mean pLDDT score, was selected from five generated models. E. Signal trap assay, using the yeast secretion system, validating the functionality of the predicted signal peptide (SP). All transformants of the invertase-negative yeast strain (YTK12) grew on CMD-W; however, only YTK12 transformants carrying the SP^PbSTMI^ grew on YPRAA selective media and reduced colorless TTC to a red-colored formazan. Positive control was the SP from Arabidopsis low-molecular-weight cysteine-rich 78 (*At*LCR78). Three independent biological replicates, C1, C2, and C3, were plated with comparable results. F. Relative expression profiles of *PbSTMI* and the three *PbSTMI* paralogs, *PbSTMI-L1* (*Pbr_CEO97020.1*), *PbSTMI-L2* (Pbr_CEO97004.1), and *PbSTMI-L3* (*Pbr_CEO98209.1*) at 0, 2, 5, 7, 14, 21, and 28 days post-inoculation (DPI) during clubroot infection in Arabidopsis. G. Nucleo-cytoplasmic localization of ^Δsp^PbSTMI-GFP in leaf epidermal cells of *N. benthamiana*. White arrowheads indicate nuclear colocalization between ^Δsp^PbSTMI-GFP and DAPI fluorescence signals. Scale bar = 10 µm. H. Subcellular localization of ^Δsp^PbSTMI-GFP in Arabidopsis overexpression (OE) lines. The green channel shows the nuclear and cytoplasmic localization of ^Δsp^PbSTMI-GFP in leaf epidermal cells (top panel) and root tissues (bottom panel). White arrowheads indicate the nuclear localization of ^Δsp^PbSTMI-GFP. Scale bars = 10 μm.

Three additional PbSTMI family members were identified: PbSTMI-L1, L2, and L3, based on AA sequence identity (**Figure 1A-C**) and structural conservation (**Figure 1D**), suggesting possible functional redundancy in *P. brassicae*-host interactions. Both PbSTMI and PbSTMI-L1 were predicted to have a functional N-terminal signal peptide (SP) necessary for effector secretion into host cells (**Figure 1B**) and a SP trap assay confirmed the secretory function of PbSTMI SP (**Figure 1E**). No SP was initially predicted for PbSTMI-L2 and PbSTMI-L3 (**Table S2**), however, analysis with ATGpr predicted SPs between 21-46 amino acids downstream of their originally predicted N termini. Similar approaches were used to predict SP containing PbSTMI orthologs in other phytopathogens (**Table S1**). The MSA of PbSTMI and representative orthologs from each cluster support cross-kingdom structural conservation of STMI (**Figure S1A,B; Data S1**). No STMI orthologs were identified from necrotrophic fungi.

Transcript levels of PbSTMI and its paralogs during primary infection (0, 2, 5 DPI) and secondary infection (7, 14, 21, 28 DPI) in *P. brassicae*-infected Arabidopsis root tissues demonstrated two distinct expression patterns: *PbSTMI* and *PbSTMI-L1* transcripts appear between 2-5 DPI, peak at 7 DPI, then decline from 7-14 DPI before plateauing at a higher level than *PbSTMI-L2* and *PbSTMI-L3*, that are upregulated from 5 DPI through late secondary infection, peaking at 21 DPI before declining through 28 DPI (**Figure 1F**). No transcripts were detected in *P. brassicae* resting spores before infection.

Transiently expressed in *N. benthamiana*, ^Δsp^PbSTMI-GFP showed nucleo-cytoplasmic localization (**Figure 1G**), with small cytoplasmic puncta (**Figure S2**). Arabidopsis 35S::^Δsp^PbSTMI-GFP overexpression (OE) lines exhibited a similar nucleo-cytoplasmic localization in both leaves and root tissues (**Figure 1H**).

### 2.2. ^Δsp^PbSTMI modulates flg22-triggered, salicylic acid (SA)-dependent, late PTI responses and increases susceptibility to *P. brassicae* and *Colletotrichum higginsianum*

To investigate the function of ^Δsp^PbSTMI and its role during infection, transgenic Arabidopsis 35S::^Δsp^PbSTMI-GFP OE lines were examined for immune responses. No discernible phenotypic differences were observed between ^Δsp^PbSTMI-GFP OE lines and wild-type (WT; Col-0) plants in the absence of infection **(Figure 2A)**. When treated with flg22, to induce PAMP-trigged immunity (PTI), ^Δsp^PbSTMI-GFP OE lines did not exhibit significant changes in reactive oxygen species (ROS) accumulation, compared to WT **(Figure 2B)**; however, a suppression of late-stage ROS accumulation was observed 18 hours after infiltration **(Figure S3A)**. Callose deposition was significantly reduced in ^Δsp^PbSTMI-GFP OE lines compared to WT **(Figure S3B)** and an attenuated upregulation of the SA-marker genes *PR1* and *PR2* **(Figure 2C)** was observed in OE lines compared to WT. Similarly, transcript levels of *AtICS1*, a key enzyme in SA biosynthesis, were substantially lower in ^Δsp^PbSTMI-GFP OE plants than in WT when challenged with flg22 **(Figure 2C)**.

**FIGURE 2:**
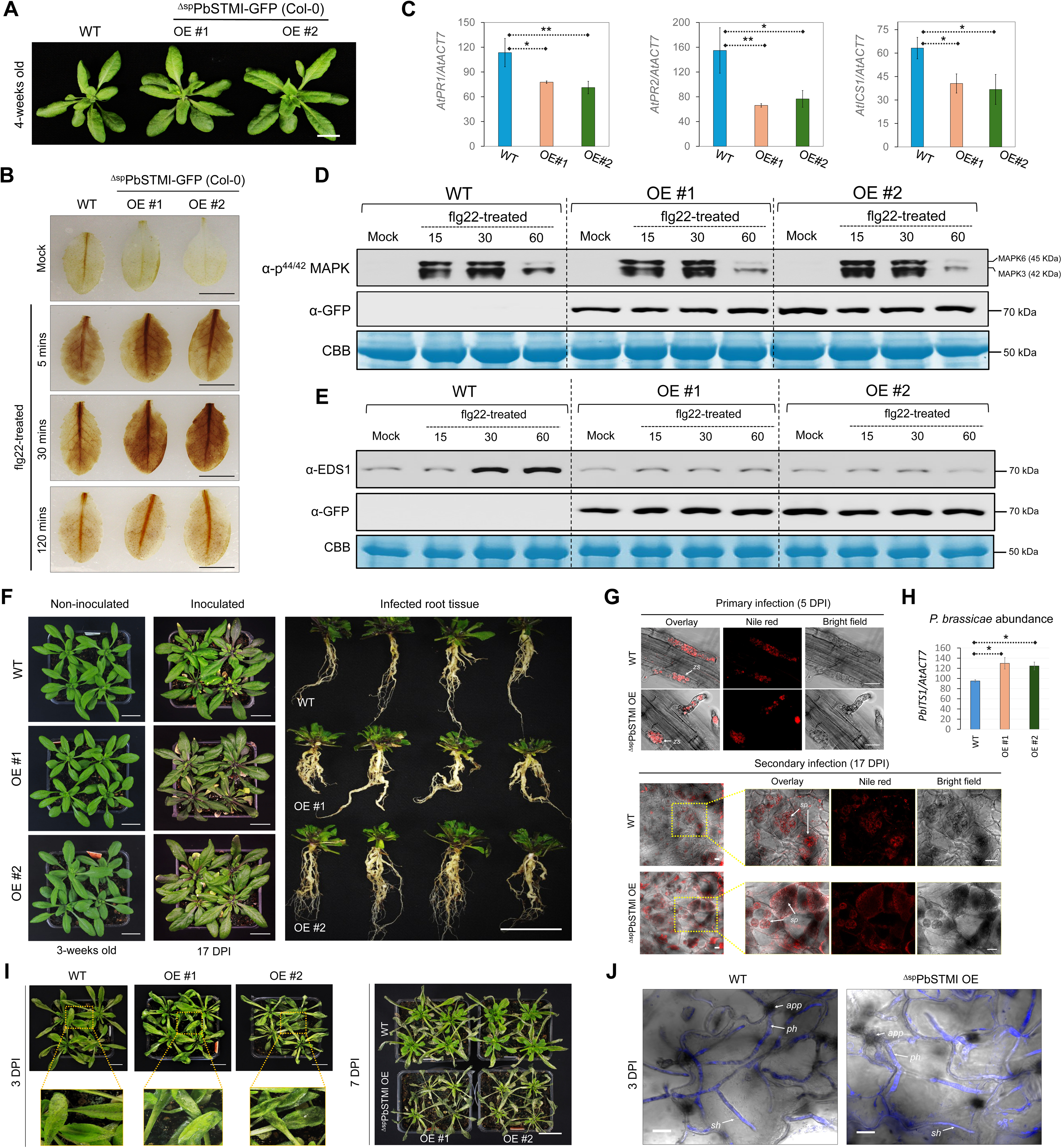
Expression of ^Δsp^PbSTMI in Arabidopsis leads to decreased flg22-triggered late PTI responses and increased susceptibility to *P. brassicae* and *Colletotrichum higginsianum*. (A). Above-ground phenotypes of four week-old WT and two independent Arabidopsis ^Δsp^PbSTMI-GFP lines. Scale bar = 1 cm. (B). Leaves of four week-old Arabidopsis treated with 1 µM flg22 and assessed for ROS through DAB staining for H_2_O_2_ to detect early PTI responses at 5, 30 and 120 min post-infiltration. Scale bars = 1 cm. (C). Relative expression analysis of *AtPR1*, *AtPR2*, and *AtICS1* from four week-old soil-grown plants 18 hours post-infiltration with 1 µM flg22. Expression was normalized relative to Arabidopsis *AtACTIN7* (At5g09810). (D). Western blots from leaves of WT and ^Δsp^PbSTMI-GFP lines 15, 30, and 60 min post-infiltration with 1 µM flg22 or H_2_O. Equal loading was assessed by Coomassie Brilliant Blue (CBB) staining and the molecular weights (kDa) of GFP, MAPK3, and MAPK6 are provided. This experiment was repeated three times with consistent results. (E). Western blots of flg22-treated samples from WT and ^Δsp^PbSTMI-GFP lines using anti-EDS1 to detect endogenous AtEDS1 protein level. The anti-GFP antibody detects expressed ^Δsp^PbSTMI-GFP in OE lines. This experiment was repeated three times with consistent results. (F). Arabidopsis ^Δsp^PbSTMI OE lines at 17 DPI show greater symptoms of clubroot disease, both in above- and below-ground tissues, compared to WT. Scale bar = 2 cm. (G). Representative microscopic images of the primary (5 DPI) and secondary (17 DPI) infection stages of clubroot development in *P. brassicae*-inoculated plants. Nile red stains (red) lipid droplets of *P. brassicae* developmental structures in infected plant root tissues, showing zoosporangia at primary infection (5 DPI) of epidermal cells and secondary plasmodia during secondary cortical infection at 17 DPI. *zs*, zoosporangia; *sp*, secondary plasmodia. Scale bar = 20 µm. (H). Increased *P. brassicae* abundance (percentage) in below-ground tissues of infected Arabidopsis WT and ^Δsp^PbSTMI OE lines as quantified by PCR of genomic DNA using *PbITS1* as the marker for *P. brassicae* and *AtACT7* as the plant-specific control for relative abundance measurements. Mean percentages are presented with error bars ±SD. Three independent biological replicates were performed, and statistical differences were assessed using one-way analysis of variance (ANOVA) followed by a post-hoc Tukey’s HSD multiple comparison test. (*) and (**) denote data points showing significant differences at P value < 0.05 and P < 0.01, respectively. (I). PbSTMI lines, at 3 DPI, show increased susceptibility to hemi-biotrophic *C. higginsianum* that causes anthracnose infection in plants. Inset images show water-soaking lesions in the infected leaves. Infected ^Δsp^PbSTMI OE lines at 7 DPI show a complete collapse of above-ground tissues. Scale bar = 2 cm. (J). *C. higginsianum-*infected leaf epidermal tissues at 3 DPI, showing the development of primary and secondary hyphae in the ^Δsp^PbSTMI OE line compared to WT. Infected leaf tissues were stained with trypan blue. *app*, appressorium; *ph*, primary hyphae; *sh*, secondary hyphae. Scale bars = 10 µm.

Despite the suppression of flg22-induced callose deposition and defense gene expression, ^Δsp^PbSTMI-GFP lines showed no significant difference in MAPK3/6 phosphorylation (activation) profiles relative to WT (**Figure 2D**), suggesting that ^Δsp^PbSTMI does not inhibit rapid and transient early PTI responses but rather modulates flg22-induced downstream immune responses that occur independently of MAPK.

EDS1 functions as a central immune signaling hub integrating responses from both PRR-and NLR-mediated defense pathways. As expected in WT plants, EDS1 levels increased significantly 30 and 60 minutes post-flg22 infiltration, while no notable EDS1 accumulation was seen in the ^Δsp^PbSTMI-GFP OE lines **(Figure 2E)**. Together, these data suggest that ^Δsp^PbSTMI acts downstream of early PTI signaling in Arabidopsis, modulating late-stage SA-mediated defenses and EDS1 accumulation.

Having demonstrated that ^Δsp^PbSTMI-GFP OE attenuates immune signaling, we next examined whether OE altered disease susceptibility. Compared to WT, ^Δsp^PbSTMI-GFP OE lines exhibited significantly greater clubroot disease severity at 17 DPI (**Figure 2F**) and a substantially higher pathogen load, as determined by microscopic examination **(Figure 2G)** and PCR quantification of *P. brassicae* genomic DNA from infected roots (**Figure 2H**).

When exposed to the hemibiotrophic anthracnose fungus *Colletotrichum higginsianum* at 3 DPI, ^Δsp^PbSTMI-GFP OE leaves exhibited widespread water-soaked lesions compared with the limited symptoms in WT leaves (**Figure 2I**). Microscopic examination revealed invasive primary hyphae and thin necrotrophic secondary hyphae associated with necrotic lesion development at 3 DPI (**Figure 2J**). Between 5 and 7 DPI, aggressive tissue maceration rapidly spread across ^Δsp^PbSTMI-GFP OE leaves, leading to complete tissue collapse by 7 DPI, whereas infected WT leaves showed partial recovery with renewed meristematic growth (**Figure 2I**). These observations suggest that ^Δsp^PbSTMI suppresses immunity in Arabidopsis against a diverse spectrum of phytopathogens.

### 2.3. PbSTMI interacts with and targets AtEDS1 for proteasome-mediated degradation, disrupting its nucleo-cytoplasmic balance

Targeted Y2H screening of potential host interactors from SA-dependent and/or R-gene-mediated defense pathways confirmed a direct interaction between ^Δsp^PbSTMI and AtEDS1 (**Figure 3A,B**), that was further validated *in planta* by bimolecular fluorescence complementation (BiFC) of ^Δsp^PbSTMI-nYFP transiently co-expressed with AtEDS1-cYFP, and vice versa. (**Figure 3C**) and co-immunoprecipitation (Co-IP) assays involving transiently co-expressed ^Δsp^PbSTMI-GFP and HA-AtEDS1 in *N. benthamiana* leaves, followed by protein pull-down using GFP-trap agarose (**Figure 3D**). Reciprocal Co-IP confirms the ^Δsp^PbSTMI-AtEDS1 interaction *in planta* (**Figure 3E)**. Transient co-expression of free GFP and HA-AtEDS1 in *N. benthamiana* leaves served as a negative control.

**FIGURE 3:**
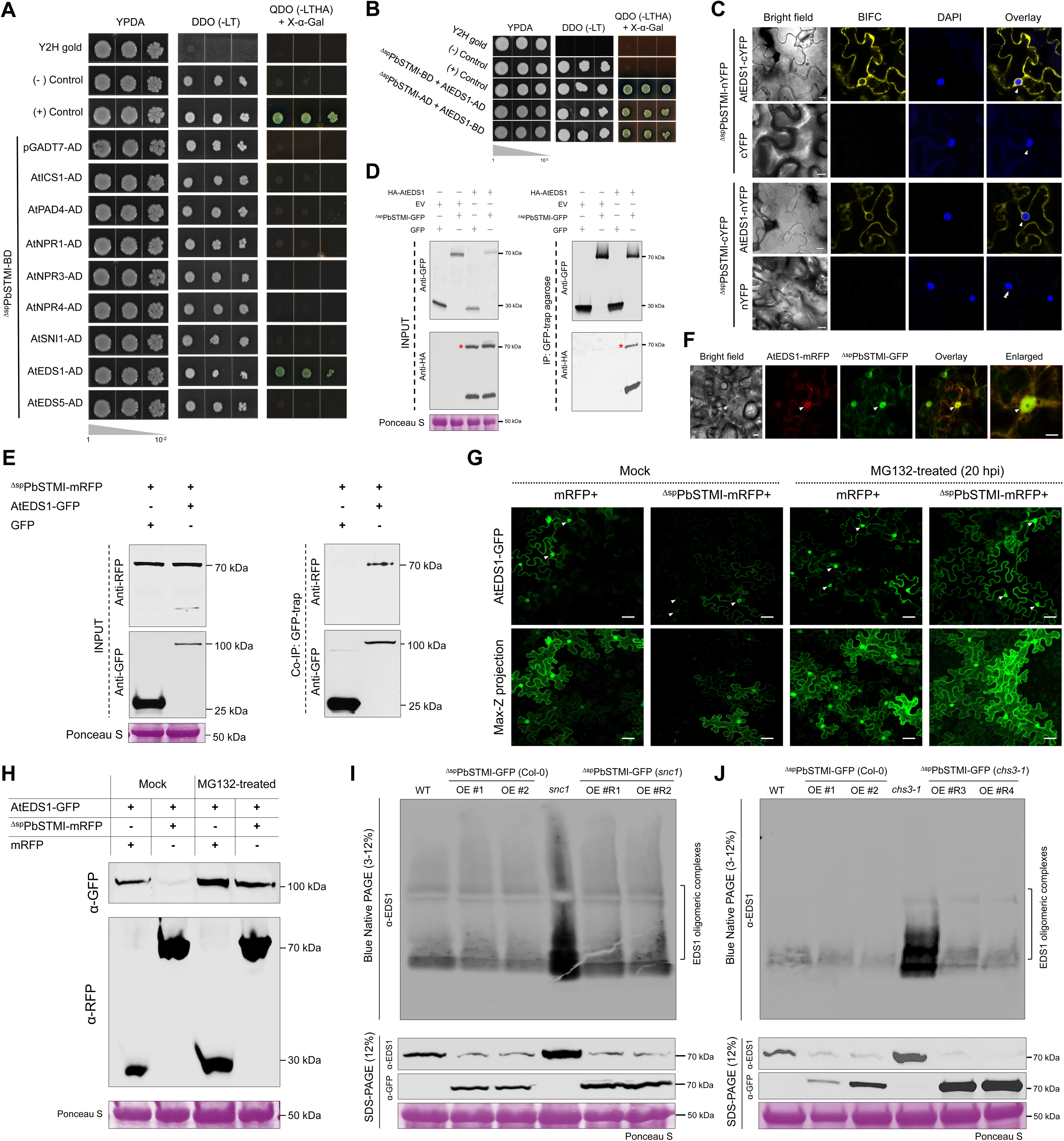
^Δsp^PbSTMI interacts with and degrades AtEDS1 via proteasome-mediated degradation, thereby regulating AtEDS1 oligomerization. (A). Y2H assay identification of host target protein AtEDS1 from Arabidopsis. BD, GAL4 DNA binding domain; AD, GAL4 DNA activation domain; DDO, double dropout; QDO, quadruple dropout. Arabidopsis proteins: ICS1-Isochorismate synthase 1, PAD4- Phytoalexin deficient 4, NPR1- Non-expressor of PR gene 1, NPR3- Non-expressor of PR gene 3, NPR4- Non-expressor of PR gene 4, SNI1- Suppressor of NPR1-1, EDS1- Enhanced disease susceptibility 1, and EDS5- Enhanced disease susceptibility 5. pGADT7-AD, empty vector control; pGADT7-T interaction with pGBKT7-53, positive control; pGADT7-T interaction with pGBKT7-Lam, negative control. (B). ^Δsp^PbSTMI interacts with AtEDS1 in both BD and AD orientations—the same controls as in A. (C). ^Δsp^PbSTMI interacts with AtEDS1 *in planta* in BIFC assays. *A. tumefaciens* containing the indicated construct combinations was co-infiltrated (OD_600_ = 0.5) into *N. benthamiana* leaves. Empty vector with nYFP or cYFP plus ^Δsp^PbSTMI-cYFP or nYFP were the negative controls. Confocal images were obtained 2 days post-infiltration. Positive interactions, marked by complemented YFP signals, colocalized with DAPI nuclear labeling, indicating extranuclear interactions between ^Δsp^PbSTMI and AtEDS1. (D). Co-immunoprecipitation assay validation of ^Δsp^PbSTMI-AtEDS1 interaction using GFP-trap agarose beads after the 26S proteasome inhibitor MG132-treatment for stabilization of the interaction. Anti-GFP and anti-HA antibodies were used to detect free GFP, ^Δsp^PbSTMI-GFP and HA-AtEDS1, respectively. Red stars indicate the expected molecular weight of the desired protein. (E). Co-immunoprecipitation assay further validates the interaction between ^Δsp^PbSTMI-mRFP and AtEDS1-GFP using GFP-trap agarose beads. Protein loading is shown by Ponceau S staining of Rubisco large subunit and the molecular weights (kDa) of GFP, mRFP, AtEDS1-GFP, ^Δsp^PbSTMI-mRFP, and RuBisCo LSU are provided. An anti-GFP antibody was used to detect free GFP and AtEDS1-GFP, while an anti-RFP antibody was used to detect ^Δsp^PbSTMI-mRFP and free mRFP. (F). Co-localization of ^Δsp^PbSTMI-GFP and AtEDS1-mRFP during transient expression in *N. benthamiana* leaves. The overlay image indicates nuclear and cytoplasmic co-localization of ^Δsp^PbSTMI-GFP and AtEDS1-mRFP in *N. benthamiana*. Scale bar = 10 µM. (G). Comparative confocal images show AtEDS1-GFP fluorescence in combination with ^Δsp^PbSTMI-mRFP and the control, after treatment with MG132 in *N. benthamiana* leaves. Scale bar = 10 µm. (H). Immunoblot detection of co-expressed proteins with and without MG132 treatment; mRFP, GFP, ^Δsp^PbSTMI-mRFP and AtEDS1-GFP using anti-GFP and anti-RFP, as indicated. (I-J). BN-PAGE and western blots were performed on lysates from WT, ^Δsp^PbSTMI-GFP, *snc1*, ^Δsp^PbSTMI-GFP (*snc1*), *chs3-1* and ^Δsp^PbSTMI-GFP (*chs3-1*) lines. AtEDS1-oligomeric associations were detected using anti-EDS1 antibody, as indicated by an open square bracket on the right side of the blot. Western blots of the same lysates using anti-EDS1 and anti-GFP antibodies confirmed the levels of AtEDS1. These experiments were repeated three times with consistent results.

Nuclear and cytoplasmic localization of AtEDS1 has been reported previously, with a balanced distribution between the two compartments essential for optimal plant defense responses; notably, extranuclear EDS1 significantly impairs EDS1-dependent immune functions in plants (42). Both the fluorescence protein-tagged ^Δsp^PbSTMI and AtEDS1 independently localize to the nucleus and cytoplasm, and a reduced AtEDS1/^Δsp^PbSTMI fluorescence was observed in transiently co-expressing *N. benthamiana* cells (**Figure 3F; Figure S4A; Movie S1**). EDS1 accumulation in plants is controlled by 26S proteasome-mediated degradation, and an AvrPphB susceptible 3 (PBS3)-EDS1 interaction protects EDS1 from this process (43,44). To determine whether ^Δsp^PbSTMI-mediated AtEDS1 turnover occurs through the 26S proteasome pathway, the potential ^Δsp^PbSTMI-AtEDS1 interaction was tested transiently with the proteasome inhibitor MG132. As expected, mock-treated ^Δsp^PbSTMI-mRFP/AtEDS1-GFP co-infiltrated leaves showed decreased AtEDS1 fluorescence compared to control AtEDS1-GFP infiltrated leaves. However, in the presence of the proteasome inhibitor, a significant increase in AtEDS1 fluorescence is seen in ^Δsp^PbSTMI-mRFP/AtEDS1-GFP co-infiltrated leaves relative to mock-treated leaves (**Figure 3G**). Similarly, a reduced AtEDS1 protein level was observed in ^Δsp^PbSTMI-mRFP/AtEDS1-GFP co-infiltrated leaves compared to controls, with an increased level of AtEDS1 seen in MG132-treated ^Δsp^PbSTMI-mRFP/AtEDS1-GFP co-infiltrated leaves (**Figure 3H**). Additionally, BiFC co-infiltration with DAPI staining **(Figure 3C; Figure S4B)** supports an extranuclear interaction between ^Δsp^PbSTMI and AtEDS1, with the resulting ^Δsp^PbSTMI-AtEDS1 complex predominantly retained in the cytoplasm. However, MG132 treatment resulted in a significant nuclear enrichment of the ^Δsp^PbSTMI-AtEDS1 complex with cytosolic aggregates **(Figure S4B)**, suggesting that the ^Δsp^PbSTMI-AtEDS1 complexes are degraded via the 26S proteasome, disrupting the nucleo-cytoplasmic balance of EDS1 in plants.

Homomeric EDS1 dimerization promotes association with PAD4/SAG101 (45,46), and TIR-NLRs like SNC1 and CHS3 GOF mutants, *snc1* and *chs3-1*, depend on EDS1 for resistance (47,48). To investigate the effect of ^Δsp^PbSTMI on EDS1 oligomerization in *snc1* and *chs3-1* plants, ^Δsp^PbSTMI-GFP OE lines were created in each mutant background. BN-PAGE analysis showed that increased AtEDS1 levels in *snc1* and *chs3-1* are associated with increased AtEDS1 oligomeric complexes when compared to WT (**Figure 3I, J**). The ^Δsp^PbSTMI-GFP (*snc1*) lines and ^Δsp^PbSTMI-GFP (*chs3-1*) lines showed reduced EDS1 oligomers compared to the *snc1* and *chs3-1* mutants (**Figure 3I,J**). Similarly, SDS-PAGE analysis of both ^Δsp^PbSTMI-GFP (*snc1*) and ^Δsp^PbSTMI-GFP (*chs3-1*) lines showed a significant decrease in AtEDS1 accumulation **(Figure 3I,J)**, suggesting that the reduction in AtEDS1 oligomeric complexes is associated with ^Δsp^PbSTMI-mediated AtEDS1 degradation rather than a specific effect on EDS1 oligomerization.

### 2.4. PbSTMI rescues *snc1* and partially suppresses *chs3-1* autoimmunity

A recent report suggests SNC1 acts as a sensor NLR, activating TIR-NLR-triggered resistance by guarding helper NLRs targeted by bacterial effectors (29). Overexpression or GOF mutants of SNC1 activate R-proteins, leading to autoimmune phenotypes such as dwarfism, curly leaves, and loss of apical dominance, even in the absence of pathogens (49). Since ^Δsp^PbSTMI interacts with AtEDS1 (**Figure 3**) and EDS1 is critical for TIR-NLR resistance (47,50), we investigated the effect of ^Δsp^PbSTMI in the AtEDS1-dependent *snc1* autoimmune background. ^Δsp^PbSTMI-GFP expression in ^Δsp^PbSTMI-GFP (*snc1*) lines rescued the autoimmune traits, restoring rosette area and root biomass (**Figure 4A,B; Figure S5A-C**).

**FIGURE 4:**
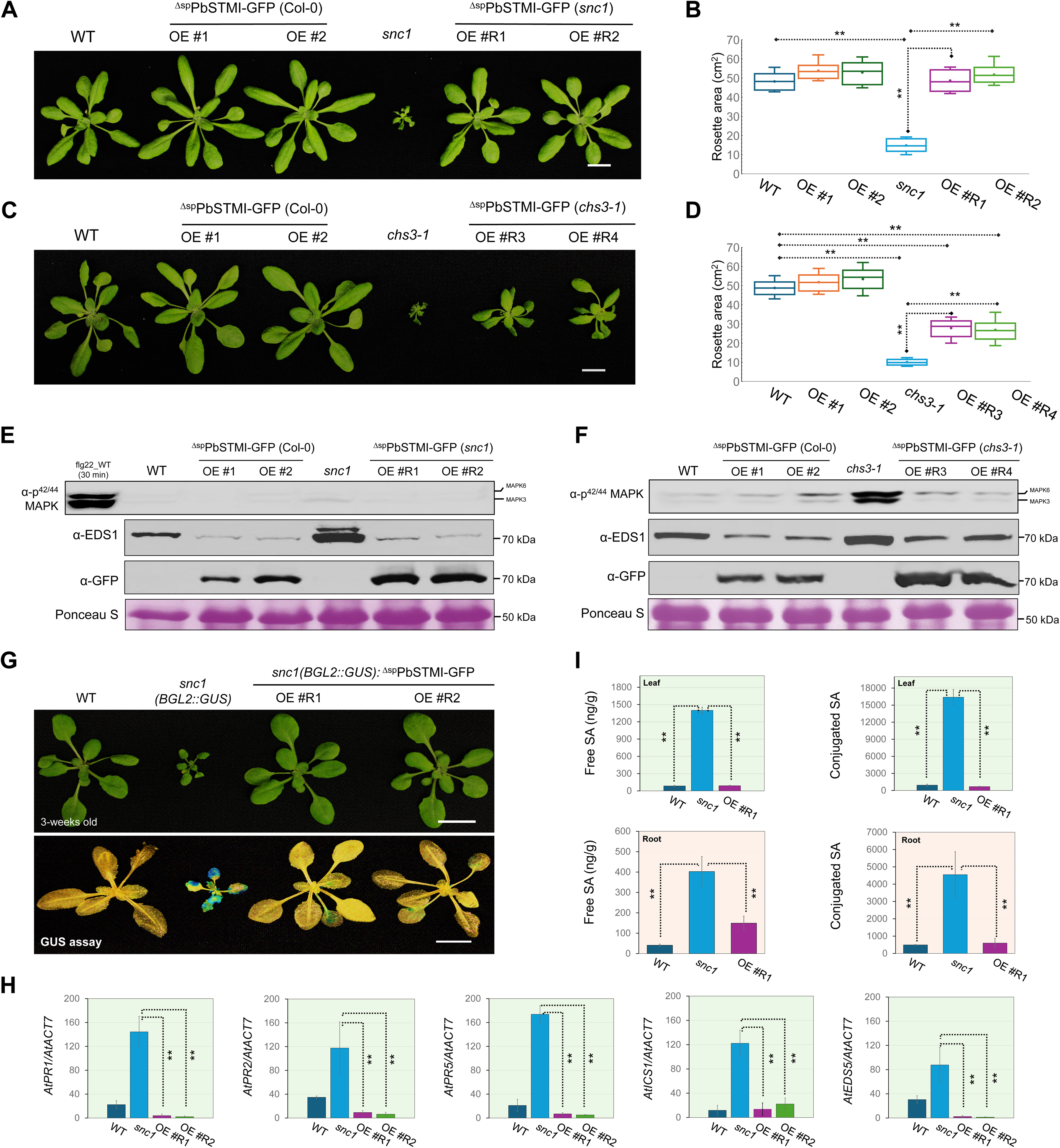
^Δsp^PbSTMI-GFP attenuates *snc1* and *chs3-1* mediated autoimmunity, decreasing EDS1 level while downregulating SA biosynthesis and defense gene expression. (A-D). Vegetative above-ground phenotypes of four week-old soil-grown Arabidopsis WT col-0, *snc1, chs3-1,* and two of each independent ^Δsp^PbSTMI-GFP OE lines in WT, *snc1,* and *chs3-1* backgrounds, as indicated. Scale bar = 1 cm. The quantified rosette area (cm²) for the independent genotypes is presented as color-coded box plots, with “x” denoting the mean and the horizontal line within each box indicating the median. n = 10 individual plants. Statistical differences were assessed with one-way ANOVA followed by a post-hoc Tukey’s HSD multiple comparison test, and data points indicated with (*) and (**) show significant differences at P value < 0.05 and P < 0.01, respectively. (E-F). Immunoblots of phosphorylated MAPK3/6, AtEDS1 and ^Δsp^PbSTMI-GFP in the indicated genotypes using anti-p^42/44^ MAPK, anti-EDS1 and anti-GFP antibodies, respectively. flg22-treated WT was the positive control for MAPK phosphorylation. Equal loading is shown by Ponceau S staining of the Rubisco large subunit and the molecular weights (kDa) of GFP, EDS1 and Rubisco LSU are provided. This experiment was repeated three times with consistent results. (G). Vegetative morphology of three week-old soil-grown WT (Col-0), *snc1* and two ^Δsp^PbSTMI-GFP (*snc1*) lines. Three-week-old soil-grown plant rosettes were stained for beta-glucuronidase (GUS) activity. Scale bar = 1 cm. (H). Gene expression analysis of Arabidopsis *PR1, PR2, PR5, ICS1* and *EDS5* in three week-old soil-grown plants. Relative expression of each gene was determined with respect to *AtACTIN7* (At5g09810). (I). Free and conjugated SA (ng/g) in foliar and root tissues of WT (Col-0), *snc1* and ^Δsp^PbSTMI-GFP (*snc1*) line. Error bars represent mean ±SD. Statistical differences were assessed using one-way analysis of variance (ANOVA) followed by a post-hoc Tukey’s HSD multiple comparison test, and data points with significant differences are indicated with (*) and (**) at P values < 0.05 and < 0.01, respectively. Three independent biological replicates were performed.

CHS3 is also a TIR-NLR but with a C-terminal LIM-peptidase (LP) integrated domain (ID) that acts as a decoy for effector recognition (51,52). The GOF *chs3-1* mutation in the LIM domain activates EDS1-dependent autoimmune responses (48). Unlike the complete rescue in the ^Δsp^PbSTMI-GFP (*snc1*) lines, ^Δsp^PbSTMI-GFP (*chs3-1*) lines partially rescue the *chs3-1* phenotype, showing intermediate size (**Figure 4C**), rosette area (**Figure 4D**), and root mass (**Figure S5D-F**) compared to WT and *chs3-1* plants, indicating that ^Δsp^PbSTMI partially suppresses TIR-NLR-triggered autoimmune phenotypes in these plants.

Both PTI and ETI often elicit common downstream cellular immune responses, including Ca^2+^ influx, apoplastic ROS accumulation, callose deposition, and activation of the MAPK cascade. When WT, *snc1*, *chs3-1*, ^Δsp^PbSTMI-GFP (*snc1*), and ^Δsp^PbSTMI-GFP (*chs3-1*) lines were monitored for TIR-NLR-triggered ROS accumulation, callose deposition, and MAPK activation, elevated ROS levels were observed in both *snc1* and *chs3-1* plants, whereas ^Δsp^PbSTMI-GFP (*snc1*) and ^Δsp^PbSTMI-GFP (*chs3-1*) lines showed WT levels of ROS (**Figure S6A,B**). *snc1* plants showed no significant increase in MAPK activation (**Figure 4E**) or callose deposition, supporting the absence of spontaneous cell death in this autoimmune mutant (**Figure S6C**). Conversely, *chs3-1* autoimmunity, marked by heightened MAPK activation (**Figure 4F**) and strong callose deposition (**Figure S6D**), is associated with a cell death phenotype (**Figure S7**). These immune responses were notably suppressed in ^Δsp^PbSTMI-GFP (*chs3-1*) lines, indicating that PbSTMI suppresses the cell death phenotype via suppression of EDS1-dependent TIR-NLR-triggered immunity **(Figure 4F; Figure S6B,D).** Furthermore, the PbSTMI-GFP (*snc1*) and PbSTMI-GFP (*chs3-1*) lines have significantly lower EDS1 levels than their autoimmune backgrounds, similar to or less than WT, supporting the suggestion that ^Δsp^PbSTMI-mediated EDS1 turnover delivers the phenotypic rescue seen in these lines (**Figure 4E,F**).

*snc1* plants contain the constitutively active *BGL2 (PR2)::GUS* reporter expression cassette that is significantly suppressed in *snc1 eds1* plants (47,53). GUS expression is similarly suppressed in the leaves of ^Δsp^PbSTMI-GFP (*snc1*) lines (**Figure 4G**), together with decreased transcript levels of the SA-responsive PR genes, *PR1, PR2,* and *PR5,* relative to the high, constitutively expressed levels in *snc1* **(Figure 4H)**. Likewise, transcript levels of SA biosynthesis and transport genes, *ISOCHORISMATE SYNTHASE 1* (*ICS1*) and *ENHANCED DISEASE SUSCEPTIBILITY 5* (*EDS5*), were significantly lower in ^Δsp^PbSTMI-GFP (*snc1*) lines compared to *snc1* plants (**Figure 4H**).

Quantitative analysis of leaves and roots of *snc1*, WT and ^Δsp^PbSTMI-GFP (*snc1*) plants showed elevated levels of both free and conjugated SA in *snc1* compared to WT and ^Δsp^PbSTMI-GFP (*snc1*) plants, with inactive conjugated SA more than 10-fold higher than free active SA in all plants (**Figure 4I**). WT levels of free SA in ^Δsp^PbSTMI-GFP (*snc1*) plants, together with decreased *ICS1* transcript levels, suggest that ^Δsp^PbSTMI suppresses SA biosynthesis and thereby suppresses the autoimmune responses of *snc1* in an EDS1-dependent manner.

Plant NLR-mediated immunity is tightly balanced to avoid detrimental effects on plant growth. In Arabidopsis, SNC1 is tightly regulated by the F-box protein CPR1 via the SCF proteasome complex, and the *cpr1* mutant exhibits a *snc1-1*-like autoimmune phenotype due to hyperaccumulation of the active SNC1 receptor, thereby activating downstream signaling (54). To determine whether ^Δsp^PbSTMI inhibits TIR-NLR-triggered immunity resulting in increased EDS1 protein turnover by promoting upstream degradation of TIR-NLR receptors via the ubiquitin-proteasome system, AtSNC1 protein was transiently co-expressed with ^Δsp^PbSTMI-GFP with or without the proteosome inhibitor MG132 **(Figure S8A)**. The protein level of AtSNC1 did not change upon addition of MG132 **(Figure S8A)**, and PbSTMI showed no direct interaction with AtSNC1 *in planta* **(Figure S8B)**. Furthermore, ^Δsp^PbSTMI-GFP lines generated in the EDS1-independent MECHANOSENSITIVE CHANNEL OF SMALL CONDUCTANCE-LIKE 10 (MSL10) GOF mutant *msl10-3G* (*rea1*) (55,56) showed no meaningful phenotypic change relative to *msl10-3G* (**Figure S9A-C**) indicating that ^Δsp^PbSTMI specifically affects EDS1-dependent plant immunity and that ^Δsp^PbSTMI-mediated suppression of TIR-NLR immunity acts downstream of TIR-NLR overexpression, pointing directly to signaling suppression downstream of AtSNC1, predominantly through a modulation of EDS1-dependent signaling. In all three mutant backgrounds, *snc1*, *chs3-1*, and *msl10-3G*, ^Δsp^PbSTMI-GFP shows a similar nucleo-cytoplasmic localization, with more abundant cytosolic puncta than in ^Δsp^PbSTMI-GFP (Col-0) plants (**Figure S10**).

### 2.5. Conserved N-terminal sequences of PbSTMI are required for *in planta* homo- and hetero-oligomerization and essential for autoimmune rescue in *snc1* plants

Co-IP with GFP- and mRFP-tagged ^Δsp^PbSTMI, from transient expression in *N. benthamiana*, indicated ^Δsp^PbSTMI can form homo-oligomers (**Figure 5A**); confirmed through *in planta* BIFC assays (**Figure 5B**). Previously, we showed that PbSTMI paralogs are expressed at different stages of clubroot infection in Arabidopsis (**Figure 1F**) and to investigate potential hetero-oligomerization, we selected the distant paralog ^Δsp^PbSTMI-L3 (Pbr_CEO98209.1) from the PbSTMI family. Both homo- and hetero-oligomerization between ^Δsp^PbSTMI and ^Δsp^PbSTMI-L3 was demonstrated in Co-IP and BIFC experiments (**Figure 5A,B**), suggesting that PbSTMI-L1 and PbSTMI-L2, with higher similarity to PbSTMI, should similarly form homo- and hetero-oligomers. These interactions could regulate ^Δsp^PbSTMI functions through oligomerization at different stages of clubroot infection, potentially influencing spatio-temporal plant immune responses. Both ^Δsp^PbSTMI-GFP and ^Δsp^PbSTMI-L3-mRFP localize to the nucleus and cytoplasm in *N. benthamiana* leaves (**Figure 5C**).

**FIGURE 5:**
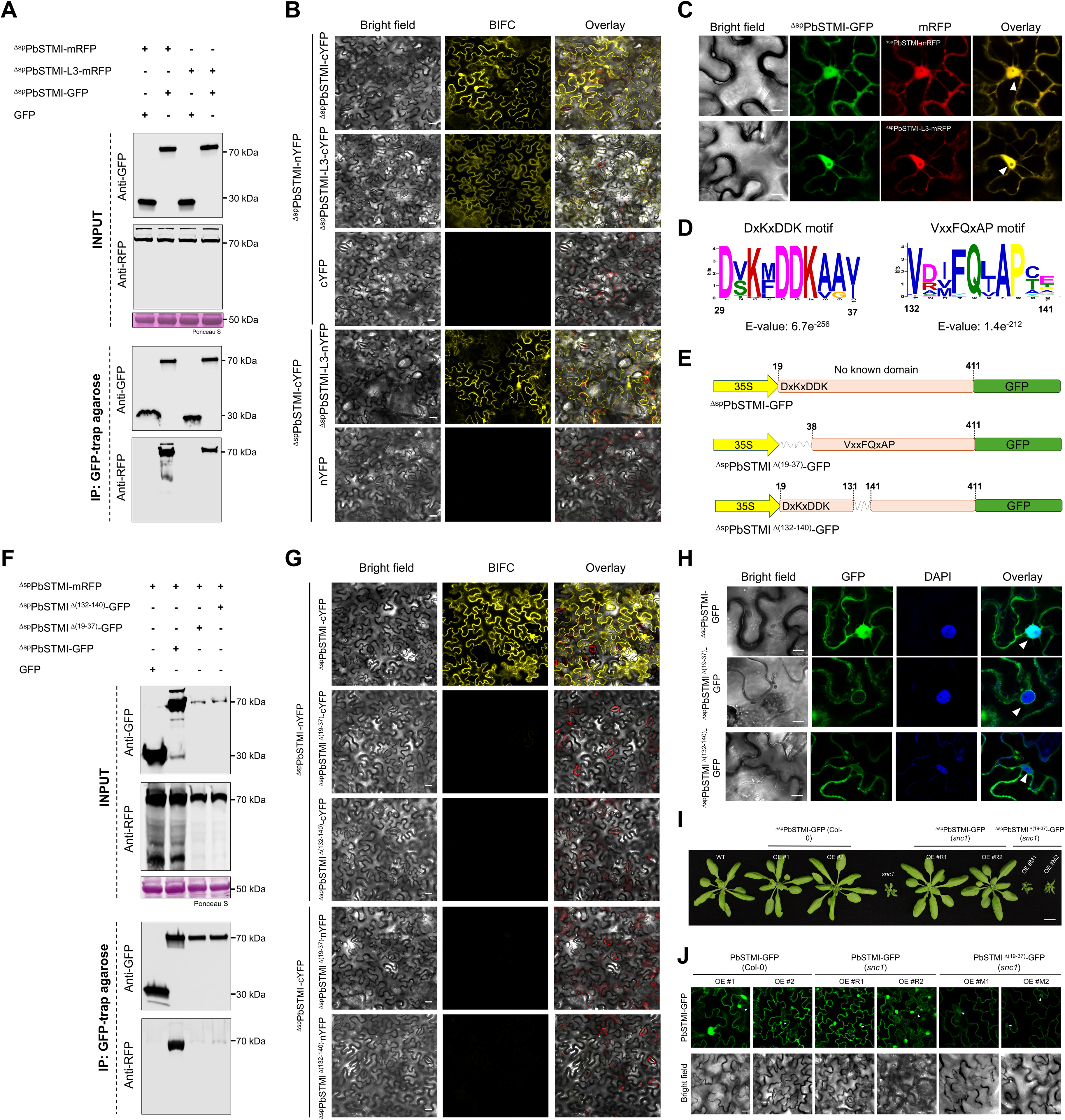
Homo- and hetero-oligomerization of ^Δsp^PbSTMI and ^Δsp^PbSTMI-L3 *in planta* requires conserved N-terminal motifs and is essential for autoimmune rescue in *snc1* plants. (A). Co-immunoprecipitation assay to investigate homo and hetero oligomerization of ^Δsp^PbSTMI interactions using GFP-trap agarose beads. An anti-GFP antibody was used to detect free GFP and ^Δsp^PbSTMI-GFP, while an anti-RFP antibody was used to detect ^Δsp^PbSTMI-RFP and ^Δsp^PbSTMI- L3-RFP. Protein loading is shown by Ponceau S staining of Rubisco large subunit and the molecular weights (kDa) of GFP, ^Δsp^PbSTMI-GFP, ^Δsp^PbSTMI-RFP, ^Δsp^PbSTMI-L3-RFP, and RuBisCo LSU bands are provided. This experiment was repeated three times with consistent results. (B). BIFC assays validating ^Δsp^PbSTMI-oligomerization *in planta*. *A. tumefaciens* containing the indicated construct combinations was co-infiltrated (OD_600_ = 0.5) into *N. benthamiana* leaves. Empty vector with nYFP or cYFP plus ^Δsp^PbSTMI-cYFP or ^Δsp^PbSTMI-nYFP were the negative controls. Confocal images were obtained 2 days post-infiltration. (C). Independent localization of ^Δsp^PbSTMI-GFP and ^Δsp^PbSTMI-L3-mRFP after transient co-expression in *N. benthamiana* leaves. The overlay image demonstrates nuclear (indicated by arrowheads) and cytoplasmic co-localization of ^Δsp^PbSTMI-GFP and ^Δsp^PbSTMI-L3-mRFP in *N. benthamiana*. Scale bar = 10 µM. (D). Conserved ‘DxKxDDK’ and ‘FxxFQxAP’ motifs identified among PbSTMI orthologs (Data S1) by MEME scan with the highest E-value (6.7e^-256^ and 1.4e^-212^, respectively). Visualization of protein motif (in 10-amino acids length) as a logo, where the height of the letter (y-axis) indicates the level of amino acid conservation between the orthologs at the given position (x-axis) in the motif. (E). Domain architectures of the GFP-tagged ^Δsp^PbSTMI and N-terminal mutation constructs under the control of the CaMV 35S promoter. (F). Co-immunoprecipitation assay with ^Δsp^PbSTMI-GFP and GFP-tagged deletion mutants, ^Δsp^PbSTMI^Δ(19–37)^-GFP and ^Δsp^PbSTMI^Δ(132–140)^-GFP, using GFP-trap agarose beads to determine the importance of each motif in ^Δsp^PbSTMI-oligomerization. (G). BIFC validation of lack of ^Δsp^PbSTMI oligomerization with ^Δsp^PbSTMI^Δ(19–37)^-GFP and ^Δsp^PbSTMI^Δ(132–140)^-GFP. Confocal images were obtained 2 days post-infiltration. (H). Confocal images of ^Δsp^PbSTMI^Δ(19–37)^-GFP and ^Δsp^PbSTMI^Δ(132–140)^-GFP showing extra-nuclear localization after DAPI staining. Arrowheads indicate the nuclear and extra-nuclear localizations of ^Δsp^PbSTMI-GFP and ^Δsp^PbSTMI^Δ^(^19–37^)-GFP in *N. benthamiana* leaf epidermal cells. Scale bar = 10 µM. (I). Vegetative above-ground phenotypes of four week-old Arabidopsis WT, *snc1*, ^Δsp^PbSTMI-GFP (Col-0), ^Δsp^PbSTMI-GFP (*snc1*), and ^Δsp^PbSTMI^Δ^(^19–37^)-GFP (*snc1*) lines grown in soil at 22°C. Scale bar = 1 cm. (J). Subcellular localization of ^Δsp^PbSTMI-GFP and ^Δsp^PbSTMI^Δ^(^19–37^)-GFP in Arabidopsis OE lines in the *snc1* autoimmune background. Arrowheads indicate nuclear and extra-nuclear localizations of ^Δsp^PbSTMI-GFP and ^Δsp^PbSTMI^Δ^(^19–37^)-GFP in leaf epidermal cells, respectively. Scale bars = 20 μm.

The N-terminal region of PbSTMI and its orthologs contains two conserved motifs, ^29^DxKxDDK^35^ and ^132^VxxFQxAP^139^ (**Figure 5D**), that are part of a large binding pocket, possibly required for oligomerization ( **Figure S11A**). 3D modelling of ^Δsp^PbSTMI mutants lacking these motifs, ^Δsp^PbSTMI^Δ^(^19–37^) and ^Δsp^PbSTMI^Δ(132–140)^, showed no significant change in conformation (**Figure 5E**; **Figure S11B**); however, no oligomeric interaction of these mutant proteins was observed (**Figure 5F,G**). Both mutant proteins showed reduced fluorescence and protein levels, along with extranuclear localization with cytosolic aggregates, compared to native ^Δsp^PbSTMI (**Figure 5H, Figure S11C,D**), suggesting that ^Δsp^PbSTMI oligomerization is necessary for its nuclear localization and stability *in planta*.

To determine whether oligomerization mediated by the N-terminal sequence is dispensable for ^Δsp^PbSTMI-mediated autoimmune rescue in *snc1*, ^Δsp^PbSTMI^Δ^(^19–37^)-GFP (*snc1*) lines were generated and evaluated for phenotypic rescue. No significant rescue phenotype was observed in these plants, indicating that N-terminal sequence-mediated oligomerization is required for autoimmune rescue (**Figure 5I**). ^Δsp^PbSTMI^Δ^(^19–37^)-GFP exhibited a similar extranuclear localization pattern in these ^Δsp^PbSTMI^Δ^(^19–37^)-GFP (*snc1*) Arabidopsis lines (**Figure 5J**).

### 2.6. ^Δsp^PbSTMI OE in *snc1* restores susceptibility to a diverse range of phytopathogens

To investigate this suggestion, the *P. brassicae* resistance/susceptibility profiles of WT, ^Δsp^PbSTMI-GFP (Col-0), *snc1*, and ^Δsp^PbSTMI-GFP (*snc1*) lines were determined at 21 DPI. WT, ^Δsp^PbSTMI-GFP (Col-0) and ^Δsp^PbSTMI-GFP (*snc1*) plants showed severe above-ground symptoms compared to *snc1*, with ^Δsp^PbSTMI-GFP (Col-0) plants showing more severe symptoms than WT (**Figure 6A**). Infected *snc1* roots showed small clubs and approximately 50% less pathogen load than WT (**Figure 6B,C**). Both primary and secondary infection stages are present in *snc1* clubs (**Figure 6D**), indicating that *snc1* autoimmunity does not prevent *P. brassicae* infection, but that increased resistance is conferred through constitutive host responses. ^Δsp^PbSTMI-GFP (*snc1*) lines showed restored susceptibility, with larger swollen roots (**Figure 6B**) and increased pathogen load (**Figure 6C**), indicating that ^Δsp^PbSTMI disrupts *snc1* resistance to *P. brassicae*.

**FIGURE 6:**
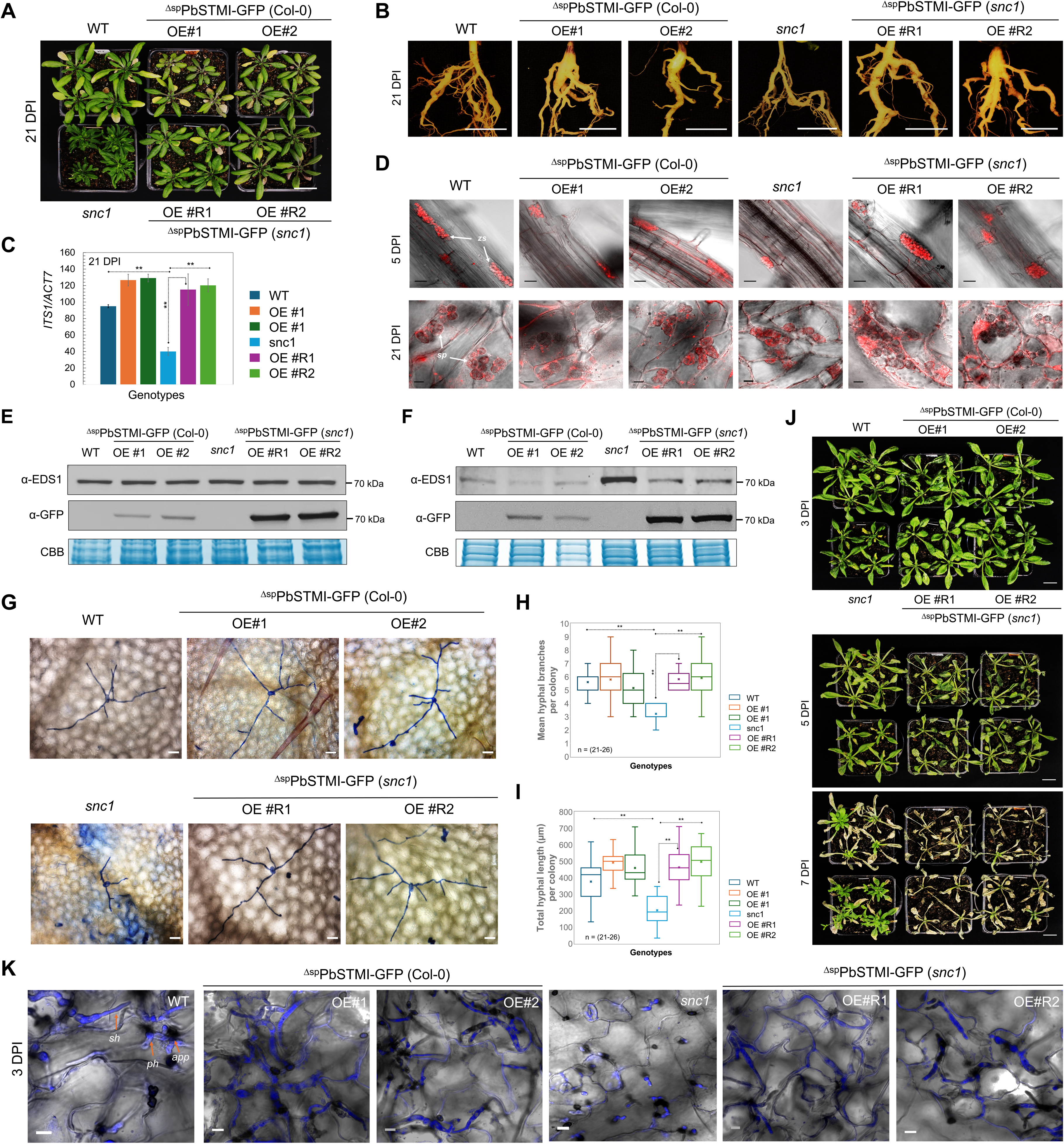
Stable ^Δsp^PbSTMI-GFP expression in Arabidopsis results in restored susceptibility of *snc1* to *P. brassicae, Erysiphe cichoracearum* (powdery mildew) and *C. higginsianum* infection. (A). Above-ground symptoms of representative soil-grown WT (col-0), *snc1*, two ^Δsp^PbSTMI-GFP (Col-0) and two ^Δsp^PbSTMI-GFP (*snc1*) OE lines inoculated with *P. brassicae* pathotype 3 at 21 DPI. (B). Disease symptoms of *P. brassicae*-infected roots in the genotypes as mentioned above at 21 DPI. (C). Pathogen load quantified from total genomic DNA extracted from infected root tissues relative to plant biomass for each genotype. Error bars show mean ±SD. Statistical differences were tested with one-way ANOVA and Tukey’s HSD; P-values indicate significant differences; (*) and (**) denote P < 0.05 and P < 0.01, respectively. n = 3 independent biological replicates. (D). Primary (5 DPI) and secondary (21 DPI) infection stages of clubroot development in *P. brassicae*-inoculated plants. Nile red stains (red) lipid droplets of *P. brassicae* developmental structures in infected plant root tissues. The white arrows indicate infected root epidermis with zoosporangia at primary infection (5 DPI), and secondary plasmodia during secondary cortical infection at 21 DPI. *zs*, zoosporangia; *sp*, secondary plasmodia. (E-F). Immunoblots of ^Δsp^PbSTMI-GFP and AtEDS1 in root tissues of the indicated Arabidopsis genotypes using anti-GFP and anti-EDS1 antibody, respectively; (E) without and (F) 21 DPI with *P. brassicae* infection. Equal loading is shown by CBB staining of total protein and the molecular weight (kDa) of GFP, and EDS1 is provided. This experiment was repeated three times with consistent results. (G). Representative images of *E. cichoracearum*-infected leaves of WT*, snc1,* ^Δsp^PbSTMI-GFP (Col-0) and ^Δsp^PbSTMI-GFP (*snc1*) lines stained with 0.05% aniline blue to show hyphal growth (H-I). Quantitative analysis of *E. cichoracearum* growth on leaves of the above genotypes. Total hyphal mean branch number (H) and hyphal length per colony (I) were scored for each genotype, and the mean values were box-plotted with error bars ±SD. n = the number of *E. cichoracearum* colonies scored for each genotype is 21-26. (*) and (**) indicate data points with significant differences at P value < 0.05 and P < 0.01, respectively. (J). Disease symptoms of WT, ^Δsp^PbSTMI-GFP (Col-0), *snc1* and ^Δsp^PbSTMI-GFP (*snc1*) lines after infection with *C. higginsianum* at 3, 5, and 7 DPI. (K). *C. higginsianum*-infected leaf epidermal tissues at 3 DPI showing much smaller primary hyphae development and no necrotrophic secondary hyphae in *snc1* compared to the bulbous primary hyphae along with thin necrotrophic hyphae into the adjoining cells seen in WT and the ^Δsp^PbSTMI-GFP (Col-0) and ^Δsp^PbSTMI-GFP (*snc1*) lines. Infected leaf tissues were stained with trypan blue—*app*, appressorium; *ph*, primary hyphae; *sh*, secondary hyphae. Scale bars = 10 µm.

While AtEDS1 levels in leaves of *snc1* plants are higher than those of WT, ^Δsp^PbSTMI-GFP (Col-0) or ^Δsp^PbSTMI-GFP (*snc1*) lines (**Figure 4E**), AtEDS1 levels in non-infected root tissues of *snc1* are comparable to those of WT, ^Δsp^PbSTMI-GFP (Col-0) and ^Δsp^PbSTMI-GFP (*snc1*) lines (**Figure 6E**). However, where AtEDS1 level remained high in *P. brassicae-*infected root tissues of *snc1,* it decreased in infected root tissues of WT, ^Δsp^PbSTMI-GFP (Col-0) and ^Δsp^PbSTMI-GFP (*snc1*) lines (**Figure 6F**), suggesting that partial resistance against *P. brassicae* infection in *snc1* is likely due to an increase in SA and subsequent AtEDS1-mediated, SA-dependent immunity in *snc1* roots.

Arabidopsis *eds1* shows increased penetration and compromised non-host resistance after infection with *Blumeria graminis* (57). Since Arabidopsis *snc1* autoimmunity requires AtEDS1 and ^Δsp^PbSTMI can impair this autoimmunity by impacting plant EDS1 levels, it is relevant to ask whether *snc1* autoimmunity confers resistance to other biotrophic or hemi-biotrophic pathogens and if PbSTMI will suppress this resistance. When infected with biotrophic *Erysiphe cichoracearum, snc1* showed slower fungal growth, shorter hyphal length, and decreased branching, with no difference in penetration resistance, compared to WT, ^Δsp^PbSTMI-GFP (Col-0) and ^Δsp^PbSTMI-GFP (*snc1*) lines **(Figure 6G-I).** However, the presence of ^Δsp^PbSTMI restored susceptibility in ^Δsp^PbSTMI-GFP (*snc1*) plants **(Figure 6G-I)**. Similarly, *snc1* plants showed resistance to *C. higginsianum*, but ^Δsp^PbSTMI-GFP (*snc1*) lines became susceptible, with the presence of necrotrophic secondary hyphae at 3 DPI (**Figure 6J,K**), highlighting the conserved role of PbSTMI in virulence beyond clubroot.

## 3. Discussion

Pathogens and host plants are locked in an ongoing arms race, with pathogens evolving effectors to manipulate host defenses and facilitate successful colonization. Some effectors are shared among related species, supporting their evolutionary success. This study identifies the effector PbSTMI from *P. brassicae*, which is conserved across Eumycota, including the hemibiotrophic ascomycetes *Colletotrichum* and *Pyricularia*, that infect both monocots and dicots. PbSTMI orthologs are also present in pathogens within Basidiomycetes, which are biotrophic, gall-inducing smut fungi. Recently, PbSTMI (PBTT_09143) and its paralogs were clustered with the oomycete *Albugo candida* secretome (58), underscoring their evolutionarily conserved role in virulence during infection. The four PbSTMI paralogs show differential expression during *P. brassicae* infection of Arabidopsis, with PbSTMI being most abundant, indicating tight regulation linked to different infection stages and disease progression. Their close genomic location suggests gene duplication and chromosomal rearrangements occurred during evolution.

Overexpressing ^Δsp^PbSTMI in Arabidopsis increased susceptibility to *P. brassicae* in roots as well as *C. higginsianum* in leaves. When treated with the flg22 peptide, we found that ^Δsp^PbSTMI overexpression does not affect flg22-triggered early PTI responses but does affect late immune responses. flg22-triggered PTI involves SA-related genes and proteins such as BAK1/BKK1, CPK5/6/11, and EDS1. The EDS1 protein level was significantly reduced in flg22-treated leaves of ^Δsp^PbSTMI-GFP (Col-0) lines compared with the typical EDS1 upregulation observed in WT Arabidopsis; this aligns with published findings on *EDS1* upregulation (44,59). Overexpression of EDS1 is known to confer disease resistance (60), prompting us to explore ^Δsp^PbSTMI’s role in EDS1-mediated immunity.

Selective yeast two-hybrid screening identified AtEDS1 as a potential interactor of PbSTMI, validated *in planta* through BiFC and co-IP. EDS1 localizes to both the nucleus and cytoplasm, with a balanced distribution essential for transcriptional reprogramming of many downstream NLR genes, as well as SA-regulatory and defense-related genes (42). Extranuclear interactions between ^Δsp^PbSTMI and AtEDS1, however, nuclear enrichment of complemented YFP signals observed after MG132 treatment further indicates that ^Δsp^PbSTMI disrupts the nucleo-cytoplasmic balance of AtEDS1. The differential nucleo-cytoplasmic localization of both fluorescent-tagged ^Δsp^PbSTMI and AtEDS1 in plant cells suggests that ^Δsp^PbSTMI may influence AtEDS1 function by disrupting its balanced nucleo-cytoplasmic gradient. Previous studies show that effectors, AvrRps4, HopA1 and PcAvh103, target EDS1 and impair immune responses and disease resistance, potentially by disrupting its interactions with intracellular TIR-NLRs (40) or by disrupting the EDS1-PAD4 interaction essential for EDS1-mediated immune signaling (41). EDS1 is a critical regulator of plant immunity that acts as a common hub and target for pathogen virulence effectors, which are protected by sensor, TIR-type NLR receptors.

In plants, TIR-NLR immunity relies on EDS1-PAD4/SAG101-hNLR modules for resistosome assembly and disease resistance (50,61). NLRs AT5G47260 and ADR1-L3 provide TIR-NLR resistance to clubroot in Arabidopsis Bur-0 (7). The sensor TIR-NLR, SNC1, monitors ADR1-L1/L2 and triggers resistance when interacting with the pathogen effector AvrPtoB (28,29). Auto-active gain-of-function (GOF) TIR-NLR mutants constitutively activate defense pathways (47,48,62), and knockout mutations of EDS1-PAD4/SAG101-hNLR components alleviate autoimmune responses in TIR-NLR GOF mutants like *snc1*, *chs3-1*, and *chs3-2D* (47,48,62). Overexpressing ^Δsp^PbSTMI rescues *snc1* and *chs3-1* autoimmunity by decreasing AtEDS1 protein levels, and the increased susceptibility of ^Δsp^PbSTMI-GFP (*snc1*) lines to both biotrophic and hemibiotrophic pathogens confirms ^Δsp^PbSTMI’s role in suppressing plant immunity. No significant effect was observed in ^Δsp^PbSTMI-GFP (*msl10-3G*) lines, in which autoimmunity is EDS1-independent and involves Ca^2+^ surges from mechanosensitive responses (55,56).

The balance between defense and growth is carefully maintained by the delicate regulation of abundance and sensitivity of both PRR and NLR receptors. Overexpression or autoimmune activation of these receptors often leads to excessive defense responses, with a penalty on plant growth. Many *modifiers of snc1* (*MOS*) genes have been identified and function in the nuclear-cytosolic shuttling of the NLR receptor SNC1, aiding in the regulation of gene expression related to plant immunity (63). Negative regulation of the sensor TIR-NLR SNC1 is mediated via the 26S proteasome, which is controlled by CPR1/CPR30, an F-box protein in the SCF (Skp1-cullin-F-box) complex (54). Our results showed that co-expressing SNC1 and ^Δsp^PbSTMI, with or without MG132, did not significantly alter SNC1 abundance, indicating that the suppression of *snc1* autoimmunity mediated by ^Δsp^PbSTMI does not target receptor turnover but instead affects signaling pathways downstream of SNC1.

A basal EDS1 level is regulated by Cullin3-based E3 ligase adaptors NPR3 and NPR4 (44), which act as transcriptional repressors and SA receptors, promoting EDS1 degradation through the 26S proteasome, while PBS3 protects EDS1 from degradation (43,44). Inhibition of the 26S proteasome with MG132 restores AtEDS1 levels in *N. benthamiana* leaves co-infiltrated with AtEDS1-GFP/^Δsp^PbSTMI-mRFP suggesting that ^Δsp^PbSTMI-triggered AtEDS1 turnover occurs through proteasome-mediated degradation. It remains unclear whether ^Δsp^PbSTMI directly recruits AtEDS1 to the proteasome or causes the loss of PBS3-mediated stabilization, leading to its degradation. Our results indicate that the interaction between ^Δsp^PbSTMI-AtEDS1 and subsequent AtEDS1 turnover negatively impacts the formation of EDS1-PAD4/SAG101-hNLR complex. This suppression of TIR-NLR-mediated immunity occurs by disrupting downstream resistosome assembly via changes in AtEDS1 abundance, rather than by directly interfering with oligomerization.

SA has a critical role in plant resistance to both biotrophic (64–66) and hemibiotrophic (67,68) pathogens. In turn, EDS1 is a master regulator in both SA and TIR-NLR-mediated immunity (34,36,45). This study confirmed elevated SA levels in the autoimmune *snc1* mutant (47,49). Overexpression of PbSTMI reduced SA levels *in planta* and transcription of SA-responsive defense genes (*PR1*, *PR2*, *PR5*), as well as SA biosynthesis and transport genes (*ICS1*, *EDS5*), resulting in decreased SA-mediated immunity.

EDS1-dependent pathway components are vital for TIR-NLR-mediated ETI, influencing SA accumulation, resistance gene responsiveness, basal resistance, and PTI responses (37,45). PTI and TIR-NLR-mediated ETI converge in Arabidopsis, with EDS1 working with LRR receptors to boost resistance (37,38). The aphid effector Mp10 suppresses PTI and triggers ETI via EDS1 by destabilizing receptors like FLS2 and FER, rendering plants insensitive to PAMPs (69,70). Loss of the receptor kinases BAK1 and BKK1 in *bak1* and *bkk1* mutants can activate NLR-dependent ETI through EDS1 (51,71). Our findings, showing PTI responses, increased EDS1 levels and upregulated SA genes in WT, support PTI and ETI convergence in resistance (44,72). The TIR-NLR GOF mutant *chs3-1*, in collaboration with CSA1 (62), exhibits elevated apoplastic ROS levels, increased callose deposition, and MAPK activation, leading to spontaneous cell death. Expression of PbSTMI in *chs3-1* reduces those TIR-NLR-triggered responses, preventing MAPK activation in *chs3-1* lines but not in wild-type plants after flg22 treatment, indicating it is not involved in PAMP perception or MAPK relay. Instead, PbSTMI disrupts TIR-NLR-mediated ETI pathways and suppresses TIR-NLR-triggered responses via EDS1-dependent signaling.

Similar to previous studies on pathogen effectors that form homo-oligomers (73), PbSTMI can exist in both monomeric and oligomeric forms. Furthermore, PbSTMI can undergo both homo-and hetero-oligomerization. The two N-terminal conserved motifs, ^29^DxKxDDK^35^ and ^132^VxxFQxAP^139^, are essential for this oligomerization and likely serve as the binding interface. Hetero-oligomerization between PbSTMI and PbSTMI-L3 suggests that oligomerization extends across the family and, as a key regulator of PbSTMI activity, provides insight into how *P. brassicae* effectors could coordinate host immune suppression with oligomerization preferences among paralogs, possibly fine-tuning effector activity and the spatio-temporal regulation of immune responses during different stages of clubroot infection.

Our findings identify PbSTMI as an effector that modulates TIR-NLR plant immunity in an EDS1-dependent manner **(Figure 7**). Understanding how PbSTMI regulates TIR-NLR signaling provides insight into how pathogens hinder broad-spectrum plant immunity. The presence of this effector family across various plant pathogens suggests that additional functions may arise from paralogs or orthologs. Gaining insight into the functional diversification of this effector family will be crucial for understanding how new, more virulent pathotypes with recently evolved effector repertoires emerge, thereby enabling the development of novel strategies for robust, durable plant resistance.

**FIGURE 7:**
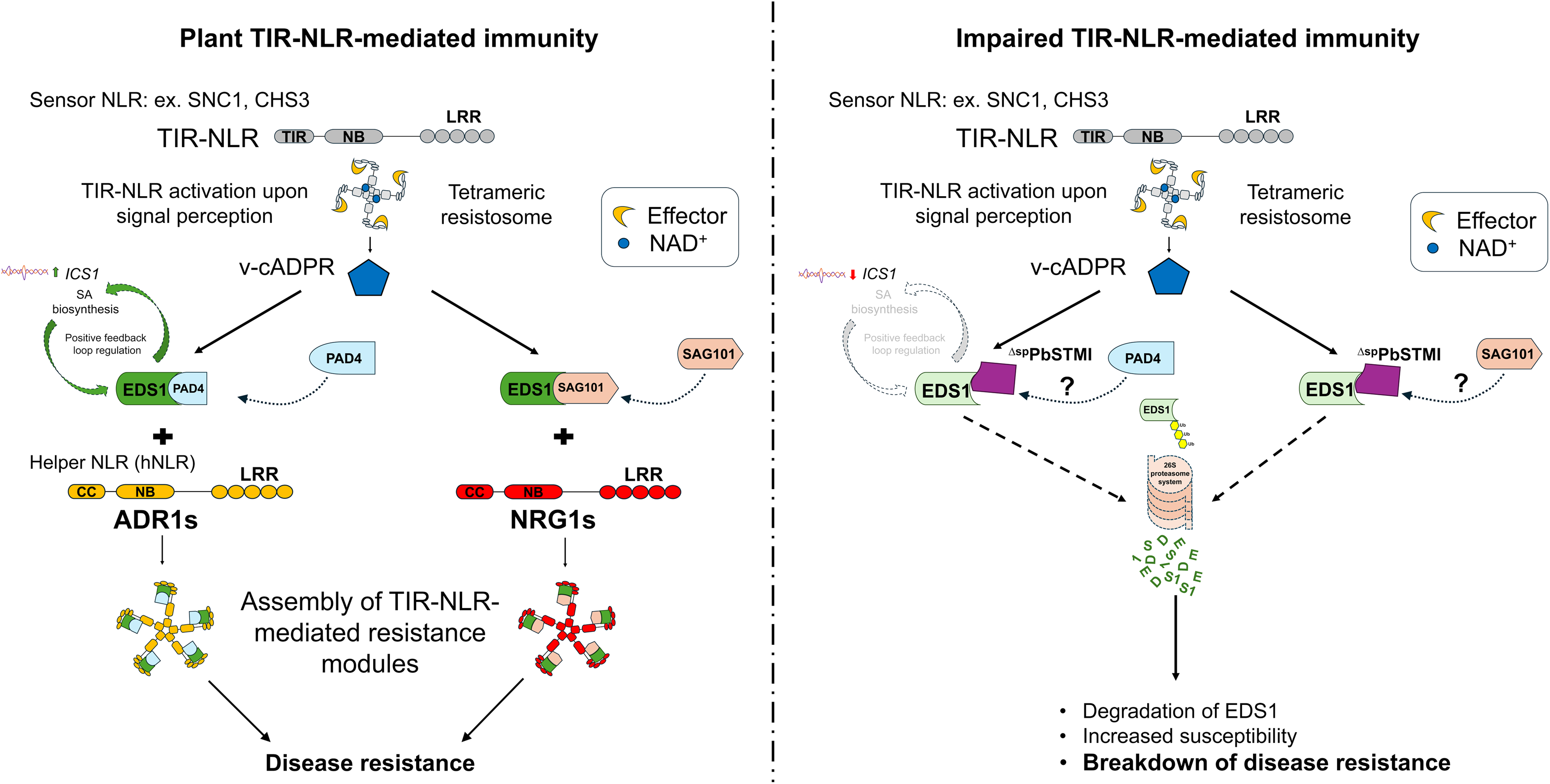
Proposed model of ^Δsp^PbSTMI-mediated impairment of TIR-NLR immunity in plants. In TIR-NLR-mediated plant immunity, TIR-NLRs recognize pathogen effectors and transduce signals via EDS1, PAD, and SAG101, activating defenses (20,21) through Helper NLRs like ADR1 and NRG1 (20,23,25). In ^Δsp^PbSTMI-mediated impairment of TIR-NLR immunity, ^Δsp^PbSTMI promotes EDS1 degradation via the 26S proteasome, thereby regulating downstream EDS1 oligomerization, which is necessary for the modular assembly of the resistosome and subsequent defense signaling for disease resistance. The ^Δsp^PbSTMI-mediated impairment of TIR-NLR immunity, along with downregulation of EDS1-regulated SA biosynthesis and transcription of defense regulatory genes, increases plant susceptibility to diverse pathogen infection.

## 4. Materials and Methods

### 4.1. Plant materials and growth conditions

*Arabidopsis thaliana* Col-0 and mutant plants were grown in Sunshine Mix #3 soil (Sun Gro Horticulture Inc., Vancouver, BC, Canada) under controlled conditions: 22 °C, 16 hours of light and 8 hours of darkness, with a light intensity of 100 µmol photons m^-2^ sec^-1^ in a growth chamber (Conviron E8, Winnipeg, MB, Canada): *Nicotiana benthamiana*, and *Brassica napus* cv. Westar (canola) was also grown in the same soil after stratification for 2 days at 4 °C. Seedlings were transplanted into pots 10 days post-germination and then grown at 25 °C under a 16 hour light and 8 hour dark cycle, with a light intensity of 160 µmol photons m^-2^ sec^-1^. To select transgenic plants, seeds were sterilized with 70% ethanol for 2 minutes and 50% bleach on a rolling rotor for 10 minutes followed by incubation at 4°C for two days and sowing on plates containing ½ Murashige and Skoog (MS, PhytoTech, Kansas, USA) salts, 3% sucrose, and 50 mg/mL hygromycin (PhytoTech, Kansas, USA).

### 4.2. Strains, culture media, and growth conditions

*Agrobacterium tumefaciens* GV3101 and *Escherichia coli* DH5α and Top10 cells were cultured at 28°C and 37°C, respectively, in Luria-Bertani (LB) medium supplemented with the appropriate antibiotics. *E. coli* was transformed using a heat shock method (74), while *A. tumefaciens* transformations were performed using an ECM 399 electroporator (BTX, Harvard Bioscience Inc., city, state, USA) under standardized electroporation conditions (75). The *Saccharomyces cerevisiae* strain YTK12, for the signal peptide (SP) validation assay, was grown on yeast extract-peptone-dextrose (YPD) media. For effective selection, yeast cultures were cultivated on CMD-W and YPRAA media supplemented with antimycin A (76), and transformations were carried out using the established LiAc/SS carrier DNA/PEG method (77).

### 4.3. Pathogen inoculation and infection assay

*P. brassicae* pathotype-3 was obtained from Dr. Gary Peng (AAFC Saskatoon Research Centre) and propagated through gall production in *B. napus* cv. Westar (canola) plants. Resting spores were extracted from air-dried canola root galls in water and quantified using a hemocytometer. For routine pathogen inoculations, fourteen day-old Arabidopsis seedlings were inoculated at the base of their stems with 500 µL of a solution containing 4 × 10^7^ *P. brassicae* resting spores/mL in water. To determine the resistance/susceptibility of various Arabidopsis genotypes to *P. brassicae* infection, a larger volume of the resting spore solution at the same concentration was used. Control plants received 500 µL (or more) of distilled water under the same conditions. In both cases, three independent biological replicates were carried out.

The Arabidopsis-adapted powdery mildew fungus *Erysiphe cichoracearum* was maintained and propagated on host cucumber (*Cucumis sativus*, variety Sweet Slice; McKenzie) plants in the lab (YW). Three-to-four-week-old Arabidopsis plants were inoculated with conidiospores at a density of 5 to 10 conidia mm^-2^. Inoculated leaves at the indicated time points post-inoculation were detached, fixed, and stained as previously described (78).

For disease assays with the adapted anthracnose pathogen *Colletotrichum higginsianum*, three-to-four week-old Arabidopsis plants were sprayed with conidial suspensions or spotted with droplets and immediately placed in a 100% humidity chamber. Inoculated plants were photographed at different time points post-inoculation to document the infection time course, and infected leaves were detached, fixed, and stained with trypan blue (79).

### 4.4. Pathogen abundance assay

To quantify *P. brassicae* abundance, Arabidopsis roots from infected plants were harvested at the indicated times post-inoculation and DNA extracted using an E.Z.N.A plant DNA kit (Omega Bio-tek, Georgia, USA), following the manufacturer’s protocol. Biomass was quantified by PCR of *P. brassicae INTERNAL TRANSCRIBED SPACER 1* (*ITS1*), normalized to the Arabidopsis *ACTIN7* (*AtACT7*) gene. All primers used in this study are listed in **Table S3**. Nile red staining of lipid droplets was also used to visualize, via confocal microscopy, the extent of infection, the different life stages of *P. brassicae*, and associated changes in host tissues.

### 4.5. RNA extraction, cDNA synthesis, and relative expression analysis

*PbSTMI* paralog expression was determined through RT-PCR of root tissue samples collected from infected Arabidopsis plants at 0, 2, 5, 7, 14, 21, and 28 days post-inoculation (DPI) as well as from resting spores from 35 day-old canola galls. Total RNA was extracted using a RNeasy mini kit (Qiagen, Hilden, Germany) and cDNAs were synthesized with the QuantiTect Reverse Transcription Kit (Qiagen, Hilden, Germany). RT-PCR quantitative expression data were quantified and analyzed using FIJI ImageJ software (https://imagej.net/Fiji). The relative expression of *PbSTMI* was measured against the internal control *PbRPS17* from *P. brassicae*, with three independent biological replicates per analysis. All primers used in the expression profiling are listed in **Table S3**.

### 4.6. Bioinformatic analysis and protein modelling

PbSTMI homologs were identified through BlastP (https://blast.ncbi.nlm.nih.gov/Blast.cgi?PAGE=Proteins) searches against the non-redundant (nr) database in NCBI (https://blast.ncbi.nlm.nih.gov/Blast.cgi) and multiple sequence alignments (MSAs) of the full-length protein sequences were generated using Clustal Omega (https://www.ebi.ac.uk/Tools/msa/clustalo/). MSAs were visualized using the ESPrint 3.0 server (https://espript.ibcp.fr/ESPript/ESPript/). A phylogenetic tree was generated by Clustal Omega (https://www.ebi.ac.uk/jdispatcher/msa/clustalo) from the MSAs and visualized using ITOL (https://itol.embl.de/). PbSTMI was analysed for functional domains using the Conserved Domain Database (CDD: https://www.ncbi.nlm.nih.gov/Structure/cdd/wrpsb.cgi) and conserved motifs from PbSTMI-homologs were identified by MEME scan (https://meme-suite.org/meme/tools/meme) with default settings. Three-dimensional protein models of PbSTMI paralogs and representative orthologs were generated using AlphaFold 2.0 at Neurosnap (https://neurosnap.ai/). The protein model for each effector, based on the highest mean pLDDT score, was selected from five generated models. The protein individual/superimposed models were visualized using UCSF Chimera (https://www.cgl.ucsf.edu/chimera/).

### 4.7. Signal peptide validation using a yeast signal sequence trap assay

Signal peptides (SP) at the N-termini of full-length PbSTMI homologs were predicted using SignalP 5.0 (https://services.healthtech.dtu.dk/services/SignalP-5.0/), which was later used to create versions of the constructs coding for proteins lacking SP. Functionality of the predicted signal peptide was assessed in yeast YTK12 lacking an active invertase secretory system (80). EcoRI and XhoI restriction sites were added to the amplified PbSTMI SP coding sequence for insertion into the pSUC2 vector, in frame with the SP-deficient invertase gene. The SP^PbSTMI^-pSUC2 construct was used to transform YTK12, and positive clones were selected on CMD-W media and confirmed on YPRAA selection media (1) and the 2,3,5-triphenyltetrazolium chloride (TTC)-colorimetric assay for active invertase secretion. Invertase secretion was indicated by the reduction of TTC to red 1,3,5 triphenyl formazan (TPF) with AtLCR78 as a positive control (1,2).

### 4.8. Generation of *PbSTMI* gene-vector constructs

The PbSTMI coding sequence without the SP (^Δsp^) was amplified from cDNA and cloned into plant expression binary vectors using Gateway cloning technology (Thermo Fisher Scientific, Waltham, MA, USA), which was later used to generate Arabidopsis overexpression (OE) lines. All *PbSTMI* constructs and paralog vector constructs were generated after removing the predicted signal peptide sequences. The vectors included pH7FWG2, with a GFP tag, and pH7WG2, without a GFP tag. To generate the PbSTMI^Δ^(^19–37^)-GFP construct, the N-terminal end of PbSTMI was deleted using appropriate primers (**Table** S**3**), followed by the addition of N-terminal AttB1 and C-terminal AttB2 recombination sites for cloning into pDONR207 and subsequently shuttled into pH7FWG2 via Gateway cloning (Thermo Fisher Scientific, Waltham, MA, USA). DNA sequencing (Eurofins, Toronto, ON, Canada) and RT-PCR were conducted to confirm the targeted mutations. Coding sequences of the selected *PbSTMI-*orthologs, without the predicted SP, were commercially synthesized by IDT (Coralville, Iowa, USA), cloned into pDONR207, and shuttled into pH7FWG2 for use in subcellular localization of the protein orthologs. A GFP-expression construct, pH7WG2-GFP, was generated and used as a control. mCherry-tagged ER marker gene constructs, in the pBIN20 binary vector backbone (CD3-959), were purchased from the Arabidopsis Biological Resource Centre (https://abrc.osu.edu/). For co-localization studies, expression constructs were created using the Gateway cloning kit, with the Arabidopsis *EDS1* (AT3G48090.1) sequence inserted into both C-terminal GFP-tagged, pH7FWG2 and mRFP-tagged, pH7RWG2 via an LR reaction.

### 4.9. Subcellular localizations using confocal microscopy

Transient expression of PbSTMI was achieved by transforming *A. tumefaciens* GV3101 with the appropriate gene vector construct and selecting positive transformants on LB-spectinomycin (100 mg/L), kanamycin (50 mg/L), or rifampicin (50 mg/L) plates. The localization of ^Δsp^PbSTMI-GFP was assessed through transient expression in five-week-old *N. benthamiana* leaves. A 500 µL injection of *A. tumefaciens* GV3101 (OD_600_ of 0.3) was administered to the abaxial side of the leaves (81). Localization was observed 2-3 days later using a confocal laser scanning microscope (LSM880 with Airyscan; ZEISS) at 488/500-530 nm for GFP fluorescence (81). Z-stack and time-lapse techniques were used to analyze the distribution and movement of ^Δsp^PbSTMI-GFP in epidermal cells. All fluorescing cells, including those with weak signals, were analyzed using ImageJ (https://imagej.net/Fiji).

### 4.10. Yeast-2-Hybrid interactions

To perform Y2H screening, the *PbSTMI* coding sequence lacking a signal peptide was cloned into the pGBKT7-GW vector and used to transform the Y2H Gold strain as a bait to screen for potential plant targets. To confirm potential targets of PbSTMI, the coding sequences of potential Arabidopsis protein candidates were inserted into the prey vector pGADT7-GW, and the various plasmid combinations were co-transformed into the Y2H Gold strain. Positive clones were selected on double dropout (SD/-Leu/-Trp) and quadruple dropout media with X-α-Gal (SD/-Ade/-His/-Leu/-Trp with 30 µg/mL working concentration of X-α-Gal) plates. SV40 T-antigen (T) was used as a positive control (with pGBKT7-53) while pGBKT7-Lam served as a negative control.

### 4.11. Bimolecular fluorescence complementation (BiFC) assay

Genes were cloned into gateway-compatible vectors pGTQL1211YN for the nYFP-tagged fusion protein and pGTQL1221YC for the cYFP-tagged fusion protein and subsequently used to transform *A. tumefaciens* (GV3101). Positively transformed Agrobacterium were cultured, and an equal ratio (1:1) of both clones was resuspended in infiltration buffer (10 mM MES, 2% (w/v) glucose, 10 mM MgCl₂, 200 μM acetosyringone) to a final concentration of OD_600_ 0.6. Five-week-old *N. benthamiana* leaves were infiltrated with combinations of Agrobacterium clones to assess YFP fluorescence complementation via transient expression assays. After 2 days, fluorescence was observed using a Zeiss 880 laser-scanning confocal microscope, and the images were analyzed in Fiji ImageJ.

### 4.12. GUS assay

Arabidopsis with a reporter gene containing the promoter region of the Arabidopsis PR gene *β-1,3-GLUCANASE* (*BGL2*) and the coding sequence of β-GLUCURONIDASE (GUS) in the *snc1* background was used to monitor *BGL2* promoter-driven constitutive GUS activity (47,53,82). *snc1* was transformed with a reporter gene containing the promoter region of an Arabidopsis PR gene linked to β-1,3-GLUCANASE (BGL2) and the coding sequence of β-GLUCURONIDASE (GUS) (82). *BGL2::GUS* expression was determined by staining leaf tissue for GUS activity after vacuum infiltration with X-Gluc solution following the procedure as described previously (82).

### 4.13. MG132 treatment

For the proteasome inhibition assay, 100 µM MG132 was prepared in water from a 10 mM DMSO stock and syringe-infiltrated into the abaxial side of five week-old *N. benthamiana* leaves 24 hours after infiltration with the gene constructs. Leaf samples were collected 20 hours after treatment or for fluorescence imaging under confocal microscopy. Control samples were infiltrated with a mock solution.

### 4.14. ROS detection assay

Leaves from four week-old Arabidopsis plants treated with 1 µM flg22 were collected early (5, 30 and 120 mins) and late (18 hours)-post-infiltration and stained with DAB to detect H_2_O_2_, an indicator of ROS activity (83). Freshly collected leaves were vacuum-infiltrated for 5 minutes with a freshly prepared DAB (Sigma-Aldrich) staining solution (1 mg/ml DAB, 10 mM Na_2_HPO_4_, 0.1% Tween 20) and incubated overnight at 30^°^C in the dark. Samples were decolorized in a fixation solution (methanol: chloroform: acetic acid = 6: 3: 1) and H_2_O_2_ was visualized as a reddish-brown color in the leaves.

### 4.15. Callose deposition assay

To assess callose deposition, approximately 100 µL of 1 µM flg22 was abaxially infiltrated into the symmetric half of an Arabidopsis leaf, and the other half was infiltrated with autoclaved deionized water as a mock treatment. Leaves were harvested 18 hours after treatment. Three Arabidopsis plants with at least 2-3 leaves per plant were used for each treatment. Leaves were bleached and fixed in methanol: chloroform: acetic acid = 6: 3: 1 for 1-2 days, followed by rehydration through an ethanol gradient and washed with water to remove residual ethanol. Processed leaf samples were stained overnight in 0.05% aniline blue diluted in 150 mM K_2_HPO_4_, pH 9.5, followed by washing in 150 mM K_2_HPO_4,_ pH 9.5. Samples were mounted in 30% glycerol and observed under an Axioplan epifluorescence microscope (Carl Zeiss Canada, Toronto, CA) with UV application. Multiple Z-stacks encompassing surface-to-midplane images were taken from multiple cells for each treatment.

### 4.16. Protein extraction and western blot analysis

Quick-frozen plant tissue was ground to a fine powder in liquid nitrogen using a mortar and pestle. Protein was extracted using extraction buffer [50 mM Tris, pH 7.5; 150 mM NaCl; 10% glycerol; 2 mM EDTA; 5mM DTT, 0.1% (v/v) Triton X-100] supplemented with 2 tablets of Complete Protease Inhibitor Cocktail (Roche; Cat. A32955) per 50 mL on ice for 30 minutes. The resulting mixture was centrifuged for 15 minutes at 20000x g at 4°C, and the protein concentration of the supernatant was determined spectrophotometrically using Bradford reagent (Sigma; Bradford, 1976). Equal aliquots of supernatant were boiled for 10 minutes in 2x Laemmli sample buffer supplemented with β-Mercaptoethanol (1:1) before separation through a 12% (w/v) SDS-PAGE gel. Gels were transferred to 0.2 µM nitrocellulose membranes (Bio-Rad, Mississauga, ON, CAD) for 1 hour at 100 V in a Mini-PROTEAN® Tetra Cell (Bio-Rad, Mississauga, ON, CAD). Membranes were blocked with 5% skim milk (BD Difco, Mississauga, ON, Canada) before probing with specific primary and corresponding HRP-conjugated secondary antibodies. Chemiluminescent horseradish peroxidase activity was visualized with Super Signal™ West Pico PLUS Chemiluminescent Substrate (Thermo Fisher Scientific, Waltham, MA, USA) and documented with a ChemiDoc™ Imaging System (Bio-Rad, Mississauga, ON, Canada). For PVDF transfer, membranes were additionally activated in methanol for 2-3 minutes and, after electrophoresis, incubated for 15 minutes in Ponceau S solution to stain for total proteins. A second 12% SDS-PAGE gel was stained with Coomassie blue R250 to show equal protein loading.

### 4.17. Co-Immunoprecipitation (Co-IP) assay

*N. benthamiana* leaves co-infiltrated with ^Δsp^PbSTMI-GFP and HA-tagged AtEDS1 were harvested 2 days after infiltration. Total protein was extracted using the method described above, and immunoprecipitation (IP) was conducted with 20 µl of ChromoTek GFP-Trap® Agarose beads following the manufacturer’s protocol (Proteintech, USA). Bound proteins were eluted by boiling for 10 minutes in 40 µl of 1X Laemmli sample buffer supplemented with 5% (v/v) β-mercaptoethanol. Eluted proteins were separated through 12% SDS–PAGE gels and immunoblotting with the indicated antibodies listed in **Table S4**. Ponceau S staining [0.1% (w/v) Ponceau S; 5% (v/v) acetic acid] of the membrane was used to show total loaded protein.

### 4.18. Blue-Native (BN) PAGE assay

BN-PAGE was performed using the NativePAGE™ Novex® Bis-Tris Gel System with precast NativePAGE™ 3 to 12% Bis-Tris Mini Protein Gels (10-well BN1001BOX). Electrophoresis was conducted in a Mini Gel Tank (Cat. A25977; Invitrogen) at 120 V for approximately 2 hours until the dye front reached the bottom of the gel. Electrophoresis was performed with Coomassie Blue G250 in a dark blue cathode buffer for about 30 minutes, then switched to a light blue cathode buffer for the remainder of the run. Proteins were transferred onto an activated PVDF membrane using a wet transfer for 2 hours at 110 V in a Mini-PROTEAN® Tetra Cell (Bio-Rad, Mississauga, ON, Canada). Subsequently, the membrane was blocked in 5% skim milk and incubated overnight at 4°C with an anti-EDS1 antibody, followed by incubation with the corresponding HRP-conjugated secondary antibody for 1 hour at room temperature, as described in the western blot section of the manuscript. The chemiluminescent signal was visualized using a ChemiDoc™ Imaging System (Bio-Rad, Mississauga, ON, Canada).

### 4.19. MAPK activation via phosphorylation

The abaxial side of leaves on four week-old Arabidopsis plants was infiltrated with 100-150 µL of 1 µM flg22 and harvested at 15, 30, and 60 mins post-treatment. Total protein was extracted, as described above. Total protein extracts were loaded into a 12% SDS-PAGE gel and separated using the Bio-Rad Mini Protean gel system. Proteins were blotted onto a 0.2 µm nitrocellulose membrane for immune detection using anti-phospho-p44/42 MAPK antibody (1:2000 dilution) after blocking the membrane for 1 hour in 5% BSA. Western ECL substrate solution was used for chemiluminescence detection of the target proteins after 1-hour incubation with anti-rabbit HRP-conjugated secondary antibody (1:1000 dilution) at room temperature. Coomassie blue staining of a second protein gel showed equal protein loading.

### 4.20. Salicylic acid quantification

Rosette leaves and root tissues of four week-old Arabidopsis wildtype (WT), *snc1,* and ^Δsp^PbSTMI-GFP (*snc1*) overexpression line #R1 plants were separately harvested and immediately frozen in liquid nitrogen. Extraction and quantification of salicylic acid (SA) were conducted at the NRC Aquatic and Crop Resource Development Research Center in Saskatoon, Canada, as described by Murmu *et al.* (2014)(84). Quantitative analysis of SA was performed by ultra-performance liquid chromatography-electrospray tandem mass spectrometry (UPLC/ESI–MS/MS) with data analyzed using MassLynx v4.1 (Waters Inc.) and QuanLynx v4.1 software (Waters Inc.). Three biological replicates were performed for each analysis.

## Author contributions

M.H., Y.W., P.C.B.S., and C.D.T. designed the research; M.H. performed all experiments; M.H., P.C.B.S., and C.D.T. analyzed the data; and M.H. wrote the manuscript with input from all authors. All authors read and approved the final manuscript.

## Acknowledgments

Funding was provided by SaskOilseeds, Western Grains Research Foundation, and Canola Council of Canada through the Canola Agronomic Research Program (2020.7 to CDT, PBS, YW), SaskOilseeds and Saskatchewan Agricultural Development Fund (20160138 to PBS, YW, CDT) and Natural Sciences and Engineering Research Council of Canada (RGPIN 04251-2018 to CDT).

## Data availability statement

Gene symbols and corresponding gene IDs for *Plasmodiophora brassicae* and other phytopathogens are listed in Table S1 and Table S2. The materials supporting the findings of this study can be obtained from the corresponding author upon reasonable request.

## Table of Contents

A secreted effector from the clubroot pathogen *Plasmodiophora brassicae*, PbSTMI, facilitates the proteasomal degradation of the master immuno-regulator EDS1, thereby inhibiting TIR-NLR–mediated and salicylic acid-dependent defense responses. The conservation of STMI-like proteins across kingdoms underscores an ancient immune evasion mechanism employed by gall-inducing pathogens to increase host susceptibility.

