## Supplemental Information for "A Cross-kingdom Effector Modulates EDS1-dependent TIR-NLR-mediated Plant Immunity"

**The PDF file includes:**

Figure S1 to S11

Data S1

**Other Supplementary information for this manuscript includes the following** (separate attachments)**:**

Table S1-S4

Movies S1

Supplementary Dataset 1

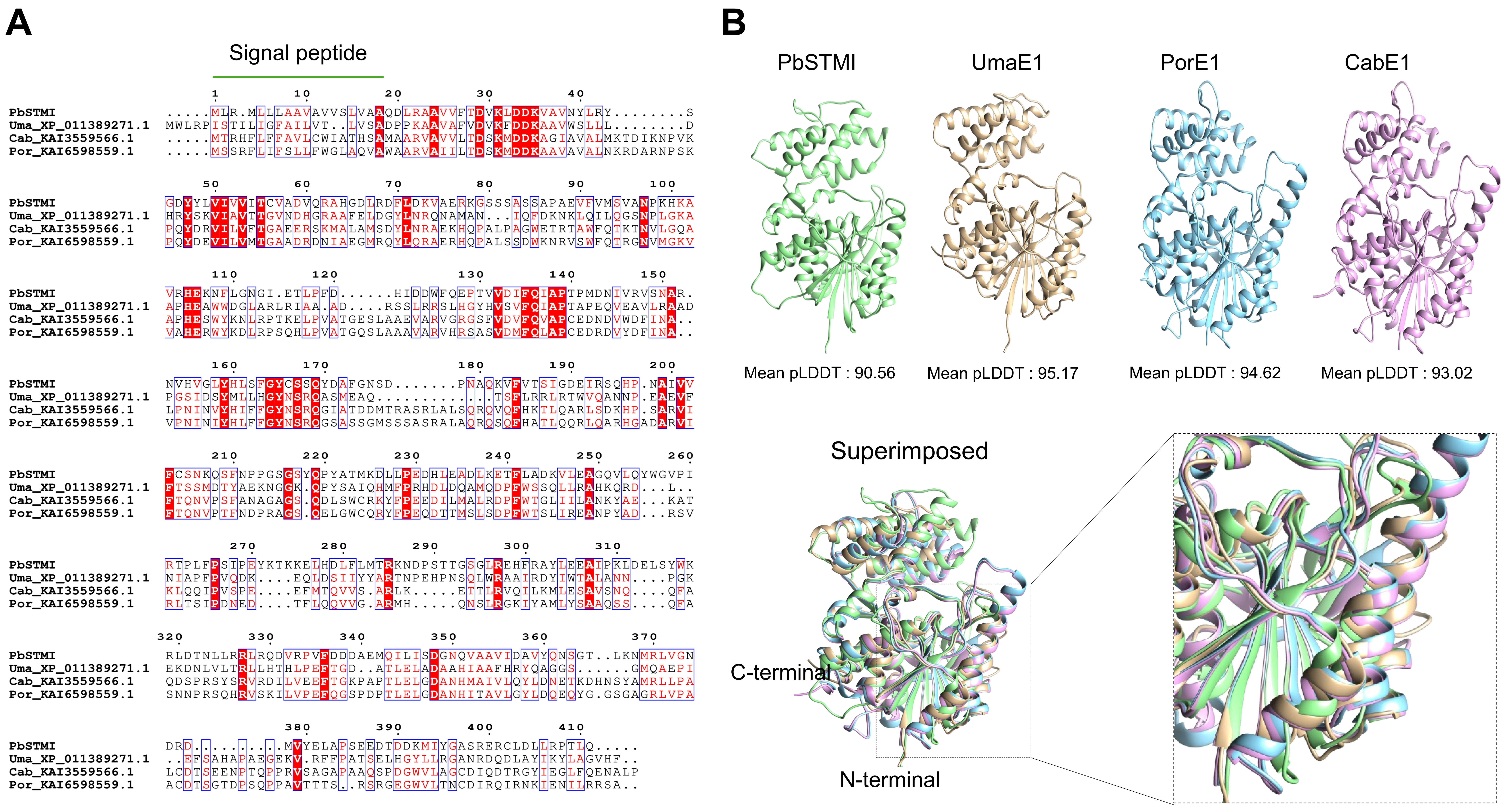


**Figure S1: Sequence and structural modelling of PbSTMI orthologs.** (A). Amino acid sequence alignment of PbSTMI and representative orthologs using Clustal Omega and visualized with ESPrint 3.0 server. Signal peptides were predicted using SignalP 5.0 and indicated with a green line. Conserved amino acids are highlighted in red, with conserved sequences boxed in blue and aligned with structural information from PbSTMI. (B). 3D models and superimposed images of PbSTMI (green), PorE1 (gold), UmaE1 (cyan), and CabE1 (magenta) generated using AlphaFold 2.0 at Neurosnap and visualized using UCSF Chimera. The best protein model for each effector, based on the highest mean pLDDT score, was selected from five generated models. UmaE1 - Uma_XP_011389271.1; PorE1 - Por_KAI6598559.1; CabE1 - Cab_KAI3559566.1.

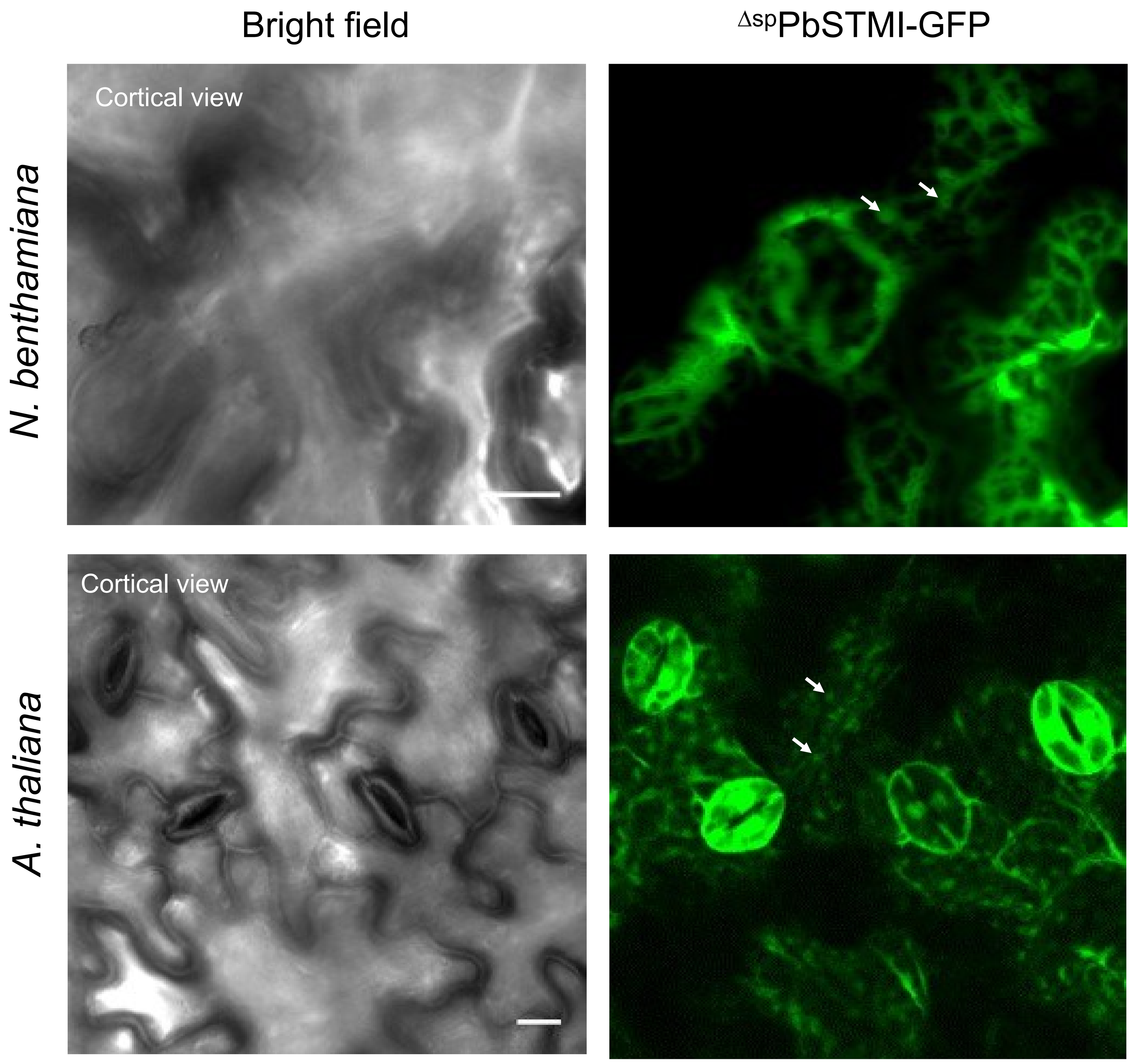


**Figure S2: Subcellular localization of $^{\Delta sp}$PbSTMI-GFP in plants.** The green channel shows the cortical view of the localization of $^{\Delta sp}$PbSTMI-GFP with cytosolic puncta (indicated by white arrows) in leaf epidermal cells. Scale bars = 10 µm.

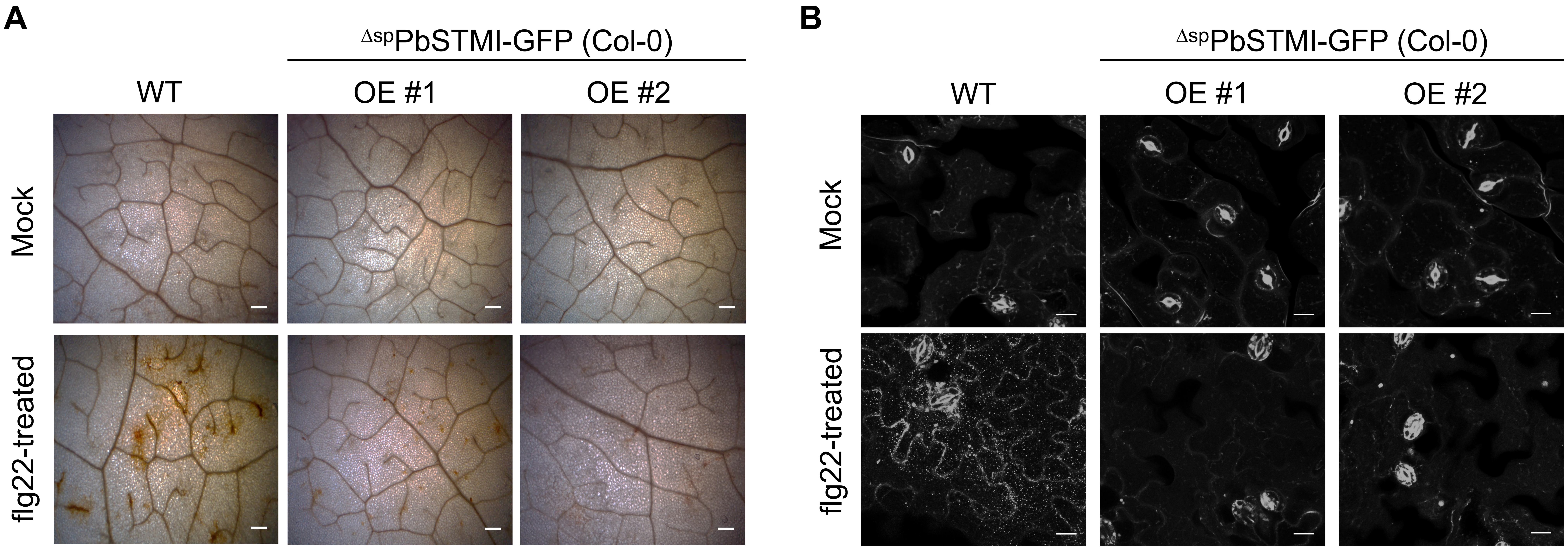


**Figure S3: $^{\Delta sp}$PbSTMI overexpression suppresses flg22-triggered late PTI responses in Arabidopsis.** (A). Leaves of four-week-old Arabidopsis treated with flg22 and assessed for ROS through DAB staining for $H_2O_2$ at 18 hours post-infiltration. Scale bars = 20 μm. (B). Rosette leaves of four week-old WT plants and $^{\Delta sp}$PbSTMI-GFP lines stained with aniline blue to show callose deposition 18 hours post-infiltration with 1 μM flg22. Callose was visualized under an epifluorescence microscope with UV illumination. Scale bars = 10 μm.

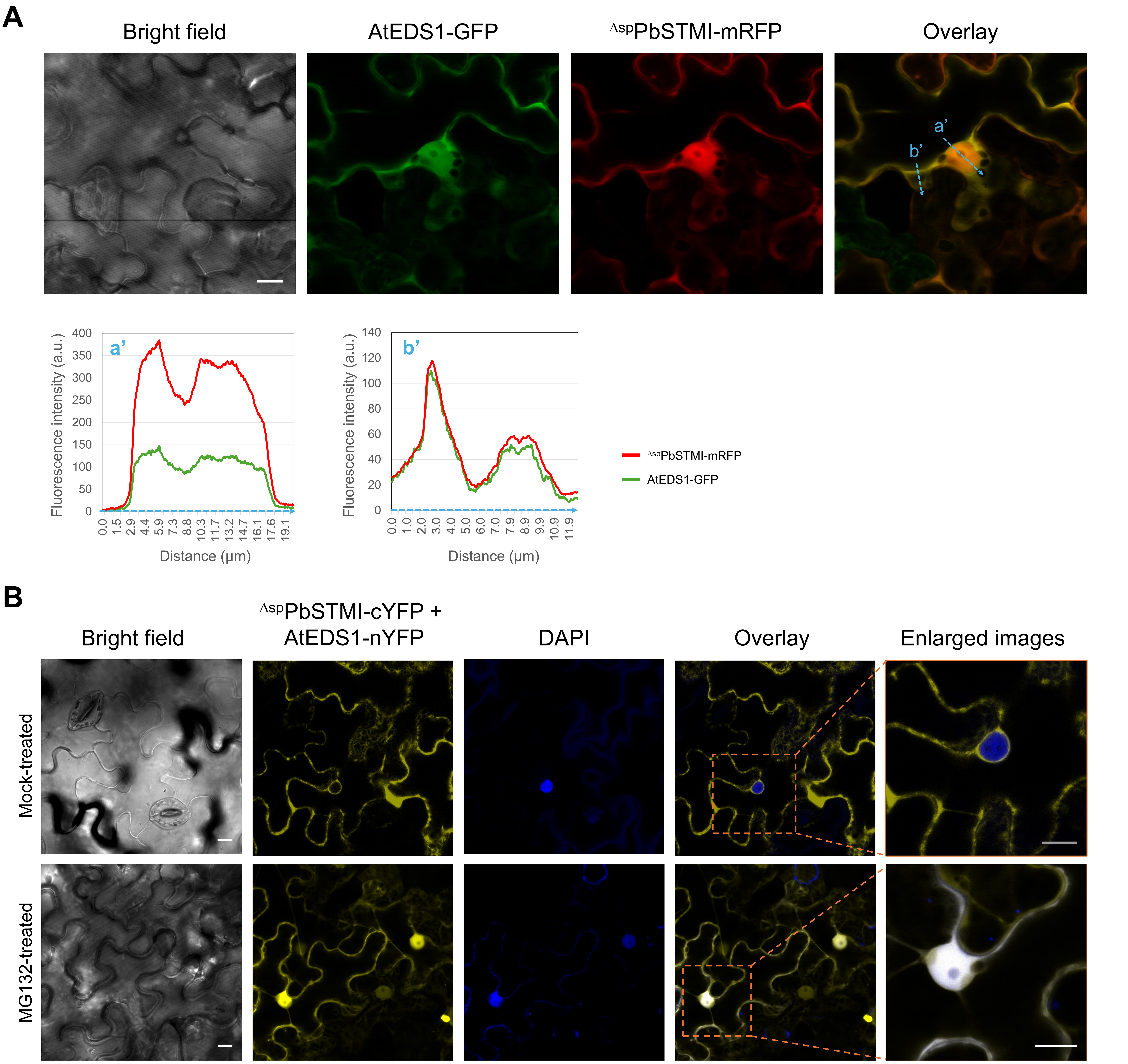


**Figure S4: Dynamics of the nucleo-cytoplasmic localization of $^{\Delta sp}$PbSTMI and AtEDS1 during transient expression in *N. benthamiana* leaves.** (A). Nuclear and cytoplasmic co-localization of $^{\Delta sp}$PbSTMI-mRFP and AtEDS1-GFP in *N. benthamiana*. Intensity plots show nuclear and cytosolic fluorescent enrichment of the overexpressed proteins. (B). The dynamic changes in nuclear BIFC interaction fluorescent signals with and without the proteasome inhibitor MG132. Scale bar = 10 μm.

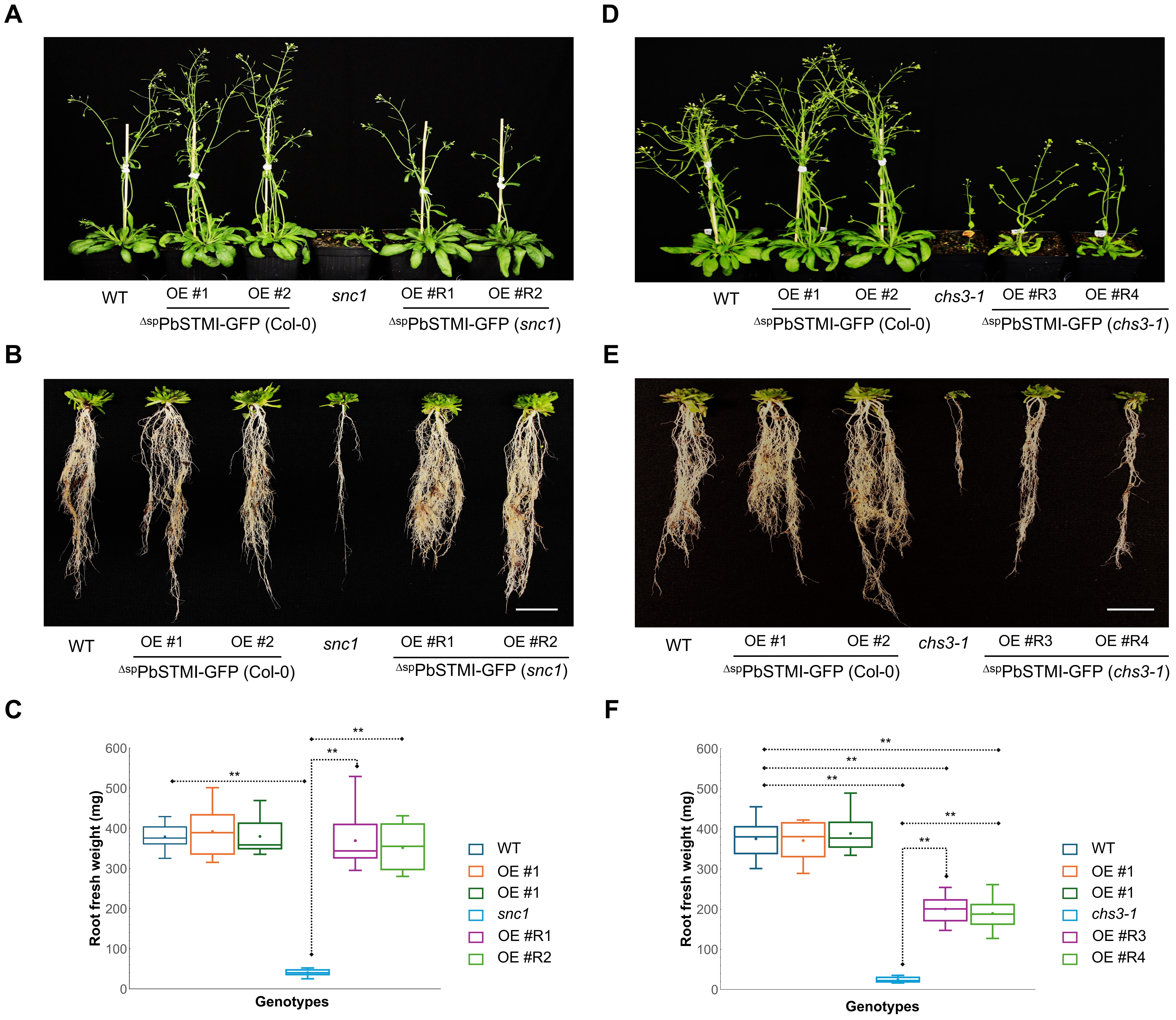


**Figure S5: $^{\Delta sp}$PbSTMI rescues *snc1* and *chs3-1* mediated autoimmunity in Arabidopsis.** Reproductive and root phenotypes of seven-week-old plants of the indicated genotypes (A, B, D, E). Scale bar = 2 cm. Quantified root fresh weight (mg) of the independent genotypes (C, F) is presented as colour-coded box plots where "x" denotes the mean value and the horizontal line in each box indicates the median of the data. n=10 individual plants. Statistical differences were assessed with one-way ANOVA followed by a post-hoc Tukey's HSD multiple comparison test. (*) and (**) indicate data points with significant differences at P value $< 0.05$ and $P < 0.01$, respectively.

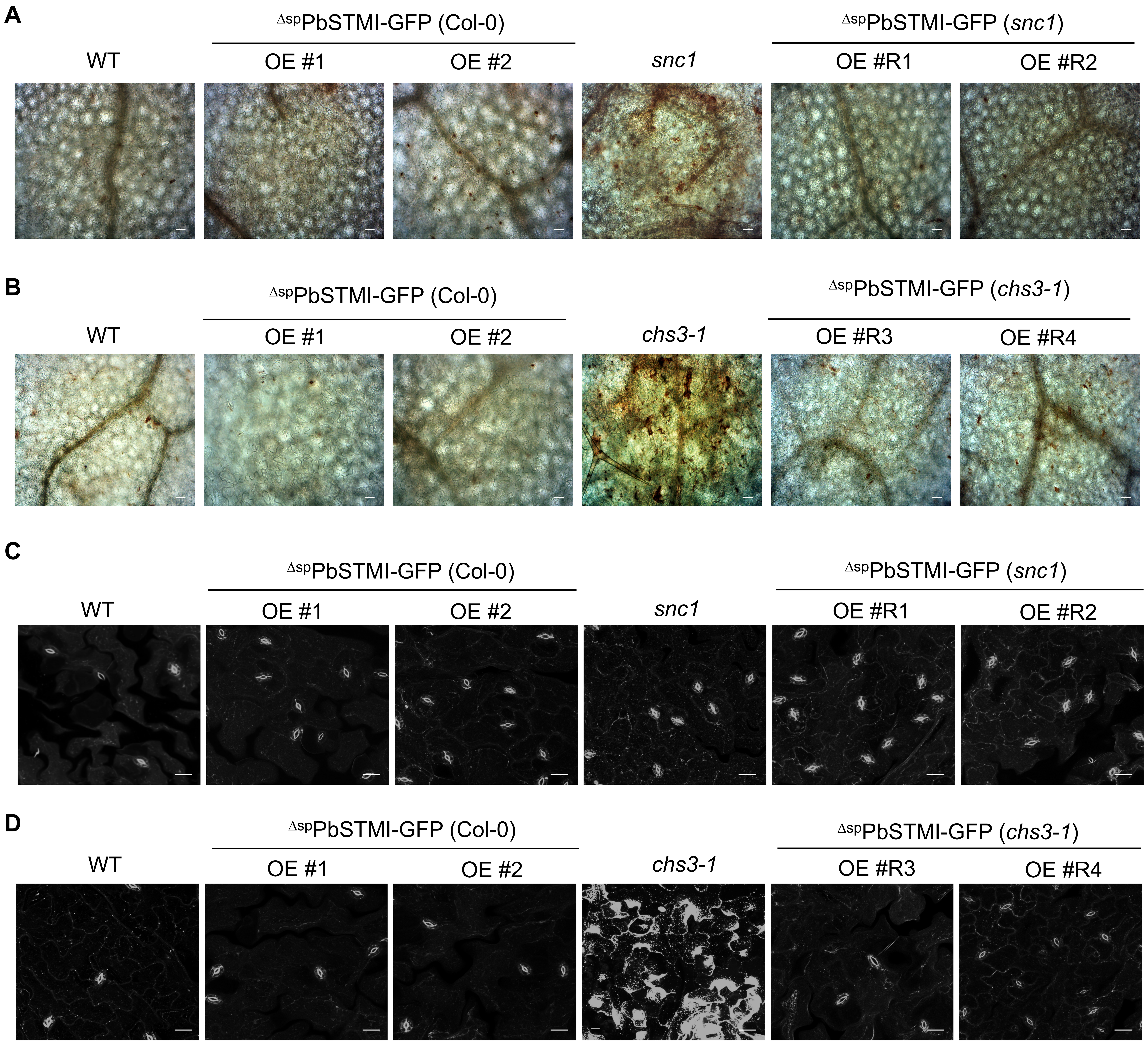


**Figure S6**. **$^{\Delta sp}$PbSTMI overexpression suppresses the TIR-NLR-mediated immune responses in Arabidopsis.** Detection of (A-B). $H_2O_2$ using DAB (Scale bars = 300 µm) and (C-D). callose using 0.05% basic aniline blue staining (Scale bars = 20 µm), in the leaves of three week-old WT, $^{\Delta sp}$PbSTMI-GFP (Col-0), snc*1,* $^{\Delta sp}$PbSTMI-GFP (*snc1*), *chs3-1*, and $^{\Delta sp}$PbSTMI-GFP (*chs3-1*) lines.

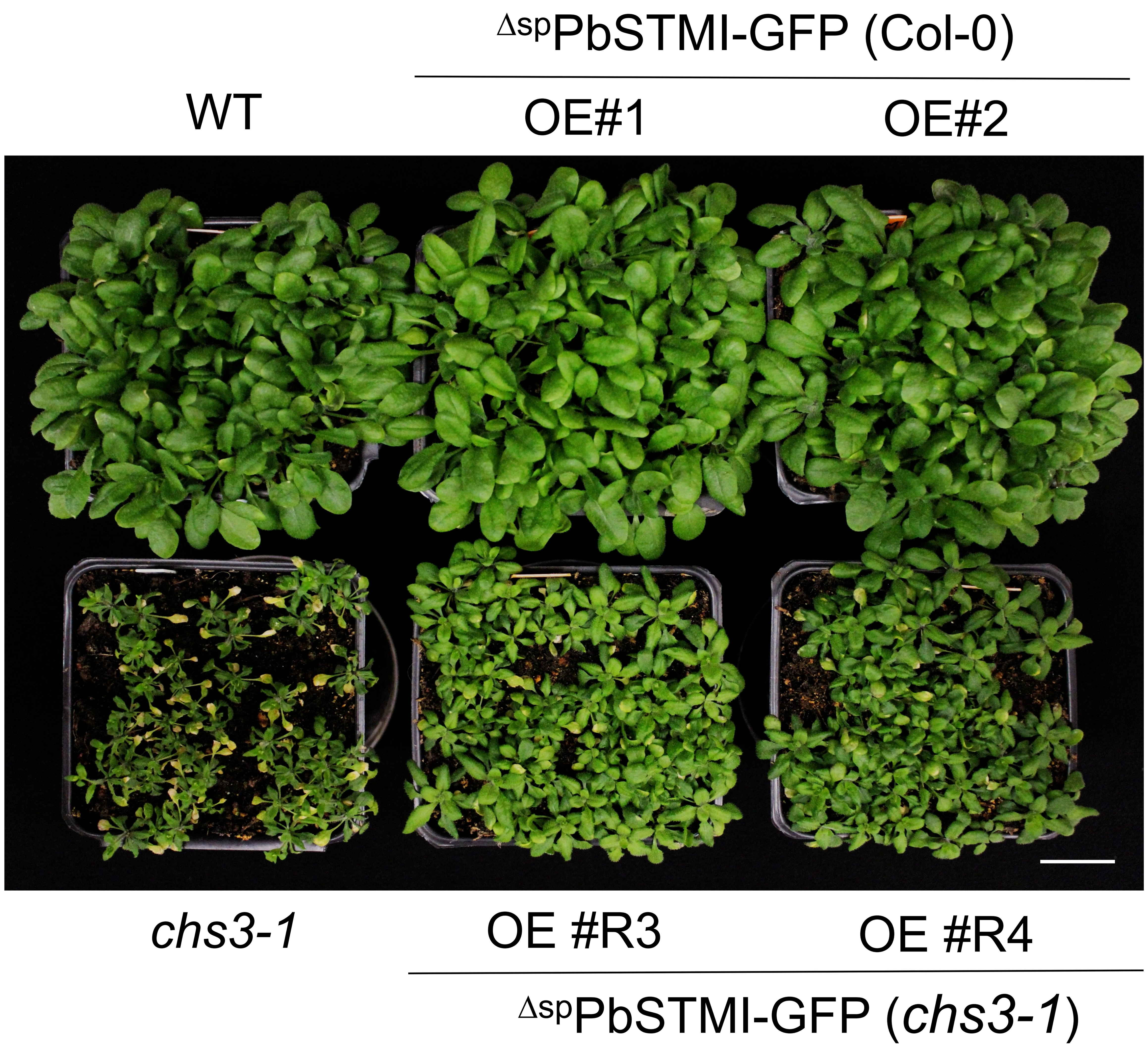


**Figure S7. Above-ground phenotypes of four week-old Arabidopsis WT, *chs3-1*, $^{\Delta sp}$PbSTMI-GFP (Col-0) and $^{\Delta sp}$PbSTMI-GFP (*chs3-1*) lines grown on soil.** Visible cell death phenotypes in *chs3-1* plants. Scale bar = 2 cm

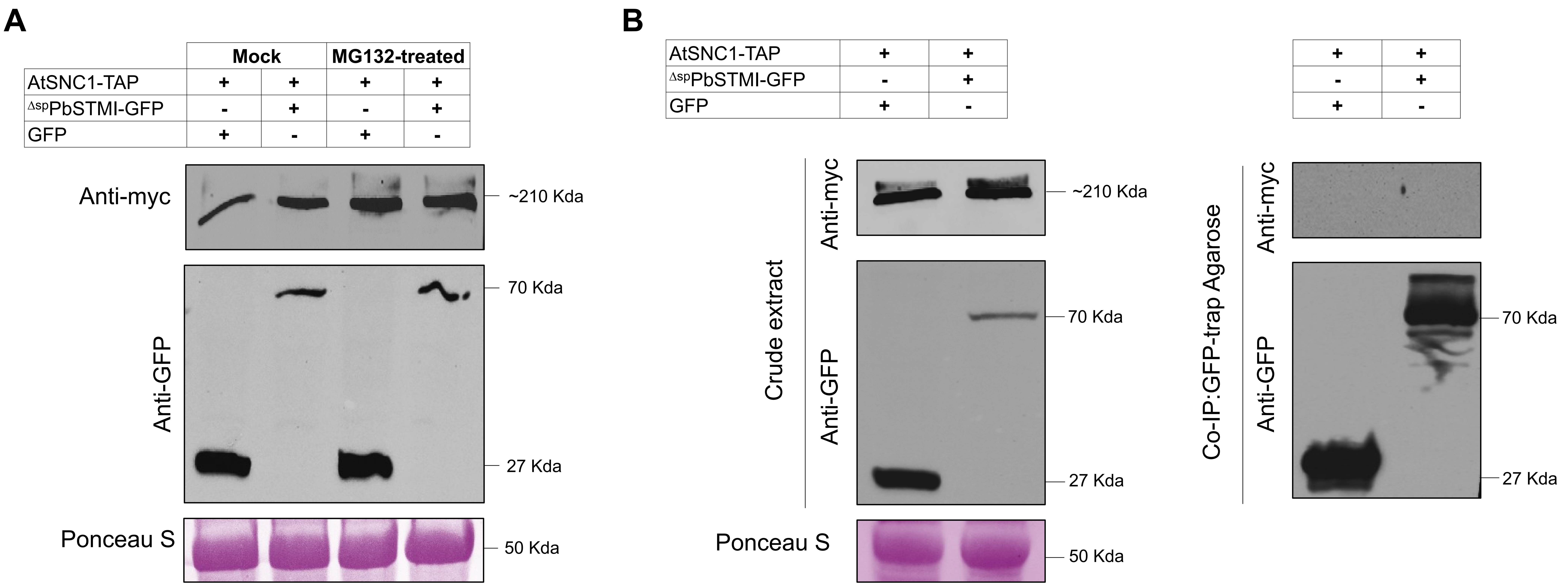


**Figure S8: $^{\Delta sp}$PbSTMI inhibits the downstream signaling of the overexpressing TIR-NLR receptor.** (A). Immunoblot detection of co-expressed proteins AtSNC1-TAP, along with $^{\Delta sp}$PbSTMI-GFP and a control with and without MG132 treatment in *N. benthamiana* leaves. Anti-GFP detected free GFP and $^{\Delta sp}$PbSTMI-GFP, while anti-myc detected AtSNC1-TAP. (B). Co-immunoprecipitation assay shows no interaction between $^{\Delta sp}$PbSTMI-GFP and AtSNC1-TAP using GFP-trap agarose beads. Protein loading was confirmed by Ponceau S staining of the Rubisco large subunit, and molecular weights (kDa) for GFP, AtSNC1-TAP, $^{\Delta sp}$PbSTMI-GFP, and RuBisCo LSU are indicated.

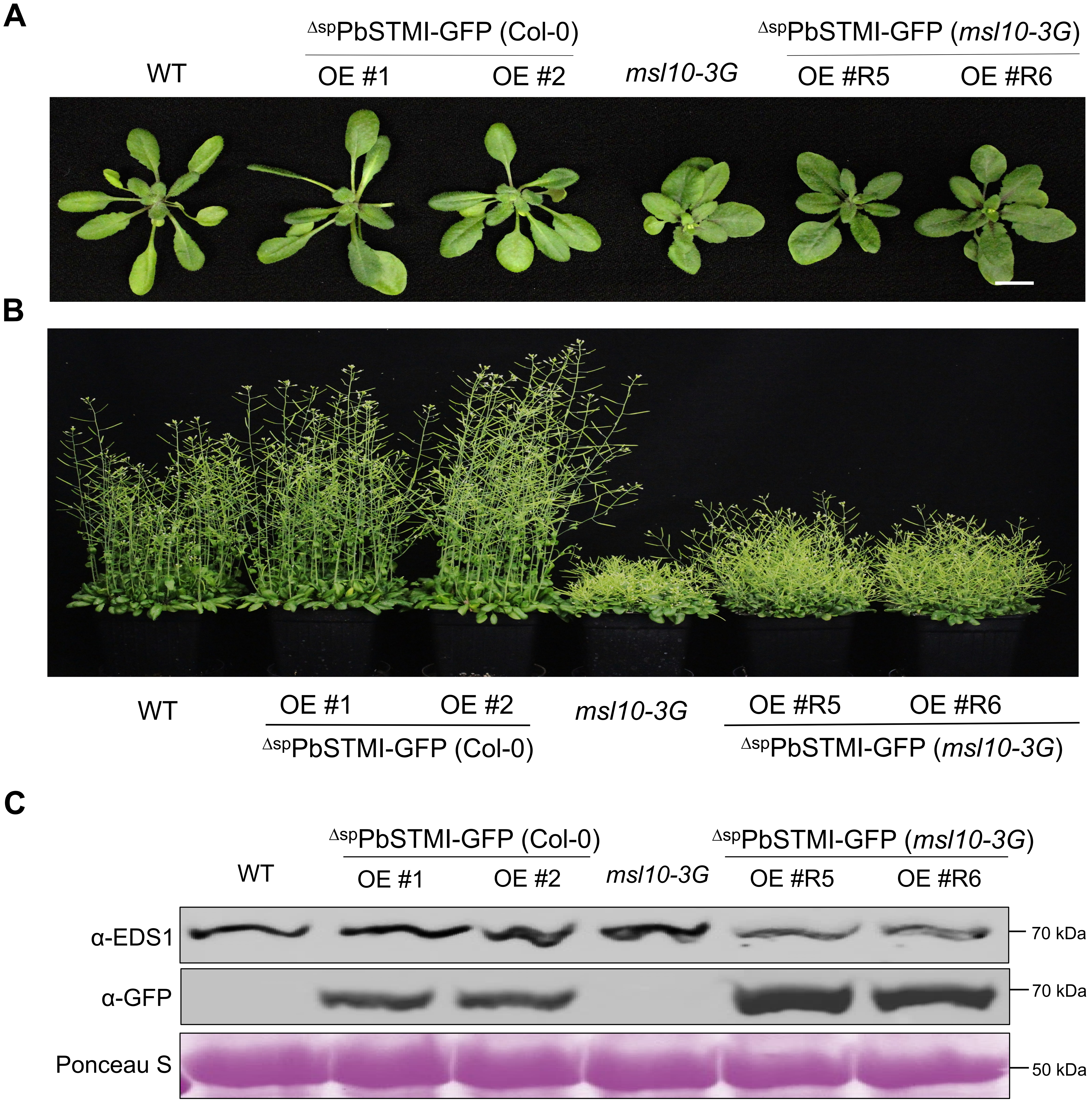


**Figure S9: $^{\Delta sp}$PbSTMI does not attenuate *msl10-3G*-mediated autoimmunity.** (A). Vegetative above-ground phenotypes of four week-old Arabidopsis WT, *msl10-3G,* $^{\Delta sp}$PbSTMI-GFP (Col-0) and $^{\Delta sp}$PbSTMI-GFP (*msl10-3G*) lines grown in soil at 22ºC. Scale bar = 1 cm. (B). Reproductive phenotypes of seven week-old plants of the indicated genotypes. (C). Immunoblot of $^{\Delta sp}$PbSTMI-GFP and *At*EDS1 in WT, *msl10-3G,* $^{\Delta sp}$PbSTMI-GFP (Col-0) and $^{\Delta sp}$PbSTMI-GFP (*msl10-3G*) lines. Equal loading is shown by Ponceau S staining of RuBisCo LSU bands and the molecular weights (kDa) of GFP, AtEDS1 and RuBisCo LSU are provided. This experiment was repeated three times with consistent results.

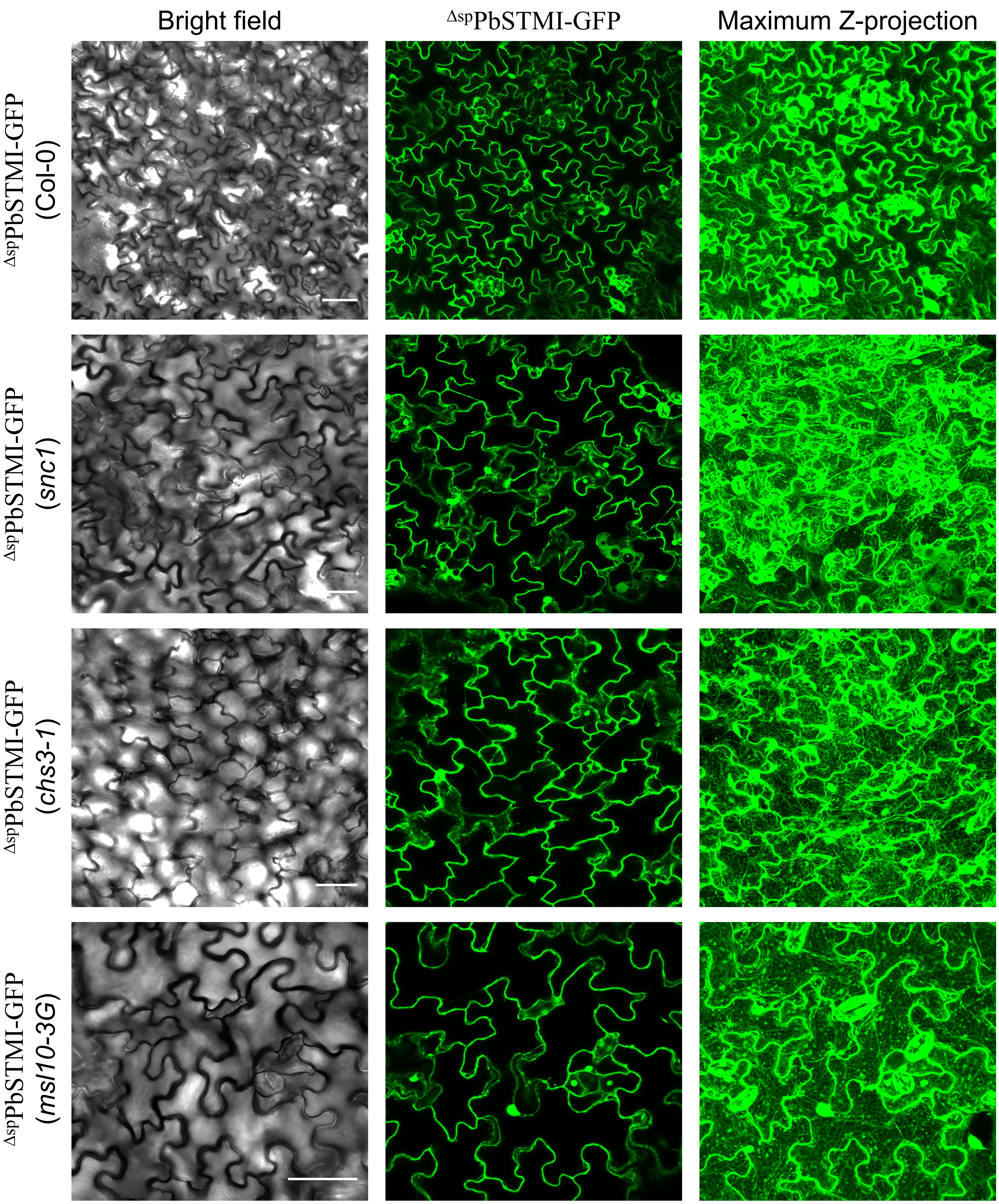


**Figure S10: Subcellular localization of $^{\Delta sp}$PbSTMI in Arabidopsis mutant backgrounds.** Subcellular localization of $^{\Delta sp}$PbSTMI-GFP in Arabidopsis OE lines in the autoimmune backgrounds, *snc1*, *chs3-1* and *msl10-3G*. The green channel shows nucleo-cytoplasmic localization of $^{\Delta sp}$PbSTMI-GFP in leaf epidermal cells. The right panel shows the maximum Z projection of the localization. Scale bars = 20 μm.

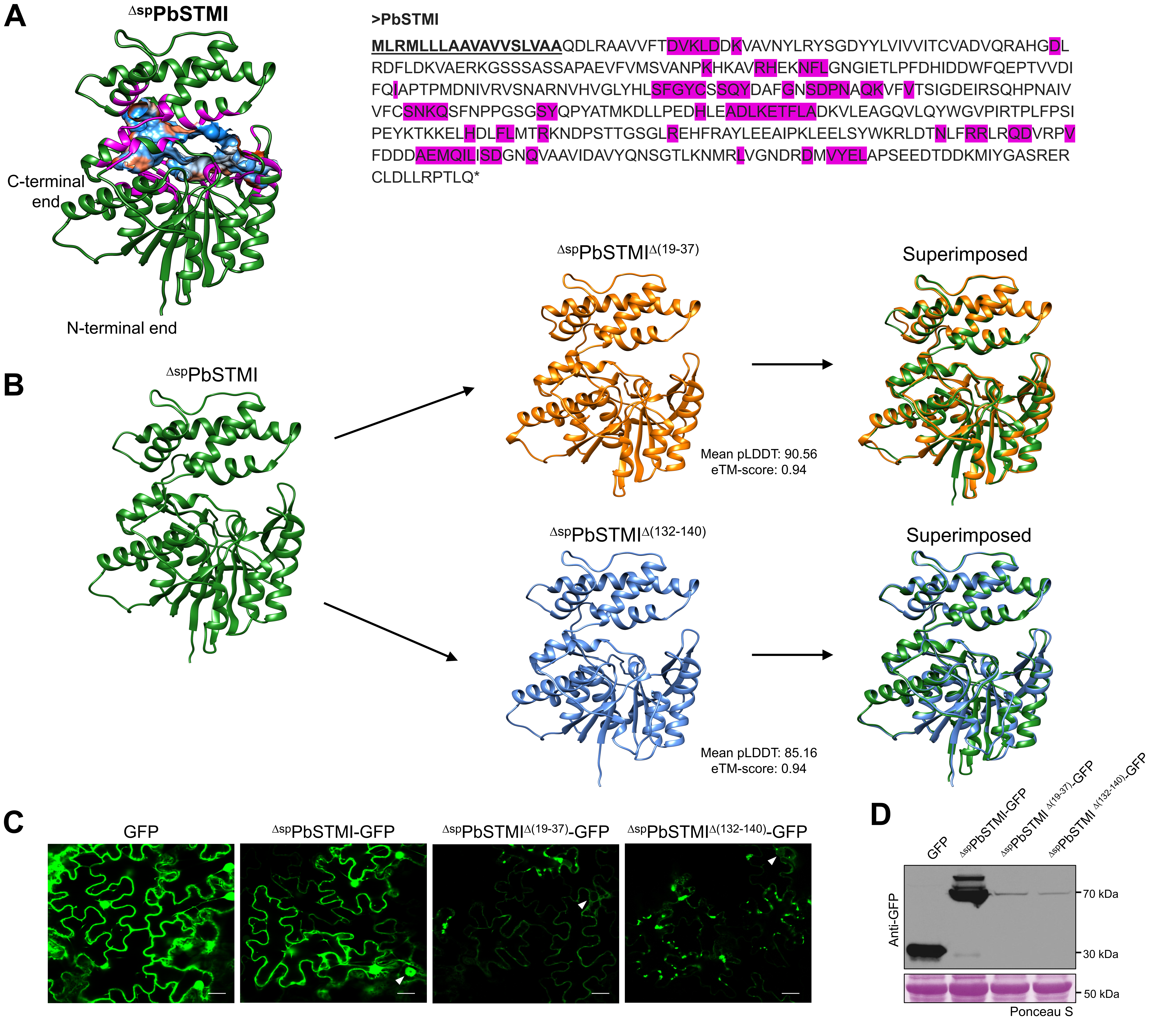


**Figure S11: Structural analysis of $^{\Delta sp}$PbSTMI and its N-terminal truncated mutants.** (A). 3D model of the $^{\Delta sp}$PbSTMI protein monomer, highlighting the binding pocket and the lining amino acid residues in pink. The pocket volume is indicated with colors (red—hydrophobic region, blue—hydrophilic) based on the hydrophobicity profile of the lining amino acids. (B). $^{\Delta sp}$PbSTMI N-terminal truncated mutants with 3D protein models, assessing structural deviation from their native conformations. eTM-score = estimated template modelling score. (C). Subcellular localization of $^{\Delta sp}$PbSTMI-GFP and GFP-tagged N-terminal truncated mutants in leaf epidermal cells of *N. benthamiana*. White arrowheads indicate nuclear and extra-nuclear localizations of the proteins. Scale bar = 10 µm. (D). Western blot of the expressed, GFP-tagged $^{\Delta sp}$PbSTMI N-terminal truncated mutants using anti-GFP antibody. Protein loading is shown by Ponceau S staining of Rubisco large subunit and the molecular weights (kDa) of GFP, $^{\Delta sp}$PbSTMI N-terminal truncated mutants, and RuBisCo LSU are provided.

**Data S1:** Multiple sequence alignment of $^{\Delta sp}$PbSTMI orthologs among plant pathogens.

Signal peptide

```
                                        1       10
PbSTMI              ...................MLRMLLLAAVAV.............VSLVAA
PbSTMI-L1           ...................MLRMSLLAAVAV.............VSLVAA
PbSTMI-L2           ...................MLRMLLLAAVAV.............VSLVAA
PbSTMI-L3           ...................MLRTLLLAAAAV.............VAVA..
Uma_XP_011389271.1  ..MWLRP....ISTILIGF....AILVTL.................VSA.
Uho_XP_041410349.1  ..MWLRP....LSTILLSS....LFLFAS.................VCA.
Uho_UTT92214.1      ..MWLRP....LSTILLSS....LFLFAS.................VCA.
Usp_SPC66142.1      ..MWLRP....LSTILLSS....LFLFAS.................VCA.
Utr_SPO24609.1      ..MGFQP....FPSLLVGL....VLLIAS.................VCA.
Ubr_SAM82506.1      ..MWLRP....LSTILLSS....LFLFAS.................VCA.
Usp_SOV03971.1      ..MWLRP....LSTILLSS....LFLFAS.................VCA.
Sre_SJX63044.1      ..MWFHP....IHTVLYGL....LVLAAV.................VRA.
Sre_CBQ73209.1      ..MRFHP....IHTVLYGL....LVLAAV.................VRA.
Ssc_CDU26470.1      ..MWFHP....IHTILYGF....LILAAL.................VRA.
Phu_XP_012187602.1  ..MWFQR....VSLLALSV....LLLSVA.................VRA.
Map_ETS62003.1      ..MHAGRFFGIVPLALIGALATSLLLSVSL..VSAAGPE....RFATAPN
Man_SPO45247.1      ..MHAGRFFGIVPLALIAALATTLLFSVSL..VSAAGPD....RFASAPY
Mpe_CDI55289.1      ..MWFRP....LLAVSLGL....LIAAIS.................VQAQ
Mpe_CDI52561.1      ..MWFRP....LLAVSLGL....LIAAIS.................VQAQ
Kbr_XP_016290109.1  ..MWFRS....LPLFLLAI....LGLALS.................AAA.
Tcy_PWY99562.1      ..MHFAA..GV...RACGLMLAAVVLATCAVAAPAFGPSSASSSSSNGAG
Cab_KAI3559566.1    ..MTRHF......LFFAV....LCWIA..................THSA
Csp_TDZ27191.1      ..MQFPR......FVLAL....VCWAA..................AQTA
Csi_TEA13839.1      ..MQFPR......FVLAL....VCWAA..................AQTA
Cob_TDZ24401.1      ..MQFPR......FVLAL....VCWAA..................AQTA
Csc_KAH8421623.1    ..MSRHF......LFFAV....LCWIA..................AHSA
Cfi_EXF77787.1      ..MSRNY......LFFVV....LCWIA..................THAA
Cny_KXH41789.1      ..MSRHF......LFFAV....LCWIA..................AHSA
Csi_CKXH31893.1     ..MSRHF......LFFAV....LCWIA..................AHSA
Cgr_XP_008100761.1  MAIASHL......SILGL....VCWLF..................AQTV
Psp_TLD34168.1      ..MTFHL......LIVTL....LFWGF..................TQVA
Pgr_KAI6345083.1    .MISFRL......LTLTL....LCWGF..................TQIA
Pgr_XP_030977794.1  .MISFRL......LTLTL....LCWGF..................TQIA
Por_KAI6598559.1    ..MSSRF......LIFSL....LFWGL..................AQVA
Por_XP_003716019.1  ..MSSRF......LIFSL....LFWGL..................AQVA
Por_KAH9427683.1    ..MSSRF......LIFSL....LFWGL..................AQVA
Por_KAI6324114.1    ..MSSRF......LIFSL....LFWGL..................AQVA
Por_KAI6271750.1    ..MSSRF......LIFSL....LFWGL..................AQVA
Por_KAH8844570.1    ..MSSRF......LIFSL....LFWGL..................AQVA
Por_KAI6541088.1    ..MSSRF......LIFSL....LFWGL..................AQVA
Por_KAI6445464.1    ..MSSRF......LIFSL....LFWGL..................AQVA
```

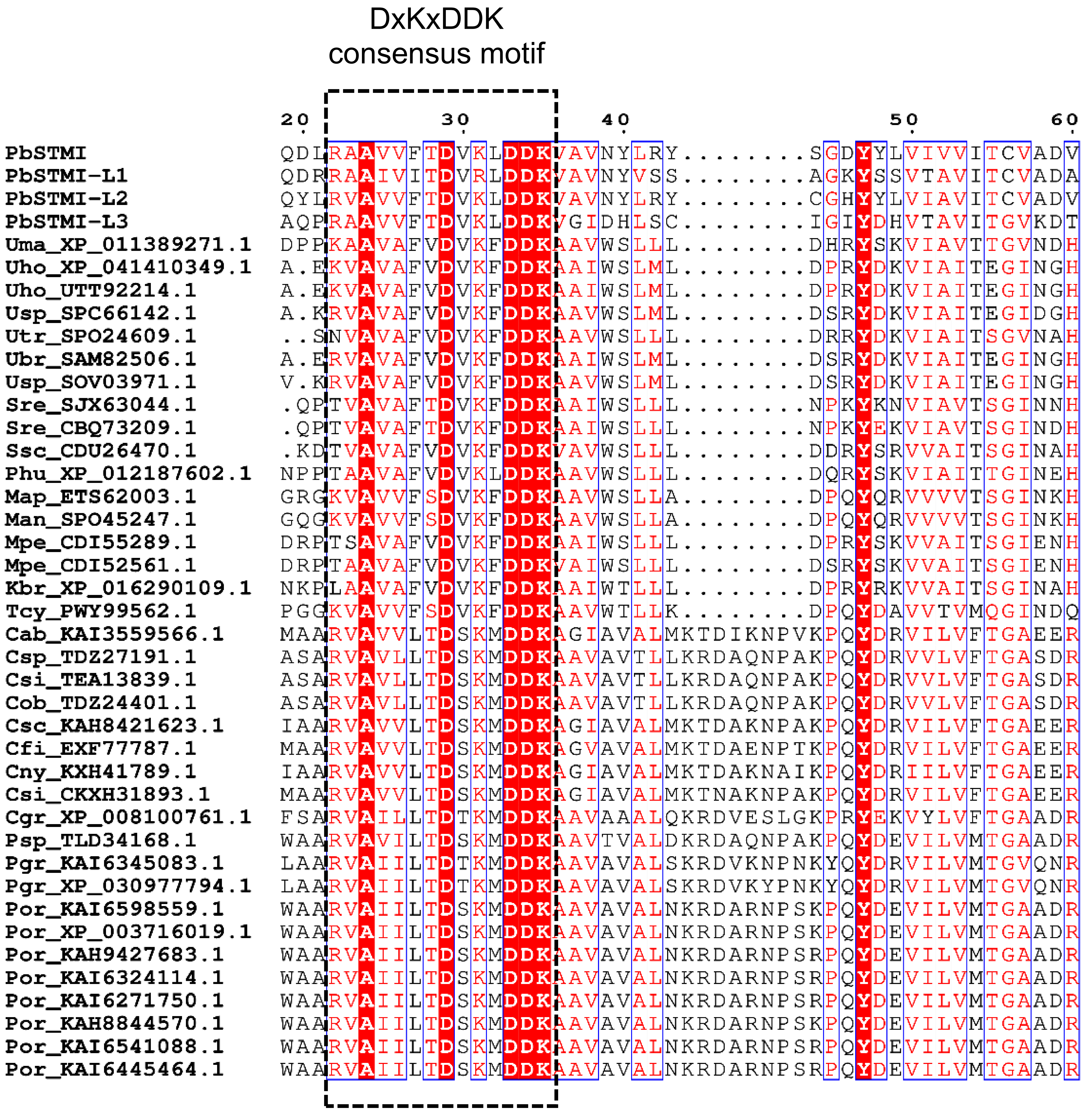
DxKxDDK
consensus motif
20 30 40 50 60
PbSTMI QDLRAAVVFTDVKLDDKVAVNYLRY........SGDYYLVIVVITCVADV
PbSTMI-L1 QDRRAAIVITDVRLDDKVAVNYVSS........AGKYSSVTAVITCVADA
PbSTMI-L2 QYLRVAVVFTDVKLDDKVAVNYLRY........CGHYYLVIAVITCVADV
PbSTMI-L3 AQPRAAVVFTDVKLDDKVGIDHLSC........IGIYDHVTAVITGVKDT
Uma_XP_011389271.1 DPPKAAVAFVDVKFDDKAAVWSLLL........DHRYSKVIAVTTGVNDH
Uho_XP_041410349.1 A.EKVAVAFVDVKFDDKAAIWSLML........DPRYDKVIAITEGINGH
Uho_UTT92214.1 A.EKVAVAFVDVKFDDKAAIWSLML........DPRYDKVIAITEGINGH
Usp_SPC66142.1 A.KRVAVAFVDVKFDDKAAVWSLML........DSRYDKVIAITEGIDGH
Utr_SPO24609.1 ..SNVAVAFVDVKFDDKAAVWSLLL........DRRYDKVIAITSGVNAH
Ubr_SAM82506.1 A.ERVAVAFVDVKFDDKAAIWSLML........DSRYDKVIAITEGINGH
Usp_SOV03971.1 V.KRVAVAFVDVKFDDKAAVWSLML........DSRYDKVIAITEGINGH
Sre_SJX63044.1 .QPTVAVAFTDVKFDDKAAIWSLLL........NPKYKNVIAVTSGINNH
Sre_CBQ73209.1 .QPTVAVAFTDVKFDDKAAIWSLLL........NPKYEKVIAVTSGINDH
Ssc_CDU26470.1 .KDTVAVAFVDVKFDDKVAVWSLLL........DDRYSRVVAITSGINAH
Phu_XP_012187602.1 NPPTAAVAFVDVKLDDKAAVWSLLL........DQRYSKVIAITTGINEH
Map_ETS62003.1 GRGKVAVVFSDVKFDDKAAVWSLLA........DPQYQRVVVVTSGINKH
Man_SPO45247.1 GQGKVAVVFSDVKFDDKAAVWSLLA........DPQYQRVVVVTSGINKH
Mpe_CDI55289.1 DRPTSAVAFVDVKFDDKAAIWSLLL........DPRYSKVVAITSGIENH
Mpe_CDI52561.1 DRPTAAVAFVDVKFDDKVAIWSLLL........DSRYSKVVAITSGIENH
Kbr_XP_016290109.1 NKPLAAVAFVDVKFDDKAAIWTLLL........DPRYRKVVAITSGINAH
Tcy_PWY99562.1 PGGKVAVVFSDVKFDDKAAVWTLLK........DPQYDAVVTVMQGINDQ
Cab_KAI3559566.1 MAARVAVVLTDSKMDDKAGIAVALMKTDIKNPVKPQYDRVILVFTGAEER
Csp_TDZ27191.1 ASARVAVLLTDSKMDDKAAVAVTLLKRDAQNPAKPQYDRVVLVFTGASDR
Csi_TEA13839.1 ASARVAVLLTDSKMDDKAAVAVTLLKRDAQNPAKPQYDRVVLVFTGASDR
Cob_TDZ24401.1 ASARVAVLLTDSKMDDKAAVAVTLLKRDAQNPAKPQYDRVVLVFTGASDR
Csc_KAH8421623.1 IAARVAVVLTDSKMDDKAGIAVALMKTDAKNPAKPQYDRVILVFTGAEER
Cfi_EXF77787.1 MAARVAVVLTDSKMDDKAGVAVALMKTDAENPTKPQYDRVILVFTGAEER
Cny_KXH41789.1 IAARVAVVLTDSKMDDKAGIAVALMKTDAKNAIKPQYDRIILVFTGAEER
Csi_CKXH31893.1 MAARVAVVLTDSKMDDKAGIAVALMKTNAKNPAKPQYDRVILVFTGAEER
Cgr_XP_008100761.1 FSARVAILLTDTKMDDKAAVAAALQKRDVESLGKPRYEKVYLVFTGAADR
Psp_TLD34168.1 WAARVAVILTDSKMDDKAAVTVALDKRDAQNPAKPQYDEVILVMTGAADR
Pgr_KAI6345083.1 LAARVAIILTDTKMDDKAAVAVALSKRDVKNPNKYQYDRVILVMTGVQNR
Pgr_XP_030977794.1 LAARVAIILTDTKMDDKAAVAVALSKRDVKYPNKYQYDRVILVMTGVQNR
Por_KAI6598559.1 WAARVAIILTDSKMDDKAAVAVALNKRDARNPSKPQYDEVILVMTGAADR
Por_XP_003716019.1 WAARVAIILTDSKMDDKAAVAVALNKRDARNPSKPQYDEVILVMTGAADR
Por_KAH9427683.1 WAARVAIILTDSKMDDKAAVAVALNKRDARNPSRPQYDEVILVMTGAADR
Por_KAI6324114.1 WAARVAIILTDSKMDDKAAVAVALNKRDARNPSRPQYDEVILVMTGAADR
Por_KAI6271750.1 WAARVAIILTDSKMDDKAAVAVALNKRDARNPSRPQYDEVILVMTGAADR
Por_KAH8844570.1 WAARVAIILTDSKMDDKAAVAVALNKRDARNPSKPQYDEVILVMTGAADR
Por_KAI6541088.1 WAARVAIILTDSKMDDKAAVAVALNKRDARNPSRPQYDEVILVMTGAADR
Por_KAI6445464.1 WAARVAIILTDSKMDDKAAVAVALNKRDARNPSRPQYDEVILVMTGAADR

```
                              70        80        90       100       110
PbSTMI                    QRAHGDLRDFLDKVAERKGSSSASSAPAEVFVMSVANPKHKAVRHEKNFL
PbSTMI-L1                 DRAHGDLRDFLVKTYRNG.....NS...PVRIMRVPNPRSEAVRYEKFFI
PbSTMI-L2                 QRAHGDLRDFLSETYRGD.....HSTACDVLIMSVPNAKQKPAPHEKFFI
PbSTMI-L3                 QRAHSDLRDFLNSMELLRAA...SVKCSPVFVLGVESMNEKPVPHEVNFI
Uma_XP_011389271.1        GRAAFELDGYLNRQNAM.ANI..QFDKNKLQILQGSNPLGKAAPHEAWWD
Uho_XP_041410349.1        EKAAQELQDYLGRQNNI.ANV..RVDLGKLRMFSGSNVLGESPGHESWWN
Uho_UTT92214.1            EKAAQELQDYLGRQNNI.ANV..RVDLGKLRMFSGSNVLGESPGHESWWN
Usp_SPC66142.1            GKAAQELQDYLDRQNDI.ANV..RVNLRKLKMFSGSNVLGKPPKHERWWK
Utr_SPO24609.1            GQAAHELQSYLDRQNDM.ANV..KFDSGKVQILEGSNVLRANAPHEAWWQ
Ubr_SAM82506.1            EKAAQELQDYLGRQNDI.AKV..RVNLRKLRMFSGSNVLGEPPKHESWWK
Usp_SOV03971.1            GKAAQELQDYLYRQNHI.INV..RVNLEKLRMFSGSNVLGKPPKHESWWK
Sre_SJX63044.1            WRAAQELSEYLDRQNSL.PVG.KHFDTRKLVILKGSNHQDQPAPHEVWWK
Sre_CBQ73209.1            WRAAQELSEYLDRQNSL.PVG.KHFDTRKLVILKGSNHQDQPAPHEVWWK
Ssc_CDU26470.1            GQAAEELNEYLDRQNGF.QGS.KRFNKRKLTILQGTNGLGKKAPHEEWWN
Phu_XP_012187602.1        RRAAYELDQYVDRQNAM.ANV..KFNKDKLEILQGSNPLPSSAPHEAWWG
Map_ETS62003.1            RNAAQELVNFVDRQSE...HLGSRNLKDKLKLIEGGNPLGMEAPHEVWYN
Man_SPO45247.1            RNAAQELVNFVDRQND...HLGSRDLKDKLKIMQGGNPLGVDAPHEVWYD
Mpe_CDI55289.1            GKAASELEQYLERQNAM.PNV.KKVDLDKIQIFRGTNPFGEEAPHEHWWN
Mpe_CDI52561.1            GKAASELEQYLERQNAM.SNV.RKVNLGKIQIFRGTNPFGAEVPHEYWWN
Kbr_XP_016290109.1        DQAAYELYRYVHDQNNT.PYR..KVDMSKLEILQGTNALGRAAPHEEWWA
Tcy_PWY99562.1            ERAAKALLDFVGRMNSI.PVAGARVDPSKLRLLAGSNVLGIDAPHEVWYR
Cab_KAI3559566.1          SKMALAMSDYLNRAEKHQPALPAGWETRTAWFQTKTNVLGQAAPHESWYK
Csp_TDZ27191.1            NKIASGMVDYLARAERRTPPVPKDWRDRTGWLQARTNVMVEEVPHERWYK
Csi_TEA13839.1            NKIASGMVDYLARAERRTPPVPKDWRDRTGWLQARTNVMVEEVPHERWYK
Cob_TDZ24401.1            NKIASGMVDYLARAERRTPPVPKDWRDRTGWLQARTNVMVEEVPHERWYK
Csc_KAH8421623.1          SRMGLAMSDYLTRAEKHQPALPAGWESRTAWFQTKTNVLGQAAPHESWYK
Cfi_EXF77787.1            SKMGLAMSEYLTRAENHQPALPAGWEKRTAWLQTRTNVLGQAAPHESWYR
Cny_KXH41789.1            SKLGLAMSDYLTRAEKHQPALPAGWESRTAWFQTKTNVLGQAAPHESWYK
Csi_CKXH31893.1           SKMGLAMSDYLARAEKHQPSLPAGWESRTAWFQTKTNVLGQAAPHESWYK
Cgr_XP_008100761.1        NKIAEGMSQYLQRAEKHNPALPADWRKRTEWFQTQTNVMGKAVAHERWYK
Psp_TLD34168.1            NKIAEGMRQYLIRAERHRPALSPDWRTRIRWFQTGGNVLGKVVAHERWYK
Pgr_KAI6345083.1          QIIAEGMKEYLYRAQSRRPGISHGWDTRVSWFQTRENIMDKDVSHELWYK
Pgr_XP_030977794.1        QIIAEGMKEFLYRAQSHRPGIPHGWDTRVSWFQTRGNIMDKDVSHELWYK
Por_KAI6598559.1          DNIAEGMRQYLQRAERHQPALSSDWKNRVSWFQTRGNVMGKVVAHERWYK
Por_XP_003716019.1        DNIAEGMRQYLQRAERHQPALSSDWKNRVSWFQTRGNVMGKVVAHERWYK
Por_KAH9427683.1          DNIAEGMRQYLQRAERHQPALSSDWKNRVSWFQTRGNVMGKVVAHERWYK
Por_KAI6324114.1          DNIAEGMRQYLQRAERHQPALSSDWKNRVSWFQTRGNVMGKVVAHERWYK
Por_KAI6271750.1          DNIAEGMRQYLQRAERHQPALSSDWKNRVSWFQTRGNVMGKVVAHERWYK
Por_KAH8844570.1          DNIAEGMRQYLQRAERHQPALSSDWKNRVSWFQTRGNVMGKVVAHERWYK
Por_KAI6541088.1          DNIAEGMRQYLQRAERHQPALSSDWKNRVSWFQTRGNVMGKVVAHERWYK
Por_KAI6445464.1          DNIAEGMRQYLQRAERHQPALSSDWKNRVSWFQTRGNVMGKVVAHERWYK
```

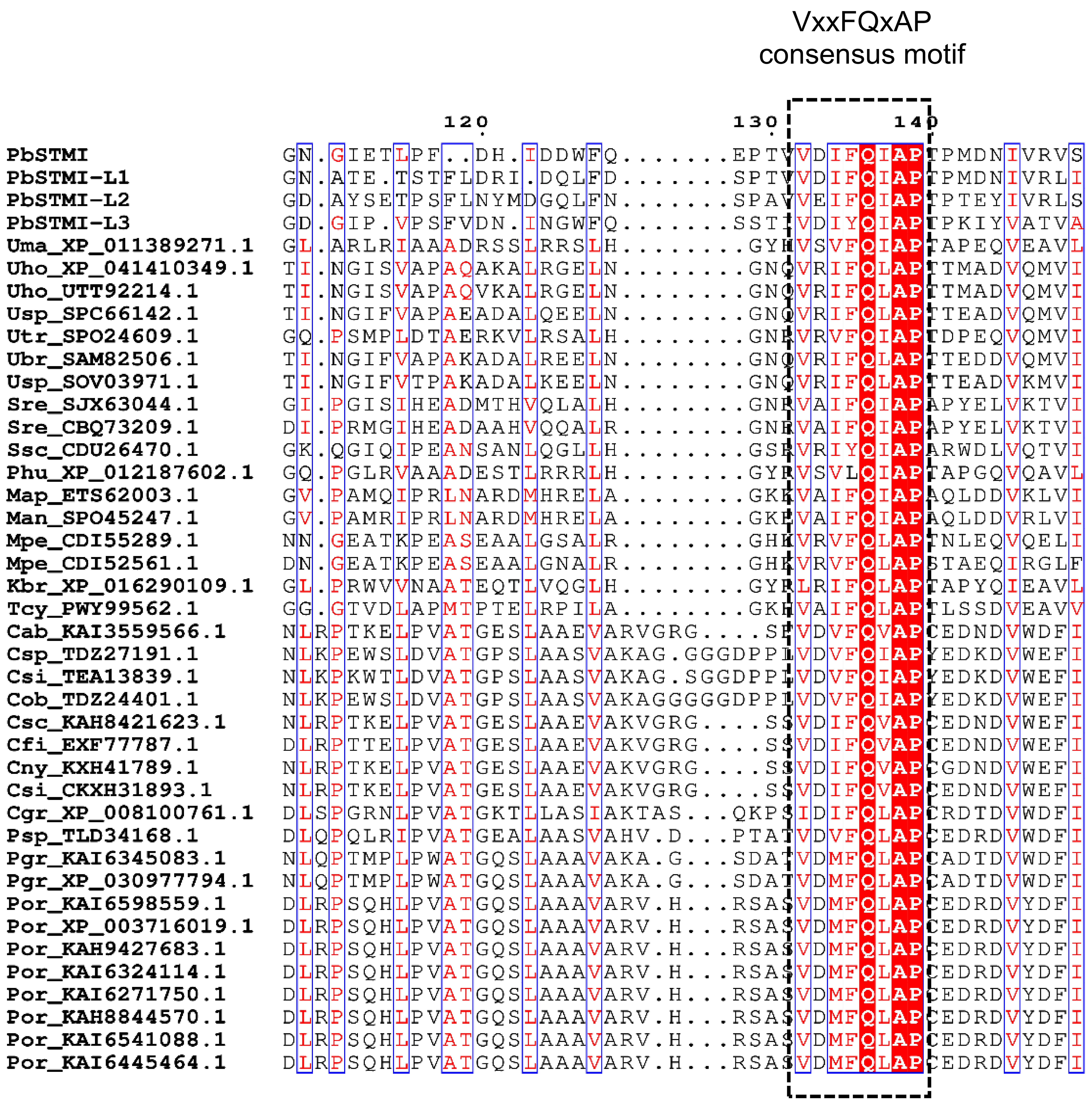
VxxFQxAP
consensus motif
120
130
140
PbSTMI
PbSTMI-L1
PbSTMI-L2
PbSTMI-L3
Uma_XP_011389271.1
Uho_XP_041410349.1
Uho_UTT92214.1
Usp_SPC66142.1
Utr_SPO24609.1
Ubr_SAM82506.1
Usp_SOV03971.1
Sre_SJX63044.1
Sre_CBQ73209.1
Ssc_CDU26470.1
Phu_XP_012187602.1
Map_ETS62003.1
Man_SPO45247.1
Mpe_CDI55289.1
Mpe_CDI52561.1
Kbr_XP_016290109.1
Tcy_PWY99562.1
Cab_KAI3559566.1
Csp_TDZ27191.1
Csi_TEA13839.1
Cob_TDZ24401.1
Csc_KAH8421623.1
Cfi_EXF77787.1
Cny_KXH41789.1
Csi_CKXH31893.1
Cgr_XP_008100761.1
Psp_TLD34168.1
Pgr_KAI6345083.1
Pgr_XP_030977794.1
Por_KAI6598559.1
Por_XP_003716019.1
Por_KAH9427683.1
Por_KAI6324114.1
Por_KAI6271750.1
Por_KAH8844570.1
Por_KAI6541088.1
Por_KAI6445464.1

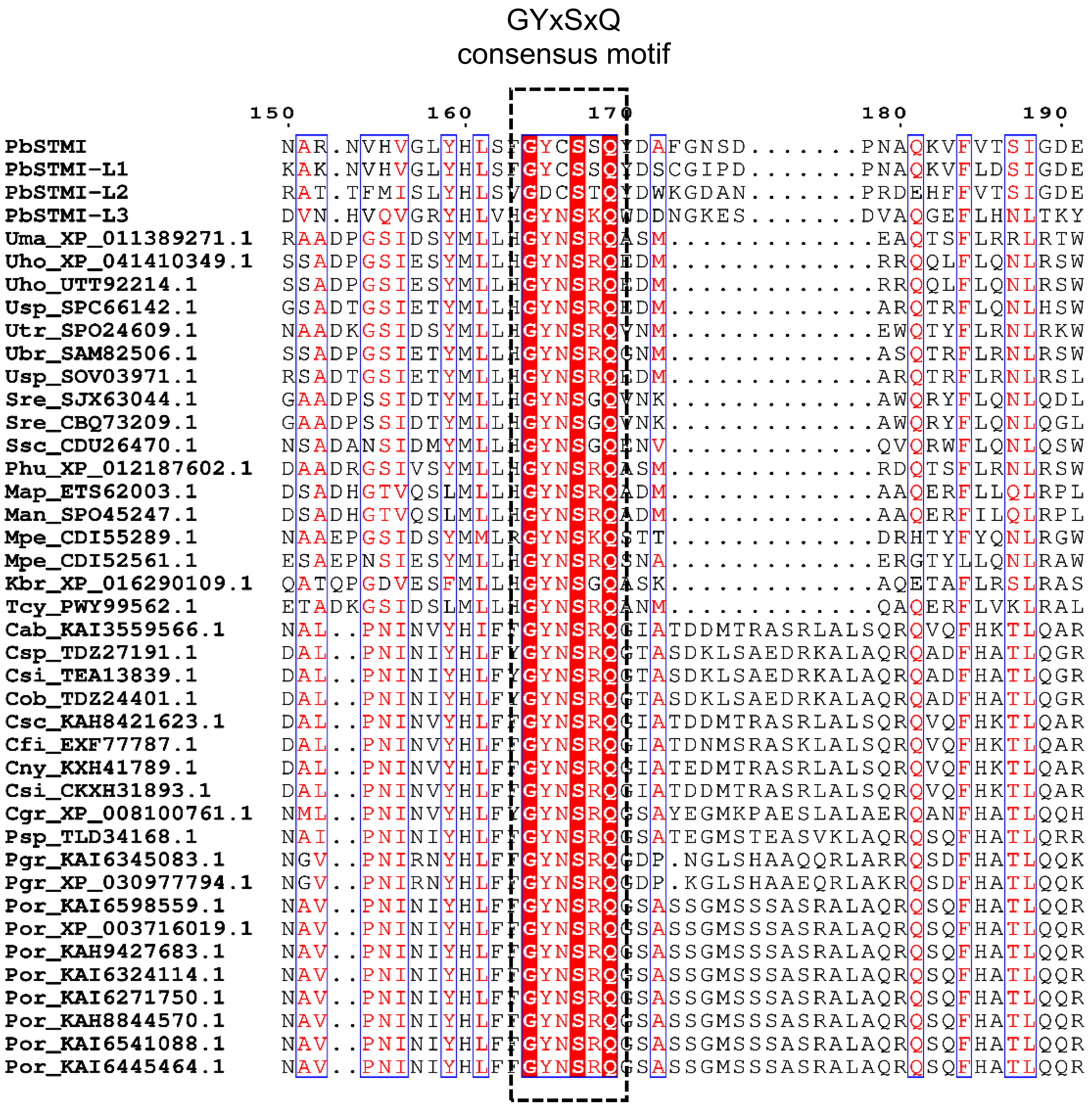
GYxSxQ
consensus motif
150 160 170 180 190
PbSTMI NAR.NVHVGLYHLSFGYCSSQYDAFGNSD.......PNAQKVFVTSIGDE
PbSTMI-L1 KAK.NVHVGLYHLSFGYCSSQYDSCGIPD.......PNAQKVFLDSIGDE
PbSTMI-L2 RAT.TFMISLYHLSVGDCSTQYDWKGDAN.......PRDEHFFVTSIGDE
PbSTMI-L3 DVN.HVQVGRYHLVHGYNSKQWDDNGKES.......DVAQGEFLHNLTKY
Uma_XP_011389271.1 RAADPGSIDSYMLLHGYNSRQASM.............EAQTSFLRRLRTW
Uho_XP_041410349.1 SSADPGSIESYMLLHGYNSRQEDM.............RRQQLFLQNLRSW
Uho_UTT92214.1 SSADPGSIESYMLLHGYNSRQEDM.............RRQQLFLQNLRSW
Usp_SPC66142.1 GSADTGSIETYMLLHGYNSRQEDM.............ARQTRFLQNLHSW
Utr_SPO24609.1 NAADKGSIDSYMLLHGYNSRQYNM.............EWQTYFLRNLRKW
Ubr_SAM82506.1 SSADPGSIETYMLLHGYNSRQGNM.............ASQTRFLRNLRSW
Usp_SOV03971.1 RSADTGSIETYMLLHGYNSRQEDM.............ARQTRFLRNLRSL
Sre_SJX63044.1 GAADPSSIDTYMLLHGYNSGQYNK.............AWQRYFLQNLQDL
Sre_CBQ73209.1 GAADPSSIDTYMLLHGYNSGQYNK.............AWQRYFLQNLQGL
Ssc_CDU26470.1 NSADANSIDMYMLLHGYNSGQENV.............QVQRWFLQNLQSW
Phu_XP_012187602.1 DAADRGSIVSYMLLHGYNSRQASM.............RDQTSFLRNLRSW
Map_ETS62003.1 DSADHGTVQSLMLLHGYNSRQADM.............AAQERFLLQLRPL
Man_SPO45247.1 DSADHGTVQSLMLLHGYNSRQADM.............AAQERFILQLRPL
Mpe_CDI55289.1 NAAEPGSIDSYMMLRGYNSKQSTT.............DRHTYFYQNLRGW
Mpe_CDI52561.1 ESAEPNSIESYMLLHGYNSRQSNA.............ERGTYLLQNLRAW
Kbr_XP_016290109.1 QATQPGDVESFMLLHGYNSGQASK.............AQETAFLRSLRAS
Tcy_PWY99562.1 ETADKGSIDSLMLLHGYNSRQANM.............QAQERFLVKLRAL
Cab_KAI3559566.1 NAL..PNINVYHIFFGYNSRQGIATDDMTRASRLALSQRQVQFHKTLQAR
Csp_TDZ27191.1 DAL..PNINIYHLFYGYNSRQGTASDKLSAEDRKALAQRQADFHATLQGR
Csi_TEA13839.1 DAL..PNINIYHLFYGYNSRQGTASDKLSAEDRKALAQRQADFHATLQGR
Cob_TDZ24401.1 DAL..PNINIYHLFYGYNSRQGTASDKLSAEDRKALAQRQADFHATLQGR
Csc_KAH8421623.1 DAL..PNINVYHLFFGYNSRQGIATDDMTRASRLALSQRQVQFHKTLQAR
Cfi_EXF77787.1 DAL..PNINVYHLFFGYNSRQGIATDNMSRASKLALSQRQVQFHKTLQAR
Cny_KXH41789.1 DAL..PNINVYHLFFGYNSRQGIATEDMTRASRLALSQRQVQFHKTLQAR
Csi_CKXH31893.1 DAL..PNINVYHLFFGYNSRQGIATDDMTRASRLALSQRQVQFHKTLQAR
Cgr_XP_008100761.1 NML..PNINVYHLFYGYNSRQGSAYEGMKPAESLALAERQANFHATLQQH
Psp_TLD34168.1 NAI..PNINIYHLFFGYNSRQGSATEGMSTEASVKLAQRQSQFHATLQRR
Pgr_KAI6345083.1 NGV..PNIRNYHLFFGYNSRQGDP.NGLSHAAQQRLARRQSDFHATLQQK
Pgr_XP_030977794.1 NGV..PNIRNYHLFFGYNSRQGDP.KGLSHAAEQRLAKRQSDFHATLQQK
Por_KAI6598559.1 NAV..PNINIYHLFFGYNSRQGSASSGMSSSASRALAQRQSQFHATLQQR
Por_XP_003716019.1 NAV..PNINIYHLFFGYNSRQGSASSGMSSSASRALAQRQSQFHATLQQR
Por_KAH9427683.1 NAV..PNINIYHLFFGYNSRQGSASSGMSSSASRALAQRQSQFHATLQQR
Por_KAI6324114.1 NAV..PNINIYHLFFGYNSRQGSASSGMSSSASRALAQRQSQFHATLQQR
Por_KAI6271750.1 NAV..PNINIYHLFFGYNSRQGSASSGMSSSASRALAQRQSQFHATLQQR
Por_KAH8844570.1 NAV..PNINIYHLFFGYNSRQGSASSGMSSSASRALAQRQSQFHATLQQR
Por_KAI6541088.1 NAV..PNINIYHLFFGYNSRQGSASSGMSSSASRALAQRQSQFHATLQQR
Por_KAI6445464.1 NAV..PNINIYHLFFGYNSRQGSASSGMSSSASRALAQRQSQFHATLQQR

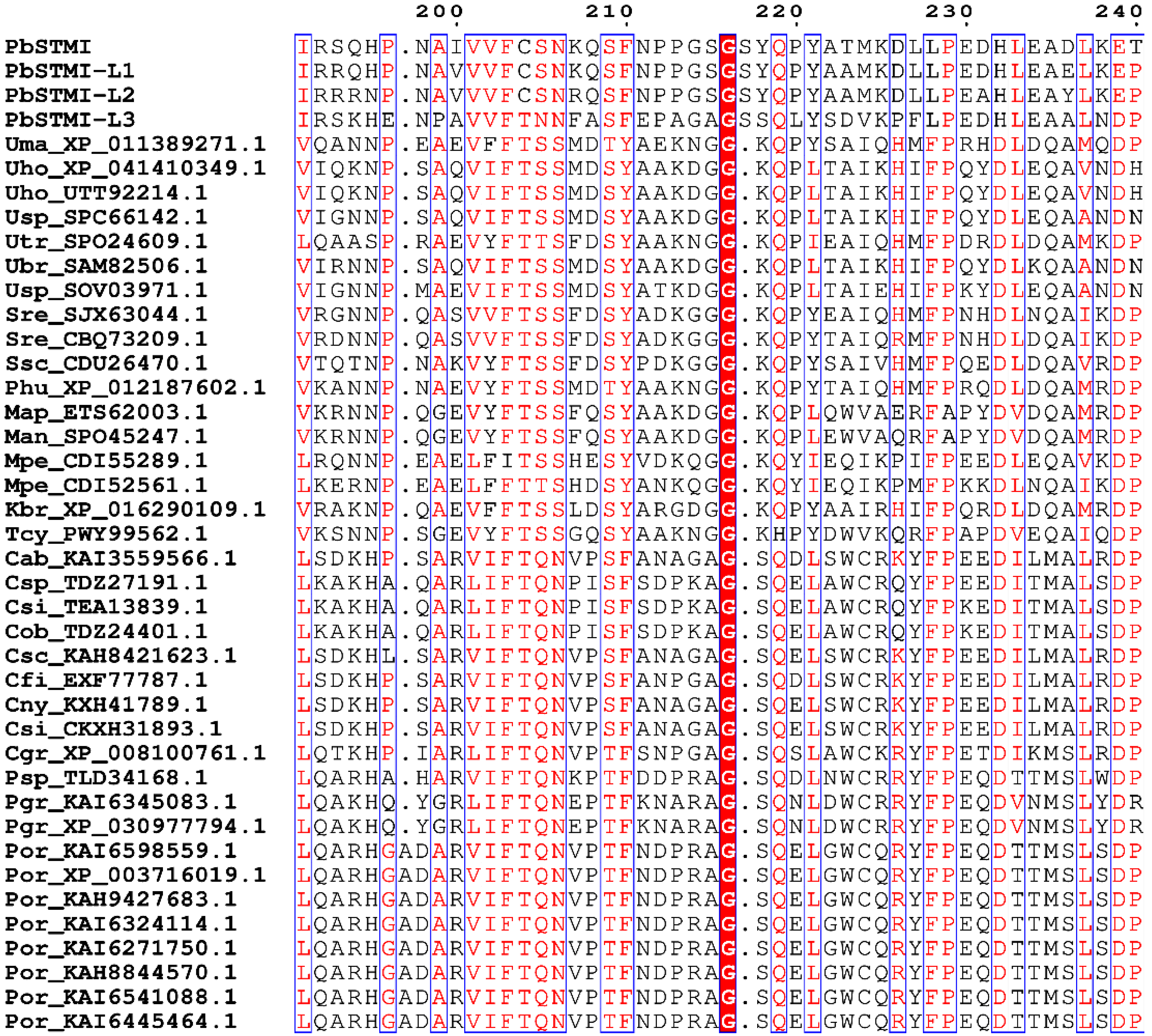
200 210 220 230 240
PbSTMI IRSQHP.NAIVVFCSNKQSFNPPGSGSYQPYATMKDLLPEDHLEADLKET
PbSTMI-L1 IRRQHP.NAVVVFCSNKQSFNPPGSGSYQPYAAMKDLLPEDHLEAELKEP
PbSTMI-L2 IRRRNP.NAVVVFCSNRQSFNPPGSGSYQPYAAMKDLLPEAHLEAYLKEP
PbSTMI-L3 IRSKHE.NPAVVFTNNFASFEPAGAGSSQLYSDVKPFLPEDHLEAALNDP
Uma_XP_011389271.1 VQANNP.EAEVFFTSSMDTYAEKNGG.KQPYSAIQHMFPRHDLDQAMQDP
Uho_XP_041410349.1 VIQKNP.SAQVIFTSSMDSYAAKDGG.KQPLTAIKHIFPQYDLEQAVNDH
Uho_UTT92214.1 VIQKNP.SAQVIFTSSMDSYAAKDGG.KQPLTAIKHIFPQYDLEQAVNDH
Usp_SPC66142.1 VIGNNP.SAQVIFTSSMDSYAAKDGG.KQPLTAIKHIFPQYDLEQAANDN
Utr_SPO24609.1 LQAASP.RAEVYFTTSFDSYAAKNGG.KQPIEAIQHMFPDRDLDQAMKDP
Ubr_SAM82506.1 VIRNNP.SAQVIFTSSMDSYAAKDGG.KQPLTAIKHIFPQYDLKQAANDN
Usp_SOV03971.1 VIGNNP.MAEVIFTSSMDSYATKDGG.KQPLTAIEHIFPKYDLEQAANDN
Sre_SJX63044.1 VRGNNP.QASVVFTSSFDSYADKGGG.KQPYEAIQHMFPNHDLNQAIKDP
Sre_CBQ73209.1 VRDNNP.QASVVFTSSFDSYADKGGG.KQPYTAIQRMFPNHDLDQAIKDP
Ssc_CDU26470.1 VTQTNP.NAKVYFTSSFDSYPDKGGG.KQPYSAIVHMFPQEDLDQAVRDP
Phu_XP_012187602.1 VKANNP.NAEVYFTSSMDTYAAKNGG.KQPYTAIQHMFPRQDLDQAMRDP
Map_ETS62003.1 VKRNNP.QGEVYFTSSFQSYAAKDGG.KQPLQWVAERFAPYDVDQAMRDP
Man_SPO45247.1 VKRNNP.QGEVYFTSSFQSYAAKDGG.KQPLEWVAQRFAPYDVDQAMRDP
Mpe_CDI55289.1 LRQNNP.EAELFITSSHESYVDKQGG.KQYIEQIKPIFPEEDLEQAVKDP
Mpe_CDI52561.1 LKERNP.EAELFFTTSHDSYANKQGG.KQYIEQIKPMFPKKDLNQAIKDP
Kbr_XP_016290109.1 VRAKNP.QAEVFFTSSLDSYARGDGG.KQPYAAIRHIFPQRDLDQAMRDP
Tcy_PWY99562.1 VKSNNP.SGEVYFTSSGQSYAAKNGG.KHPYDWVKQRFPAPDVEQAIQDP
Cab_KAI3559566.1 LSDKHP.SARVIFTQNVPSFANAGAG.SQDLSWCRKYFPEEDILMALRDP
Csp_TDZ27191.1 LKAKHA.QARLIFTQNPISFSDPKAG.SQELAWCRQYFPEEDITMALSDP
Csi_TEA13839.1 LKAKHA.QARLIFTQNPISFSDPKAG.SQELAWCRQYFPKEDITMALSDP
Cob_TDZ24401.1 LKAKHA.QARLIFTQNPISFSDPKAG.SQELAWCRQYFPKEDITMALSDP
Csc_KAH8421623.1 LSDKHL.SARVIFTQNVPSFANAGAG.SQELSWCRKYFPEEDILMALRDP
Cfi_EXF77787.1 LSDKHP.SARVIFTQNVPSFANPGAG.SQDLSWCRKYFPEEDILMALRDP
Cny_KXH41789.1 LSDKHP.SARVIFTQNVPSFANAGAG.SQELSWCRKYFPEEDILMALRDP
Csi_CKXH31893.1 LSDKHP.SARVIFTQNVPSFANAGAG.SQELSWCRKYFPEEDILMALRDP
Cgr_XP_008100761.1 LQTKHP.IARLIFTQNVPTFSNPGAG.SQSLAWCKRYFPETDIKMSLRDP
Psp_TLD34168.1 LQARHA.HARVIFTQNKPTFDDPRAG.SQDLNWCRRYFPEQDTTMSLWDP
Pgr_KAI6345083.1 LQAKHQ.YGRLIFTQNEPTFKNARAG.SQNLDWCRRYFPEQDVNMSLYDR
Pgr_XP_030977794.1 LQAKHQ.YGRLIFTQNEPTFKNARAG.SQNLDWCRRYFPEQDVNMSLYDR
Por_KAI6598559.1 LQARHGADARVIFTQNVPTFNDPRAG.SQELGWCQRYFPEQDTTMSLSDP
Por_XP_003716019.1 LQARHGADARVIFTQNVPTFNDPRAG.SQELGWCQRYFPEQDTTMSLSDP
Por_KAH9427683.1 LQARHGADARVIFTQNVPTFNDPRAG.SQELGWCQRYFPEQDTTMSLSDP
Por_KAI6324114.1 LQARHGADARVIFTQNVPTFNDPRAG.SQELGWCQRYFPEQDTTMSLSDP
Por_KAI6271750.1 LQARHGADARVIFTQNVPTFNDPRAG.SQELGWCQRYFPEQDTTMSLSDP
Por_KAH8844570.1 LQARHGADARVIFTQNVPTFNDPRAG.SQELGWCQRYFPEQDTTMSLSDP
Por_KAI6541088.1 LQARHGADARVIFTQNVPTFNDPRAG.SQELGWCQRYFPEQDTTMSLSDP
Por_KAI6445464.1 LQARHGADARVIFTQNVPTFNDPRAG.SQELGWCQRYFPEQDTTMSLSDP

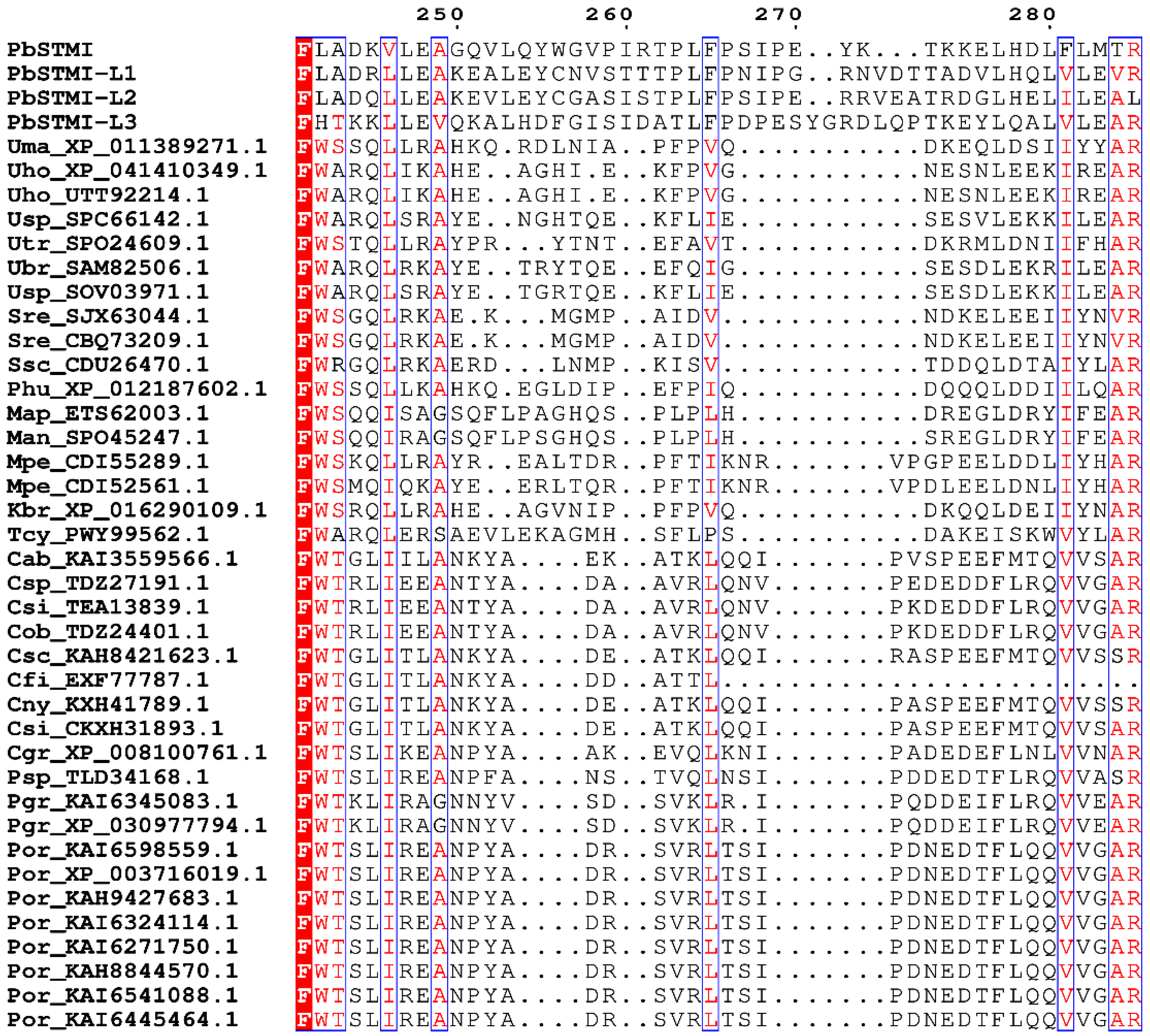

250 260 270 280
PbSTMI FLADKVLEAGQVLQYWGVPIRTPLFPSIPE..YK...TKKELHDLFLMTR
PbSTMI-L1 FLADRLLEAKEALEYCNVSTTTPLFPNIPG..RNVDTTADVLHQLVLEVR
PbSTMI-L2 FLADQLLEAKEVLEYCGASISTPLFPSIPE..RRVEATRDGLHELILEAL
PbSTMI-L3 FHTKKLLEVQKALHDFGISIDATLFPDPESYGRDLQPTKEYLQALVLEAR
Uma_XP_011389271.1 FWSSQLLRAHKQ.RDLNIA..PFPVQ..........DKEQLDSIIYYAR
Uho_XP_041410349.1 FWARQLIKAHE..AGHI.E..KFPVG..........NESNLEEKIREAR
Uho_UTT92214.1 FWARQLIKAHE..AGHI.E..KFPVG..........NESNLEEKIREAR
Usp_SPC66142.1 FWARQLSRAYE..NGHTQE..KFLIE..........SESVLEKKILEAR
Utr_SPO24609.1 FWSTQLLRAYPR...YTNT..EFAVT..........DKRMLDNIIFHAR
Ubr_SAM82506.1 FWARQLRKAYE..TRYTQE..EFQIG..........SESDLEKRILEAR
Usp_SOV03971.1 FWARQLSRAYE..TGRTQE..KFLIE..........SESDLEKKILEAR
Sre_SJX63044.1 FWSGQLRKAE.K...MGMP..AIDV...........NDKELEEIIYNVR
Sre_CBQ73209.1 FWSGQLRKAE.K...MGMP..AIDV...........NDKELEEIIYNVR
Ssc_CDU26470.1 FWRGQLRKAERD...LNMP..KISV...........TDDQLDTAIYLAR
Phu_XP_012187602.1 FWSSQLLKAHKQ.EGLDIP..EFPIQ..........DQQQLDDIILQAR
Map_ETS62003.1 FWSQQISAGSQFLPAGHQS..PLPLH..........DREGLDRYIFEAR
Man_SPO45247.1 FWSQQIRAGSQFLPSGHQS..PLPLH..........SREGLDRYIFEAR
Mpe_CDI55289.1 FWSKQLLRAYR..EALTDR..PFTIKNR......VPGPEELDDLIYHAR
Mpe_CDI52561.1 FWSMQIQKAYE..ERLTQR..PFTIKNR......VPDLEELDNLIYHAR
Kbr_XP_016290109.1 FWSRQLLRAHE..AGVNIP..PFPVQ..........DKQQLDEIIYNAR
Tcy_PWY99562.1 FWARQLERSAEVLEKAGMH..SFLPS..........DAKEISKWVYLAR
Cab_KAI3559566.1 FWTGLIILANKYA....EK..ATKLQQI......PVSPEEFMTQVVSAR
Csp_TDZ27191.1 FWTRLIEEANTYA....DA..AVRLQNV......PEDEDDFLRQVVGAR
Csi_TEA13839.1 FWTRLIEEANTYA....DA..AVRLQNV......PKDEDDFLRQVVGAR
Cob_TDZ24401.1 FWTRLIEEANTYA....DA..AVRLQNV......PKDEDDFLRQVVGAR
Csc_KAH8421623.1 FWTGLITLANKYA....DE..ATKLQQI......RASPEEFMTQVVSSR
Cfi_EXF77787.1 FWTGLITLANKYA....DD..ATTL.........................
Cny_KXH41789.1 FWTGLITLANKYA....DE..ATKLQQI......PASPEEFMTQVVSSR
Csi_CKXH31893.1 FWTGLITLANKYA....DE..ATKLQQI......PASPEEFMTQVVSAR
Cgr_XP_008100761.1 FWTSLIKEANPYA....AK..EVQLKNI......PADEDEFLNLVVNAR
Psp_TLD34168.1 FWTSLIREANPFA....NS..TVQLNSI......PDDEDTFLRQVVASR
Pgr_KAI6345083.1 FWTKLIRAGNNYV....SD..SVKLR.I......PQDDEIFLRQVVEAR
Pgr_XP_030977794.1 FWTKLIRAGNNYV....SD..SVKLR.I......PQDDEIFLRQVVEAR
Por_KAI6598559.1 FWTSLIREANPYA....DR..SVRLTSI......PDNEDTFLQQVVGAR
Por_XP_003716019.1 FWTSLIREANPYA....DR..SVRLTSI......PDNEDTFLQQVVGAR
Por_KAH9427683.1 FWTSLIREANPYA....DR..SVRLTSI......PDNEDTFLQQVVGAR
Por_KAI6324114.1 FWTSLIREANPYA....DR..SVRLTSI......PDNEDTFLQQVVGAR
Por_KAI6271750.1 FWTSLIREANPYA....DR..SVRLTSI......PDNEDTFLQQVVGAR
Por_KAH8844570.1 FWTSLIREANPYA....DR..SVRLTSI......PDNEDTFLQQVVGAR
Por_KAI6541088.1 FWTSLIREANPYA....DR..SVRLTSI......PDNEDTFLQQVVGAR
Por_KAI6445464.1 FWTSLIREANPYA....DR..SVRLTSI......PDNEDTFLQQVVGAR

```
                                 290       300       310            320
                                   .         .         .              .
PbSTMI               KNDPSTTGSGLREHFRAYLEEAIPKLDE......LSYWKRLDTNLLRRLR
PbSTMI-L1            KNENKSMSRELRQHFRTYLDEALPELNRCMHAGSSVVWQRCDNTLVKRLE
PbSTMI-L2            KNERNSMGRQLRQYFRTYLEEAIRELKRCMQEGHSRVWQRGGTNLVKLLE
PbSTMI-L3            KNEPGTQGQILRQHFINYIDESVKALERV.........EGCDTKMLLRVK
Uma_XP_011389271.1   TNPEHPNSQLWRAAIRDYIWTALANN.PGK.........EKDNLVLTRLL
Uho_XP_041410349.1   MSPER..NEQFRASVRDYITQVLGEYADKD.........VESKTLLKRLK
Uho_UTT92214.1       MSPER..NEQFRASVRDYITQVLGEYADKD.........VESKTLLKRLK
Usp_SPC66142.1       MRPEQ..NEQFRASVREYISRVLNLYKPEE.........VESDRLLKRLK
Utr_SPO24609.1       KYPNG.KWKLWRDQIAEYVKGVLDEH.PDP.........RHDDVLLSRLR
Ubr_SAM82506.1       VHPEQ..NEQFRANVRDYIRQVLDQYTPDQ.........VERDVLLKRLK
Usp_SOV03971.1       MRPEQ..NEQFRVSVGEYISRVLNHYEPEQ.........VESDLLLKRLK
Sre_SJX63044.1       MNPQG..NVAWRNHLAWYVQKTLDQN.ANQ.........EATSKTLGRYR
Sre_CBQ73209.1       MNPQT..NIAWRDYLAQYVQETLDRN.ADQ.........EATSKTLGRYR
Ssc_CDU26470.1       MEPAQ..NANWRNYLAEYVQETLNQN.SHM.........EGTSKALDRYR
Phu_XP_012187602.1   THPNHPNSAAWRNYITQYINTALANN.PGR.........DTDNLVLTRLR
Map_ETS62003.1       RDPTR..YAARRHAIYHHVDAILNSMSAEQK.....QERA...RLVGRLE
Man_SPO45247.1       REPAR..YAARRQAIYHHVNAILNSMSAEQK.....EQKA...RLVGRLE
Mpe_CDI55289.1       VNPESIQGKKWRKSITKYVQEVLHRN.SDN.........PNKSIVLRRFR
Mpe_CDI52561.1       VNPMTVRGREGRKSIASFINKVLKQH.PET.........ETESLVLKRFR
Kbr_XP_016290109.1   VHPEYPTSAQWRRYMTDYIQSALRAN.PNA...........DHMTLTRLK
Tcy_PWY99562.1       KDPAQ..YLYYREQIVKQVETTLQVIAQLPE.....QERAKYGALHFRLQ
Cab_KAI3559566.1     LK.ET....TLRVQILKMLESAVSNQQFAQD..........SPRSYSRVR
Csp_TDZ27191.1       LE.DG....PLRKQILAMLQSAAGSETFKKE..........SSRSHDRVS
Csi_TEA13839.1       LE.DG....PLRKQILAMLESAARSETFKKE..........SSRSHGRVS
Cob_TDZ24401.1       LE.DG....PLRKQILAMLESAARSETFKKE..........SSRSHGRVS
Csc_KAH8421623.1     LE.ET....TLRVQILKMLKSAISNQQFAQD..........SPRSYSRVR
Cfi_EXF77787.1       ..................................................
Cny_KXH41789.1       LK.ET....TLRVQILKMLKSAVSNQQFAQD..........SPRSYSRVR
Csi_CKXH31893.1      LK.ET....TLRFQILNMLKSAVSNPQFAQD..........SPRSYSRVR
Cgr_XP_008100761.1   LD.DN....NFRKSVHAMLESAARSETFKEH..........SQRSHYRVA
Psp_TLD34168.1       LKDEN....TLRKPIYAMVESAAQNPAFKRK....SPRLNESPRAQDRVK
Pgr_KAI6345083.1     QQ.DN....ALRRQIYAMLTQAGQSQQFVSA..........DKRSWARMV
Pgr_XP_030977794.1   QQ.DN....ALRRQIYAMLTQAGQSQQFVSA..........DKRSWARMV
Por_KAI6598559.1     MH.QN....SLRGKIYAMLYSAAQSSQFASN..........NPRSQHRVS
Por_XP_003716019.1   MY.QN....SLRGKIYAMLYSAAQSSQFASN..........NPRSQHRVS
Por_KAH9427683.1     MH.QN....SLRGKIYAMLYSAAQSSQFASN..........NPRSQHRVA
Por_KAI6324114.1     MH.QN....SLRGKIYAMLYSAAQSSQFASN..........NPRSQHRVS
Por_KAI6271750.1     MH.QN....SLRGKIYAMLYSAAQSSQFASN..........NPRSQHRVS
Por_KAH8844570.1     MY.QN....SLRGKIYAMLYSAAQSSQFASN..........NPRSQHRVS
Por_KAI6541088.1     MY.QN....SLRGKIYAMLYSAAQSSQFASN..........NPRSQHRVS
Por_KAI6445464.1     MY.QN....SLRGKIYAMLYSAAQSSQFASN..........NPRSQHRVS
```

```
                      330       340       350       360                 370
                        .         .         .         .                   .
PbSTMI                QDVRPVFDDDAEMQILISDGNQVAAVIDAVYQ........NSGTLKNMRL
PbSTMI-L1             EHVLPVFQDDAQMQILISDGNQVAAVIDAKHQ........NWTNLKGMEF
PbSTMI-L2             QNFLPAFQDDAEMQILMGNSNQVAAVIDAKET........DWANLTGMQI
PbSTMI-L3             THVLGAFRKPLS..VELCDANHIAAVVDIMEERNTISGMCKANNMRGMKF
Uma_XP_011389271.1    HTHLPEFTGD..ATLELADAAHIAAFHRYQAG......G..SGMQAEPI
Uho_XP_041410349.1    YTLLPEFTAS..TTLELADANHVAAFHRYMDNLQRHG.NGHDSGFHTVPV
Uho_UTT92214.1        YTLLPEFTAS..TTLELADANHVAAFHRYMDNLQRHG.NGHDSGFHTVPV
Usp_SPC66142.1        YTLLPEFTKA..TTLELADANHVAAFHRYLDSK.QHV.NGHDSGLYTIPV
Utr_SPO24609.1        HTLLPEFTGP..STLELADANHVAAFHRHLDG......D..TNMHWKPV
Ubr_SAM82506.1        YTLLPEFTEA..TTLELADAIHVAAFHRYLDSMQQHG.NGHGSGLYTVPV
Usp_SOV03971.1        YTLLPEFTEA..TTLELADAIHVAAFHRYLDSK.QHA.NGHDYGLHTIPV
Sre_SJX63044.1        NTLLPEFTGNNPVTLELADAGHIAALHRYWDGLR....DHKATGMQLVPI
Sre_CBQ73209.1        NTLLPEFTGNNPVTLELADAGHIAALHRYWDGLR....DRKATGMQLVPI
Ssc_CDU26470.1        HTLLPEFTGENPVTVELADAGHIAALHRYQDSLK....RGPQTPMWMQYI
Phu_XP_012187602.1    HTHLPEFSGD..ATLELADAAHIAAFHRFQDG......G..SNMHAEPV
Map_ETS62003.1        NTFKPEFGGA.GTTIEMADAAHIAHFHRAQAAN.........GA.GGVWA
Man_SPO45247.1        NTFKPEFGGA.RTTIEMADAAHIAHFHRAQAAD.........GA.GGVWA
Mpe_CDI55289.1        QTFLPEFTQA..PTLEVADANHMAAFHRNLDVR...N.TGHVSDMYWDRV
Mpe_CDI52561.1        YTFLPEFAGE..PTLELADANHVAAFHRNLDDW...H.NRGVSDMYWDRV
Kbr_XP_016290109.1    YTHLPEFSGA..PTLELADASHVAAFHRYLDE......GSRSGMHVHPV
Tcy_PWY99562.1        NTLLPEFS.E.GTTIELADAGHVALFHRAMAGD.........RRIERVPV
Cab_KAI3559566.1      DILVEEFTGKPAPTLELGDANHMAIVLQYLDNETKDH..NSY.AMRLLPA
Csp_TDZ27191.1        NILVNEFTGTPSPTLELGDANHITAVLEYLDGEAAKS..GGA.AGKLLPA
Csi_TEA13839.1        NILVNEFTGTPSPTLELGDANHITAVLEYLDGEAAKS..GDA.AGKLLPA
Cob_TDZ24401.1        NILVNEFTGTPSPTLELGDANHITAVLKYLDGEAAKS..GGA.AGKLLPA
Csc_KAH8421623.1      DILVEEFTGNPAPTLELGDANHMAIVLQYLDNETKDR..NSY.AMRLLPA
Cfi_EXF77787.1        ..................................................
Cny_KXH41789.1        DILVDEFTGKPAPTLELGDANHMAIVLQYLDNETKDR..NSY.AMRLLPA
Csi_CKXH31893.1       DILVEEFTGKPAPTLELGDANHMAIVLQYLDNETKDR..NSY.AMRLLPA
Cgr_XP_008100761.1    NILVGEFAGEPAPTLELGDANHITAVLEYLDTEMRGA.RGRS.AGKLQPA
Psp_TLD34168.1        QILVPEFCKEPSPTLEMGDANHITAVLNYLDQEKSGG...RG.AGRLVHA
Pgr_KAI6345083.1      KILIPEFEGAPKPTLELGDANHITAIFNYLDQEAGY...VSG.AGQLQYA
Pgr_XP_030977794.1    KILIPEFEGAPKPTLELGDANHITAIFNYLDQEAGY...VSG.AGQLQYA
Por_KAI6598559.1      KILVPEFQGSPDPTLELGDANHITAVLGYLDQEQYGG...SG.AGRLVPA
Por_XP_003716019.1    KILVPEFQGSPDPTLELGDANHITAVLGYLDQEQYGG...SG.AGRLVPA
Por_KAH9427683.1      KILVPEFQGSPDPTLELGDANHITAVLGYLDQEQYGG...SG.AGRLVPA
Por_KAI6324114.1      KILVPEFQGSPDPTLELGDANHITAVLGYLDQEQYGG...SG.AGRLVPA
Por_KAI6271750.1      KILVPEFQGSPDPTLELGDANHITAVLGYLDQEQYGG...SG.AGRLVPA
Por_KAH8844570.1      KILVPEFQGSPDPTLELGDANHITAVLGYLDQEQYGG...SG.AGRLVPA
Por_KAI6541088.1      KILVPEFQGSPDPTLELGDANHITAVLGYLDQEQYGG...SG.AGRLVPA
Por_KAI6445464.1      KILVPEFQGSPDPTLELGDANHITAVLGYLDQEQYGG...SG.AGRLVPA
```

```
                                  380          390         400         410
PbSTMI               VGNDRDM.......VYELAPSEE..DTDDKMIYGASRERCLDLLRPTLQ.
PbSTMI-L1            VYNHKDK.......VYEFAEAPG...AADKMLCDASPTRCLELLRPTLLS
PbSTMI-L2            VYNVEDM.......VFELSPAAH..VADDKMIYDASPTRCLDLLRSTLLP
PbSTMI-L3            VYNDGDG.......FFQFAPDDE...AGDQMLYSLNAARCLALLATKWPK
Uma_XP_011389271.1   EFSAHAP.AEGEKVRFFPAT..S..ELHGYLLRGANRDQDLAYIKYLAGV
Uho_XP_041410349.1   YYPDGDH.PENAAVPFQPTTG.E..KFHGVLLKGANRDLDLGYIKQLASL
Uho_UTT92214.1       YYPDGDH.PENAAVPFQPTTG.E..KFHGVLLKGANRDLDLGYIKQLASL
Usp_SPC66142.1       HYPDGNH.DEDTVVPFQSATE.T..ELHGFLLKGAHRDLDLGYIKRLAGL
Utr_SPO24609.1       AYPVGPH.EAGQKVSFEDATN.Q..ELHGWMLKGADRDADLLHIKRLAGW
Ubr_SAM82506.1       HYPDGNH.LETTVVPFQSATG.K..ELHGLLLKGAHRDLDLGYIKRLAGL
Usp_SOV03971.1       HYPDGNH.HENTVVPFQSATG.K..ELHGVLLKGANRDLDLSYIKRLAGL
Sre_SJX63044.1       KFLDGSD.NPDDKIDMVRSNQ.N..DNHGMLLMGADRDKDLRYIKQLAGM
Sre_CBQ73209.1       KFLDGSD.NPDDKIDMARSNR.A..DNHGMLLVGADRDKDLRYIKQLAGM
Ssc_CDU26470.1       KFDDGSN.DPGEKIKMLQGNQ.W..DHHGYLLQGANRDEDLKYIERLAKI
Phu_XP_012187602.1   RFAAGPH.AEGAKVDFSPATG.N..ELHGYLVRGADRDEDLAYIKQLAGV
Map_ETS62003.1       RHERGVT.APGAAAGFQQVTHPA..HADGVLLQGGHTALDRAWIDGLVRS
Man_SPO45247.1       RYERGVT.APGSAAGFQEVAHPA..HADGVLLQGGHTAMDRAWIDQLVRN
Mpe_CDI55289.1       RYPDGPH.DRNAIVHFEEATSDG..DTHGMLLRGADRNKDLQYIKNLAGL
Mpe_CDI52561.1       KYPDGPH.DIKTIPPFKEATSNS..DAHGWLLRGADRDEDLLHIKRLAGL
Kbr_XP_016290109.1   AYPPGPH.EAGKRVGFVSAAG.N..ELHGVLLKGADRDQDLEYIKRLAGI
Tcy_PWY99562.1       YYPSGHY.EPGAAVGFKKALDPQ..QAHGMLLQGASTELDREWIEALFHS
Cab_KAI3559566.1     LCDTSEE.NPTQP.PRVSAGAPAAQSPDGWVLAGCDIQDTRGYIEGLFQE
Csp_TDZ27191.1       VCDKTEQ.NPTLP.PKVDTGGPTA.ATEGWVLTGCDIKQTRADIERLFG.
Csi_TEA13839.1       VCDKTEQ.NPTLP.PKVDTGGPTA.ATEGWVLTGCDIKQTRADIERLFG.
Cob_TDZ24401.1       VCDKTEQ.NPTLP.PKVDTGGPTA.ATEGWVLTGCDIKQTRADIERLFG.
Csc_KAH8421623.1     LCDTSEE.NPTQP.PRVSAGAPAAQSPDGWVLAGCNIHDTRRYIEGLFQE
Cfi_EXF77787.1       ..................................................
Cny_KXH41789.1       LCDTSEE.NPTQP.PRVSAGAPAAQSPDGWVLAGCNIHDTRRYIEGLFQE
Csi_CKXH31893.1      LCDTSEE.NPTQP.PRVSAGAPAAQSPDGWVLAGCSIHDTRKYIEGLFQE
Cgr_XP_008100761.1   SCDKSGK.DPRQP.PNVIIGTRP..SNSGWVLTGCDVTQTRNNIERLLQQ
Psp_TLD34168.1       VCDTRGT.DPTQP.PVVKMGTAP..RGEGWVLTGCHIQQTRNGIESILRE
Pgr_KAI6345083.1     VCDQSNTNNPSQP.PDVRAGSSW..TGHGFILTGCDIDRVRDDIARMFH.
Pgr_XP_030977794.1   VCDQSNTNNPSQP.PDVRAGSSW..TGHGFILTGCDIDRVRDDIARMFH.
Por_KAI6598559.1     ACDTSGT.DPSQP.PAVTTTSRS..RGEGWVLTNCDIRQIRNKIENILRR
Por_XP_003716019.1   ACDTSGT.DPSQP.PAVTTTSRS..RGEGWVLTNCDIRQIRNKIENILRR
Por_KAH9427683.1     ACDTSGT.DPSQP.PAVTTTSRS..RGEGWVLTNCDIRQIRNKIENILRR
Por_KAI6324114.1     ACDTSGT.DPSQP.PAVTTTSRS..RGEGWVLTNCDIRQIRNKIENILRR
Por_KAI6271750.1     ACDTSGT.DPSQP.PAVTTTSRS..RGEGWVLTNCDIRQIRNKIEDILRR
Por_KAH8844570.1     ACDTSGT.DPSQP.PAVTTTSRS..RGEGWVLTNCDIRQIRNKIEDILRR
Por_KAI6541088.1     ACDTSGT.DPSQP.PAVTTTSRS..RGEGWVLTNCDIRQIRNKIENILRR
Por_KAI6445464.1     ACDTSGT.DPSQP.PAVTTTSRS..RGEGWVLTNCDIRQIRNKIEDILRR
```

| | |
|---|---|
| **PbSTMI** | ....... |
| **PbSTMI-L1** | Q...... |
| **PbSTMI-L2** | Q...... |
| **PbSTMI-L3** | ....... |
| **Uma_XP_011389271.1** | HF..... |
| **Uho_XP_041410349.1** | TH..... |
| **Uho_UTT92214.1** | TH..... |
| **Usp_SPC66142.1** | PH..... |
| **Utr_SPO24609.1** | LH..... |
| **Ubr_SAM82506.1** | PH..... |
| **Usp_SOV03971.1** | PH..... |
| **Sre_SJX63044.1** | R...... |
| **Sre_CBQ73209.1** | R...... |
| **Ssc_CDU26470.1** | R...... |
| **Phu_XP_012187602.1** | WH..... |
| **Map_ETS62003.1** | VRG.... |
| **Man_SPO45247.1** | VRG.... |
| **Mpe_CDI55289.1** | LR..... |
| **Mpe_CDI52561.1** | FR..... |
| **Kbr_XP_016290109.1** | PH..... |
| **Tcy_PWY99562.1** | R...... |
| **Cab_KAI3559566.1** | NALP... |
| **Csp_TDZ27191.1** | ....... |
| **Csi_TEA13839.1** | ....... |
| **Cob_TDZ24401.1** | ....... |
| **Csc_KAH8421623.1** | NALP... |
| **Cfi_EXF77787.1** | ....... |
| **Cny_KXH41789.1** | NALS... |
| **Csi_CKXH31893.1** | NALP... |
| **Cgr_XP_008100761.1** | R...... |
| **Psp_TLD34168.1** | SERSRVF |
| **Pgr_KAI6345083.1** | ....... |
| **Pgr_XP_030977794.1** | ....... |
| **Por_KAI6598559.1** | SA..... |
| **Por_XP_003716019.1** | SA..... |
| **Por_KAH9427683.1** | SA..... |
| **Por_KAI6324114.1** | SA..... |
| **Por_KAI6271750.1** | SA..... |
| **Por_KAH8844570.1** | SA..... |
| **Por_KAI6541088.1** | SA..... |
| **Por_KAI6445464.1** | SA..... |

**Other Supplementary information for this manuscript includes the following (Separate attachments):**

**Table S1:** A list of $^{\Delta sp}$PbSTMI orthologs identified after BlastP searches against the NCBI non-redundant (nr) database.

**Table S2:** $^{\Delta sp}$PbSTMI paralogs with accession numbers, genomic locations, protein sequences, and identity.

**Table S3.** List of primers used in this study.

**Table S4:** List of antibodies used in this study.

**Movies S1:** Z-stack images showing $^{\Delta sp}$PbSTMI-GFP and AtEDS1-mRFP colocalization in *N. benthamiana* leaves.

**Supplementary Dataset 1:** Summary of statistical analyses.
